## Supplementary Note for "Tracing animal genomic evolution with the chromosomal-level assembly of the freshwater sponge *Ephydatia muelleri*"

Kenny et al.

### Contents

**Supplementary Note 1:** *Ephydatia muelleri:* an introduction 3
1.1 Description and distribution 3
1.2 Position in demosponges 4
1.3 Place in metazoan phylogeny 6
**Supplementary Note 2:** Sequencing and Assembly 8
2.1 Animal source and DNA isolation 8
2.2 DNA cleanup and quality verification 8
2.3 Initial Assembly 8
2.4 Scaffolding 8
**Supplementary Note 3:** Assembly comparisons 11
3.1 Assembly metrics and comparisons 11
3.2 BUSCO results 14
3.3 Repeat content 16
3.4 Contiguity and scaffolding 19
3.5 Checks for contamination 20
3.6 Synteny analyses 22
**Supplementary Note 4:** Analysis of scaffold 25*: Flavobacterium* sp. Genome 25
4.1 Taxonomy assignment 25
4.2 Genomic Islands 31
4.3 Secondary metabolites 32
**Supplementary Note 5:** Gene models and annotation 35
5.1 Gene prediction and metrics 35
5.2 Automated annotation results 38
5.3 Functional annotation 38
**Supplementary Note 6:** Genome Architecture and Insights into Regulatory Structures 40
6.1 Insights into longer-range gene regulation and chromatin interaction 40

6.2 Cytosine DNA methylation 42
**Supplementary Note 7:** Analysis of *Ephydatia muelleri* novelties 45
7.1 Gains and losses 45
7.2 Gene overlap and novelty in freshwater sponges 49
7.3 Positive selection 51
7.4 Comparison with other freshwater lineages 58
7.5 Cluster expansion and novel *Ephydatia muelleri* genes 61
**Supplementary Note 8:** Developmental RNAseq in *Ephydatia muelleri* 74
8.1 RNAseq Methods 74
8.2 Expanded results of RNAseq 75
**Supplementary Note 9:** Gene content in *Ephydatia muelleri* 82
9.1 Ion channels 82
9.2 Epithelia 86
9.3 Wnt pathway 86
9.4 Comparative gene expression during development: 87
**Supplementary Note 10:** Amplicon analysis of holobiont content 96
10.1 Microbial community structure methods 96
10.2 Results for microbial community structure within *Ephydatia muelleri* 98
**Supplementary Note 11:** *Ephydatia* as a research model 105
**Supplementary References:** 110

### Supplementary Note 1: *Ephydatia muelleri:* an introduction

#### 1.1 Description and distribution

*E. muelleri* can occur in shades of green, yellow, or brown, but when well-lit it usually appears green, with the colour coming from the algal symbionts within its body. In different levels of shading, it becomes more yellow and lacks photosynthetic symbionts. The surface is undulating and has oscula which are not significantly raised, but with delicate translucent oscula membranes. *E. muelleri* usually inhabits lakes and streams where there a reasonable level of water flow, and ambient pH is usually greater than 5.9 and less than 9.1[1] Gemmules are yellow and range in size from 200-400 micrometers. Gemmules are distributed throughout the skeleton, but if collected in the summer, the base attached to the substrate is more likely to include gemmules.

The species can be most readily identified by the gemmule spicule type known as "birotules" as described[1]: rotules flat, umbonate, deeply and irregularly incised, forming no more than 12 long rays; shaft normally smooth, rarely with 1 or 2 spines; gemmosclere length never greater than rotule diameter. The spicules of the adult sponge are megascleres described as "stout or slender amphioxeae, usually densely covered with short conical spines, except near the tips ... in rare cases, megascleres are entirely smooth; both smooth and variably spined forms are often present in the same specimen; megasclere length 17 1 - (245) -3 1 1 pm (SDZS0 = 26.4), width 5 - ( 1 1 ) -23 pm (SD = 3.7). Microscleres absent" [1].

Records of *Ephydatia muelleri:*

Freshwater sponges are in the order Spongillida, which comprises about 3% of sponge taxa [2]. The species *E. muelleri* has a temperate distribution and is reported from all of central and northern Europe, a river near Irkutsk (southern lake Baikal), Japan, throughout North America (from west, mid, and eastern USA and Canada) as well as Iceland (Supplementary Figure 1A, Supplementary Table 1).

The genus *Ephydatia* is more widespread, and found on all continents except Antarctica[3]. Fossil records show the genus *Ephydatia* as widely distributed since the middle Eocene (approx. 40 Mya): *Ephydatia* cf. *facunda* Giraffe Formation, Canada, Middle Eocene; *Ephydatia gutenbergiana,* Central Europe, Middle Eocene; *Ephydatia fossilis*- Central Europe, Miocene; *Ephydatia chileana* - Chile, Late Miocene.

**Supplementary Table 1:** Countries in which *Ephydatia muelleri* has been recorded. Example references are given.

| Country | References | Location Detail |
|---|---|---|
| **Ireland** | Stephens, J. (1920). The freshwater sponges of Ireland. Proc Roy Irish Acad 35 (B): 205-254 | |
| | Lucey & Cocciglia (2014) Biology & Environment: Proc Roy Irish Acad 114(2):89-100 | |
| **England** | Bowerbank, J.S. (1874). A Monograph of the British Spongiadae. Volume 3. (Ray Society: London): i-xvii, 1-367, pls I-XCII. | Exe River Cornwall |
| | Evans, K. (2019). Invertebrate Biology, DOI: 10.1111/ivb.12258 | |
| **France** | Topsent, E. (1893). Note sur la faune des Spongillides de France. Bulletin de la Société Zoologique de France. 18: 176 | 15 km W of Châteaudun, on Yerre River (La Filandière) |
| **Czech Republic** | Opravilova (2006) Bohemia centralis, Praha 27:19-22 | |
| **Croatia** | Imešek et al. (2013) Organisms Diversity and Evolution 13(2):127-134 | polje Jezero and Peračko blato |
| **Belgium** | Richelle et al (1995) Arch Hydrobiol 135(2) 209-231 | |
| **Netherlands** | Van Soest (1977) Zoologische Mededelingen 50(16): 261-273 | |
| **Estonia** | Roovere et al (2006) Proc Es Acad Sci Biol Ecology 55(2)216-227 | |
| **Poland** | Czeczuga et al (2015) Afr. J. Biotechnology 14(45):3093-3100 | |
| **Switzerland** | Manconi and Desqueyroux-Faúdez, R (1999) Revue suisse de zoologie 106:571-580 | |
| **Austria** | Drosher & Waringer (2007) Freshwater Biology 52: 998–1008 | |
| **Germany** | Imsieke et al (1996) Env. Toxicol & Chem 15(8)1329-1334 | Bonn River Sieg, Opsen |
| | Vohmann et al (2009) Freshwater Biol 54:1078-1092 | Rhine |
| **Denmark** | Tendal OS, 1967a. On the freshwater sponges of Denmark. Videns. Meddel. Dansk Naturhist. For. 130:173-8. | |
| **Iceland** | Tendal OS, 1976. Freshwater spongia. Zoology Iceland 2(4a):1-4 | |
| **Russia** | Kalyuzhnaya et al (2011) Mol Biol 45(4) 567-575 | Goloustnaya river Baikal (near Irkutsk) |
| **Norway** | Økland & Økland (1996) Hydrobiologia 330: 1-30 | All Norway (survey) |
| **Japan** | Mukai (1992) J. Exp. Zool. 264:298-311 | Tataranuma, a pond near Tatebayashi, Gunma Prefecture |
| | Ishijima et al. (2008) Zool. Sci. 25:480-486 | Kamiike (KAM) in the Okayama Prefectural Nature Conservation Center |
| **Canada** | Reiswig & Miller (1998) Invertebrate Biology 117(1):1-8 | Ste. Anne-de-Bellevue, Quebec |
| | Ricciardi & Reiswig (1993) Can J. Zool 71:665-682 | Nova Scotia, Quebec, Ontario, New Brunswick, Newfoundland |
| | Elliott & Leys (2007) J. Exp. Biol. 210: 3736-3748 | Vancouver Island, British Columbia |
| | This paper | Sooke Reservoir and Head Tank, Victoria, British Columbia |
| **USA** | Elvin (1971) Trans Am Micr. Soc. 90(2):219-224 | Mill River, New Haven, Connecticut |
| | Rivera et al. (2011) BMC Biotechnology 11:67 | Salmon Lake, Montana |
| | Sowka, P.A.(1999) Southwestern Naturalist 44(2):211-212. | Arizona |

### 1.2 Position in demosponges

*Ephydatia muelleri* is a demosponge and a member of the Heteroscleromorpha[4]. Within these clades, it is a member of the Spongillidae, in the order Spongillida. This order has a sister-group relationship with Sphaerocladina (Supplementary Figure 1B), which is strongly supported by *18S* [5,6], *CO1* and *28S* [7] and mitogenomic evidence[8]. Under the birth-death clock model, the divergence time of the last common ancestor of extant Spongillida is placed at 18 My, and the divergence of Sphaerocladina from the Spongillida at 311 million years before the present day.

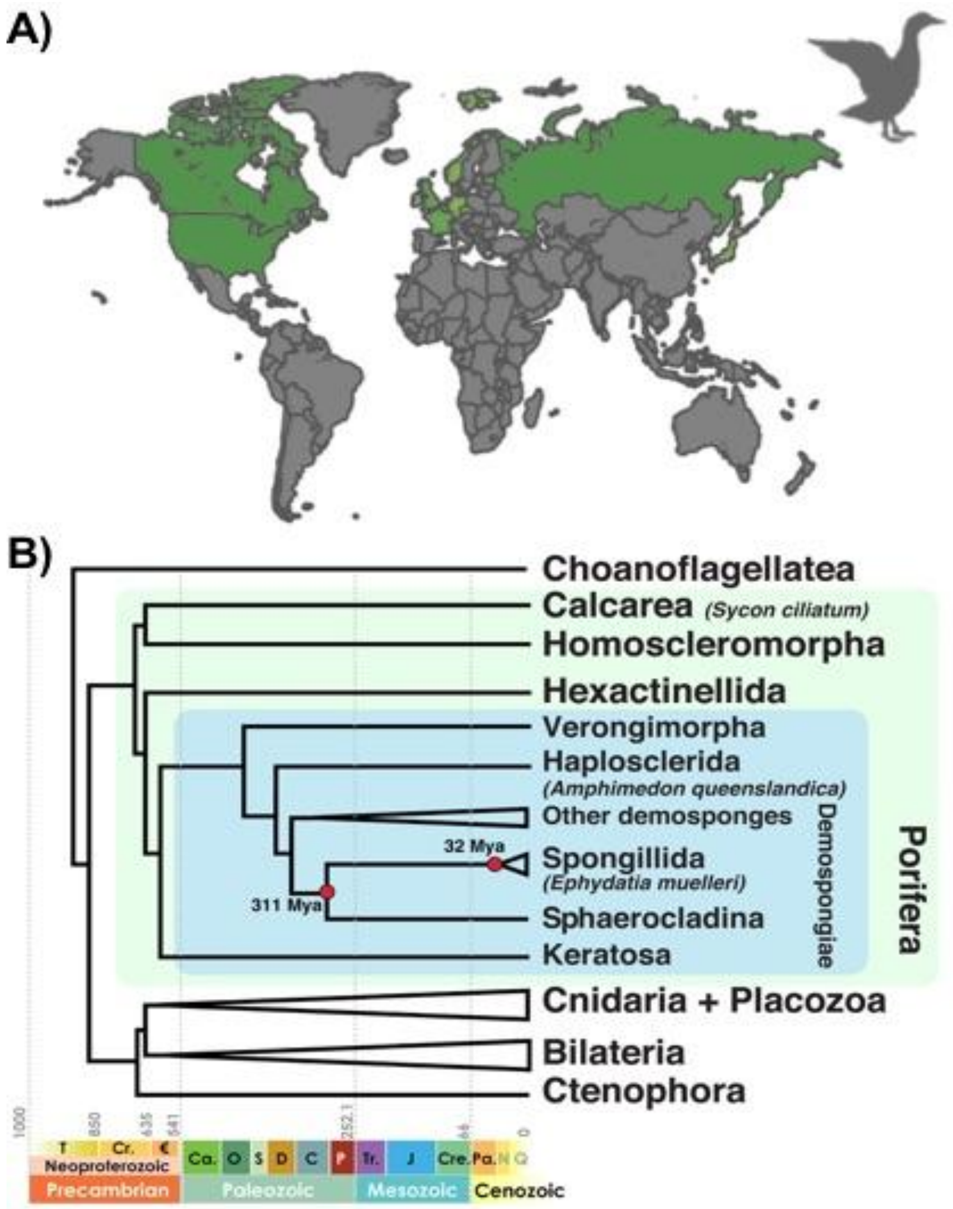

**Supplementary Figure 1:** A) Known distribution of *Ephydatia muelleri*. Other locations are almost certainly inhabited by this species, but only referenced locations (see Supplementary Table 1) marked in green. The gemmules are transferred on the feathers of waterfowl. B) Time tree modified from [8,9], based on parameters of BEAST analysis 2 and Bayesian inference analysis plotted on stratigraphic chart. Please note that the relative phylogenetic positions of outgroup taxa are the source of some debate (Supplementary Note 1.3).

#### 1.3 Place in metazoan phylogeny

The order of diversification of extant non-bilaterian phyla from the metazoan stem lineage is the source of much debate. Some studies place ctenophores as the sister taxa to all other metazoans [10,] while others place sponges in this position [11], and others suggest other potential arrangements [12]. The answer to this question is presently unclear [13]. This paper does not set out to resolve this question, but may provide extra data for future analyses. However, it is important for the discussion in this manuscript to note that sponges are one of the earliest phyla to emerge from the metazoan stem lineage and as such represent a key outgroup to other metazoan species for the consideration of traits such as the evolution of multicellularity, cell signalling apparatus, and the origin of genes and genetic pathways involved in the acquisition of neural systems, muscles and epithelia.

Supplementary Figure 2 shows one hypothesis for the phylogenetic relationships of sponges to a number of other metazoan and choanozoan species. In this phylogeny, Porifera are the sister taxa to all other metazoans. Even if the position of Porifera and Ctenophora were reversed, the utility of the *Ephydatia muelleri* genome for understanding the origin of metazoan traits is clear from the arrangement of the tree.

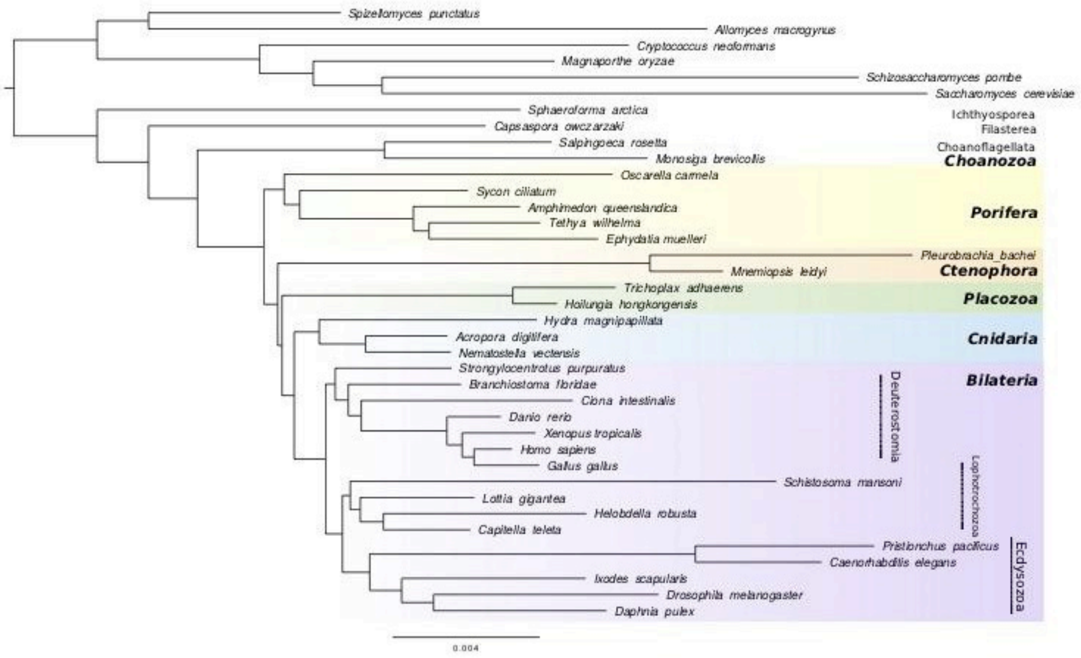

**Supplementary Figure 2***:* Phylogenetic tree reconstruction from a Bayesian analysis of orthologous gene families, including 38 species from various phyla, including several non-metazoan outgroups. All nodes had a posterior probability indistinguishable from one except where otherwise noted. Note: in most recent phylogenetic reconstructions, Homoscleromorpha and Calcarea (here, *Oscarella carmella* + *Sycon ciliatum*) form a monophyletic clade, sister to the Silicea (Hexactinellida and Demospongiae). Both *Tethya wilhelma* and *Ephydatia muelleri* are demosponges.

*Methods: Phylogenetic reconstruction:*

A dataset of 36 species was used from a previous study by Pett et al. [14], to which *Tethya wilhelma* was added together with the newly sequenced *E. muelleri* predicted proteome. The dataset included six Fungi, four Choanozoa, five Porifera, two Ctenophora and two Placozoa, three Cnidaria and 16 Bilateria. Proteome data obtained by whole genome prediction was used for all the species.

The phylogeny was generated using 23,365 orthologous gene families using RevBayes version 1.0.11 [15] following the steps described in Pett et al. [14]. The orthologous gene families were predicted using OrthoFinder 2[16]. Convergence statistics of the Bayesian analysis were observed using Tracer version 1.71 [17] and computed for the four independent Markov chains for 50,000 generations with bpcomp and tracecomp in the PhyloBayes package version 4.1 [18], resulting in maxdif $< 0.1$ and effsize $> 300$ with burn in of 1000 chains (Supplementary Figure 2).

### Supplementary Note 2: Sequencing and Assembly

#### 2.1 Animal source and DNA isolation

Material was derived from a single clone of the sponge *Ephydatia muelleri* collected as overwintering cysts (gemmules) from the head tank of the Kapoor Tunnel (Sooke Reservoir), part of the drinking water system of the city of Victoria, British Columbia, Canada. Gemmules were held at 3°C for one month at the University of Alberta and then freed from the adult skeleton, cleaned in 1% hydrogen peroxide and cultured in sterile freshwater media [19]. Tissue from one week old sponges hatched from the single clone was flash frozen in liquid nitrogen and stored at -80ºC.

#### 2.2 DNA cleanup and quality verification

DNA was extracted using Genomic-tip 20G (Qiagen, Toronto, Canada) columns immediately prior to PacBio sequencing. DNA extraction to assembly steps were carried out by Dovetail Genomics (Scotts Valley, CA, USA) with slight modifications of the manufacturer's protocols. DNA quantity and quality was verified by Qubit and gel electrophoresis respectively.

#### 2.3 Initial Assembly

The *de novo* assembly was performed using the FALCON 1.8.8 pipeline from Pacific Biosciences. First, 58x fold whole-genome, single-molecule, real-time sequencing (SMRT) data from our *Ephydatia muelleri* sample was used as input to the traditional FALCON pipeline using a length cut-off that correspond to 50x coverage of data during the initial error-correcting stage. This resulted in 1.1 million error corrected reads with an N50 read length equal to 13.6kbp covering 36.7x of the 340Mb genome. Second, the error-corrected reads were processed by the overlap portion of the FALCON pipeline. The aligned reads were assembled in the third stage of FALCON into 3590 primary contigs containing 322 Mb with an N50 contig length of 219kbp. The assembly was verified as haploid by checking coverage in the course of assembly, although heterozygosity was not successfully sampled due to inadequate coverage. Finally, the assembly was polished through PacBio's Arrow algorithm from SMRT Link 5.0.1, using the original raw-reads.

#### 2.4 Scaffolding

***Chicago library preparation and sequencing:***

Two DNA libraries were prepared based on the Chicago method [20]. Briefly, for each library, ~500 ng of HMW gDNA (mean fragment length = 100kb) was reconstituted into chromatin in vitro and fixed with formaldehyde. Fixed chromatin was digested with DpnII, the 5' overhangs filled in with biotinylated nucleotides, and then free blunt ends were ligated. After ligation, crosslinks were reversed, and the DNA purified from protein. Purified DNA was treated to remove biotin that was not internal to ligated fragments. The DNA was then sheared to ~350 bp mean fragment size and sequencing libraries were generated using NEBNext Ultra enzymes and Illumina-compatible adapters. Biotin-containing fragments were isolated using streptavidin beads before PCR enrichment of each library. The libraries were sequenced on an Illumina HiSeqX. The number and length of read pairs produced for each library was: 63 million, 2x150bp for library 1; 226 million, 2x150bp for library 2. Together, these Chicago library reads provided 155.48x physical coverage of the genome (1-50kb pairs).

***Dovetail HiC library preparation and sequencing:***

Two Dovetail HiC libraries were prepared in a similar manner as described previously [21]. Briefly, for each library, chromatin was fixed in place with formaldehyde in the nucleus and then extracted. Fixed chromatin was digested with DpnII, the 5' overhangs filled in with biotinylated nucleotides, and then free blunt ends were ligated. After ligation, crosslinks were reversed, and DNA purified of protein. Purified DNA was treated to remove biotin that was not internal to ligated fragments. The DNA was then sheared to ~350 bp mean fragment size and sequencing libraries were generated using NEBNext Ultra enzymes and Illumina-compatible adapters. Biotin-containing fragments were isolated using streptavidin beads before PCR enrichment of each library. The libraries were sequenced on an Illumina HiSeqX. The number and length of read pairs produced for each library was: 83 million, 2x150bp for library 1; 232 million, 2x150bp for library 2. Together, these Dovetail HiC library reads provided 1,490.70x physical coverage of the genome (1-50kb pairs).

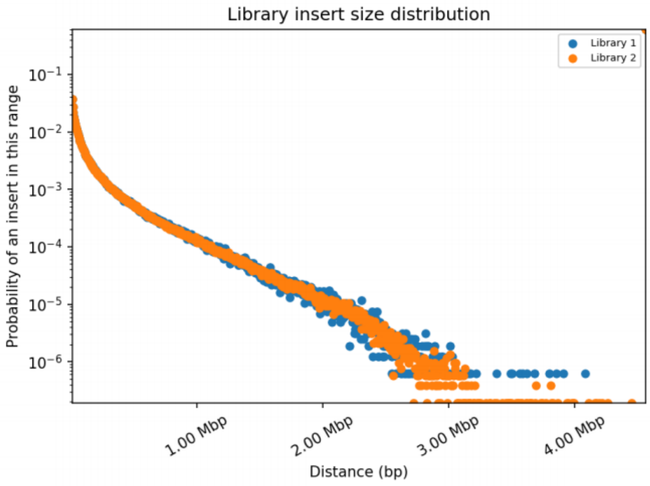

This figure shows the distribution of insert sizes in the Dovetail library. The distance between the forward and reverse reads is given on the X-axis in basepairs, and the probability of observing a read pair with a given insert size is shown on the Y-axis.

**Supplementary Figure 3:** Distribution of inserts in Dovetail libraries. Figure supplied by Dovetail genomics.

***Scaffolding the assembly with* HiRiSE*:***

The input de novo assembly, shotgun reads, Chicago library reads, and Dovetail HiC library reads were used as input data for HiRiSE™, a software pipeline designed specifically for using proximity ligation data to scaffold genome assemblies [20]. An iterative analysis was conducted. First, Shotgun and Chicago library sequences were aligned to the draft input assembly using a modified SNAP read mapper (http://snap.cs.berkeley.edu). The separations of Chicago read pairs mapped within draft scaffolds were analyzed by HiRiSE to produce a likelihood model for genomic distance between read pairs, and the model was used to identify and break putative misjoins, to score prospective joins, and make joins above a threshold. After aligning and scaffolding Chicago data, Dovetail HiC library sequences were aligned and scaffolded following the same method. After scaffolding, shotgun sequences were used to close gaps between contigs (Supplementary Figures 3, 4). The final assembly contained 26 scaffolds of 1 M bp size or greater, as seen in Fig 2 (Main Text), with further details in Supplementary Note 3.1.

Estimated physical coverage (10-10,000 kb pairs): 1,490.70X

| | Input Assembly | Dovetail HiRise Assembly |
|---|---|---|
| Total Length | 325.68 Mb | 325.72 Mb |
| L50/N50 | 92 scaffolds; 1.022 Mb | 11 scaffolds; 9.884 Mb |
| L90/N90 | 448 scaffolds; 0.089 Mb | 118 scaffolds; 0.099 Mb |

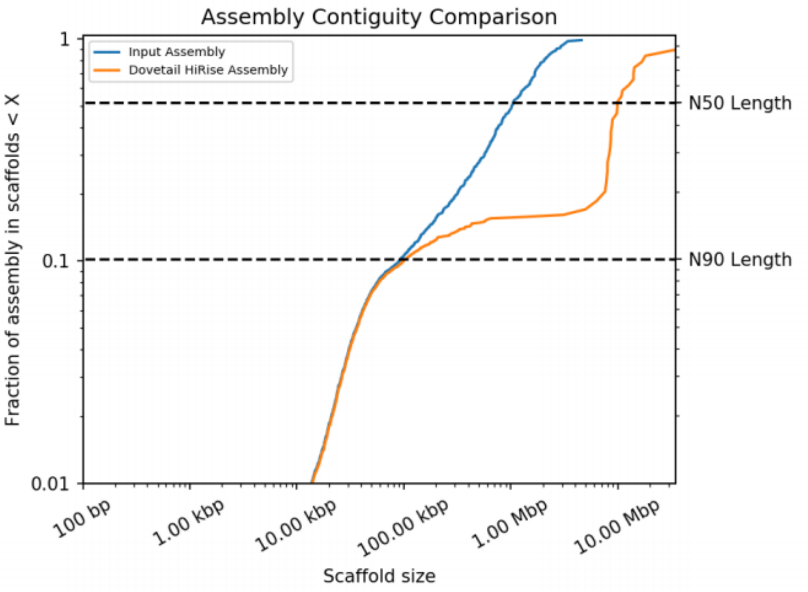

A comparison of the contiguity of the input assembly and the final HiRise scaffolds. Each curve shows the fraction of the total length of the assembly present in scaffolds of a given length or smaller. The fraction of the assembly is indicated on the Y-axis and the scaffold length in basepairs is given on the X-axis. The two dashed lines mark the N50 and N90 lengths of each assembly. Scaffolds less than 1 kb are excluded.

**Supplementary Figure 4:** Assembly contiguity comparison, supplied by Dovetail Genomics.

### Supplementary Note 3: Assembly comparisons

#### 3.1 Assembly metrics and comparisons

The assembly of the *Ephydatia muelleri* genome is the best yet available for a species of Porifera. A variety of statistics related to the assembly can be seen in Supplementary Tables 2 and 3 below. In particular, the assembly is contained on relatively few scaffolds (1,445) with a high N50 (9,883,643 bp) and with 11 scaffolds in the N50 group. Twenty-six scaffolds are 1 Mb or longer in size.

This represents a significant improvement on previously published poriferan datasets. Previously, the best available genome by many metrics was that of *Sycon ciliatum,* although the high 'N' content of that assembly (22.1% 'N') means that the scaffolds in the *S. ciliatum* genome contain many uninformative regions. The assembly of *E. muelleri* is assembled on fewer scaffolds, to a higher contiguity, and with far fewer gaps (0.05% N, more detail in Supplementary Note 3.4 below) than any other sponge resource. This is clearly apparent in Supplementary Figure 5 below.

The GC content of the assembly, 43.11%, is in line with that seen in other species, and within the normal range for eukaryotic genomes [22]. It is relatively close to the figure seen in the draft genome of *L. baikalensis* [23], which makes sense given the close relationship between these freshwater sponges.

The genome size of *E. muelleri* was first estimated to be 357 Mb (diploid content = 0.73±0.03 pg [24]). Our genome assembly contains slightly less than this figure, and is more in line with the later haploid genome size estimates [25], of 0.33 to 0.34 pg (323-332 Mb). This figure corresponds well with the 325 Mb found in our assembly. Any missing portions of the genome may be repetitive elements not well-recovered by the assembly process, particularly centromere and telomere portions, which were not noted in our data (see Supplementary Note 3.4).

The genome size of *E. muelleri* is slightly higher than the average found in demosponges [25]. The mean haploid genome size seen for sponges is around 200 Mb [25], and *E. muelleri* was the 8th largest genome of the 75 species measured. This expanded genome was nevertheless well recovered, as noted above, which suggests that the strategy used here would be good when sequencing other species in the future.

Compared to other commonly used genomes, and particularly well-known non-sponge resources (Supplementary Table 3) our data is of high quality. The *E muelleri* genome is better assembled than many smaller genomes commonly used as model species and key comparison points when discerning the origin of metazoan traits. The *E. muelleri* genome is empirically better assembled than the genomes of *A. queenslandica M. leidyi, N. vectensis, T. adherens* and *B. floridae* by almost any metric. This is a natural consequence of improvements in genome sequencing technology, but strongly recommends our resource for comparative work going forward.

**Supplementary Table 2:** Length and composition statistics for sponge genomes. For data sources, see next page.

| | ***E muelleri*** | *A queenslandica* | *T wilhelma* | *L baikailensis* | *O carmela* | *S ciliatum* |
|---|---|---|---|---|---|---|
| # of sequences | **1445** | 13397 | 5936 | 135191 | 67767 | 7780 |
| Total length (nt) | **325717041** | 166679601 | 125670620 | 209989122 | 57775306 | 357509570 |
| Longest sequence (nt) | **34737626** | 1888931 | 659656 | 124926 | 107672 | 1380240 |
| Mean sequence length (nt) | **225410** | 12442 | 21171 | 1553 | 853 | 45952 |
| Median sequence length (nt) | **23056** | 1939 | 6691 | 845 | 158 | 9478 |
| N50 sequence length (nt) | **9,883,643** | 120,365 | 73,701 | 2,213 | 5,457 | 169,232 |
| N50 sequence count | **11** | 309 | 483 | 19573 | 1950 | 597 |
| # of sequences > 10K (nt) | **1071 (74.1%)** | 1996 (14.9%) | 2075 (35.0%) | 1820 (1.3%) | 1069 (1.6% ) | 3649 (46.9%) |
| # of sequences > 100K (nt) | **115 (8.0%)** | 378 (2.8% ) | 295 (5.0% ) | 3 (0.0%) | 1 (0.0%) | 1134 (14.6%) |
| # of sequences > 1M (nt) | **26 (1.8%)** | 5 (0.0% ) | 0 (0.0%) | 0 (0.0%) | 0 (0.0%) | 2 (0.0%) |
| # of sequences > 10M (nt) | **9 (0.6%)** | 0 (0.0% ) | 0 (0.0%) | 0 (0.0%) | 0 (0.0%) | 0 (0.0%) |
| Sum length of sequences > 1M (nt) (%of total length) | **274385796 (84.2%)** | 6492255 (3.9%) | 0 (0.0%) | 0 (0.0%) | 0 (0.0%) | 2398837 (0.7%) |
| Sum length of sequences > 10M (nt) | **146175339 (44.9%)** | 0 (0.0% ) | 0 (0.0%) | 0 (0.0%) | 0 (0.0%) | 0 (0.0%) |
| GC-content (%) | **43.11** | 35.82 | 39.92 | 43.76 | 43.52 | 46.99 |

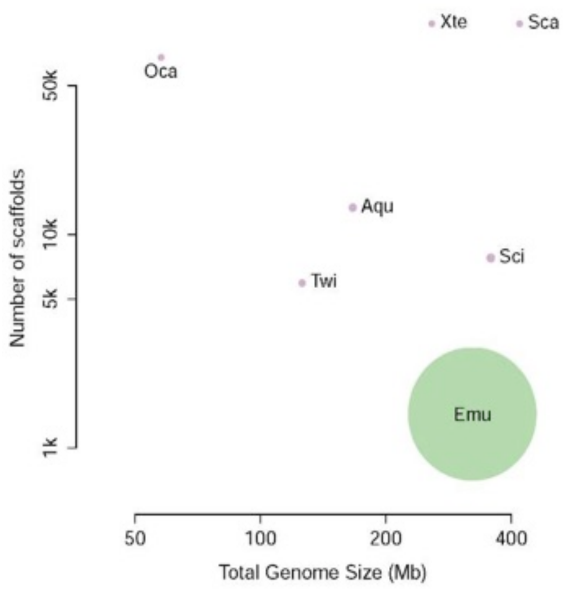

**Supplementary Figure 5:** Comparison of N50 (diameter of circles) of sponge genomes by scaffold number and genome size. (Emu, *Ephydatia muelleri*; Aqu, *Amphimedon queenslandica*; Sci, *Sycon ciliatum*, Twi, *Tethya wilhelma*; Oca, *Oscarella carmela*; Xte, *Xestospongia testudinaria*; Sca, *Stylissa carteri*)

**Supplementary Table 3:** Length and composition statistics, for non-vertebrate animal and *Capsaspora owczarzaki* genomes.

| | *C. owzarzaki* | ***E. muelleri*** | *A. queenslandica* | *M. leidyi* | *N. vectensis* | *T. adhaerens* | *B. floridae* |
|---|---|---|---|---|---|---|---|
| # of sequences | 84 | **1445** | 13397 | 5100 | 10804 | 1415 | 398 |
| Total length (nt) | 27967784 | **325717041** | 166679601 | 155865547 | 356613585 | 105632827 | 521895125 |
| Longest sequence (nt) | 3794338 | **34737626** | 1888931 | 1222598 | 626 | 13260704 | 11512737 |
| Mean sequence length (nt) | 332950 | **225410** | 12442 | 30562 | 626 | 74652 | 1311294 |
| Median sequence length (nt) | 9614 | **23056** | 1939 | 1772 | 6708 | 2278 | 750207 |
| N50 sequence length (nt) | 1,617,775 | **9,883,643** | 120,365 | 187,314 | 472,588 | 5,978,658 | 2,586,727 |
| L50 sequence count | 6 | **11** | 309 | 242 | 181 | 6 | 62 |
| # of sequences > 10K (nt) | 40 (47.6%) | **1071 (74.1%)** | 1996 (14.9%) | 1023 (24.6%) | 2969 (27.5%) | 174 (12.3%) | 398 (100.0%) |
| # of sequences > 100K (nt) | 19 (22.6%) | **115 (8.0%)** | 378 (2.8%) | 509 (10%) | 529 (4.9%) | 43 (3.0%) | 364 (91.5%) |
| # of sequences > 1M (nt) | 12 (14.3%) | **26 (1.8%)** | 5 (0.0%) | 1 (0.0%) | 66 (0.6%) | 21 (1.5%) | 169 (42.5%) |
| # of sequences > 10M (nt) | 0 (0.0%) | **9 (0.6%)** | 0 (0.0%) | 0 (0.0%) | 0 (0.0%) | 1 (0.1%) | 2 (0.5%) |
| Sum length of sequences > 1M (nt) (%of total length) | 22641449 (81.0%) | **274385796 (84.2%)** | 6492255 (3.9%) | 1222598 (0.8%) | 101305513 (28.4%) | 87945733 (83.3%) | 439987787 (84.3%) |
| Sum length of sequences > 10M (nt) | 0 (0.0%) | **146175339 (44.9%)** | 0 (0.0% ) | 0 (0.0%) | 0 (0.0%) | 13260704 (12.6%) | 22087389 (4.2%) |
| GC-content (%) | 53.82 | **43.11** | 35.82 | 38.86 | 40.64 | 32.74 | 41.19 |

**Data sources (NB also used for Supplementary Note 7):**

- ***Mnemiopsis leidyi:*** https://research.nhgri.nih.gov/mnemiopsis/
- ***Capsaspora owczarzaki***: https://protists.ensembl.org/Capsaspora_owczarzaki_atcc_30864/Info/Index
- ***Nematostella vectensis***: ftp://ftp.ensemblgenomes.org/pub/metazoa/release-44/fasta/nematostella_vectensis/dna/Nematostella_vectensis.ASM20922v1.dna.toplevel.fa.gz
- ***Trichoplax adhaerens:*** https://genome.jgi.doe.gov/portal/Triad1/Triad1.download.ftp.html
- ***Branchiostoma floridae:*** https://genome.jgi.doe.gov/portal/Brafl1/Brafl1.download.html
- ***Amphimedon queenslandica***: ftp://ftp.ensemblgenomes.org/pub/release-42/metazoa/fasta/amphimedon_queenslandica/dna/Amphimedon_queenslandica.Aqu1.dna.nonchromosomal.fa.gz
- ***Lubomirskia baikailensis***: https://ndownloader.figshare.com/files/12403526?private_link=fe36239c32bbf7342756

- ***Tethya wilhelma***: https://bitbucket.org/molpalmuc/tethya_wilhelma-genome/src/master/gene_sets
- ***Oscarella carmela***: http://www.compagen.org/datasets/OCAR_WGA_120614.zip
- ***Sycon ciliatum:*** http://www.compagen.org/datasets/SCIL_WGA_130802.zip
- ***Stylissa carteri*** http://sc.reefgenomics.org/
- ***Xestospongia testudinaria*** http://xt.reefgenomics.org/

**3.2 BUSCO results**

To gain an understanding of the completeness of our genome we used the Benchmarking Universal Single-Copy Orthologs (BUSCO v2/3) in genome mode against the eukaryotic set, which are 303 genes found in single copy in a select group of eukaryotic genomes [26]. The recovery of these from our genome is shown below alongside those of other sponges (Supplementary Table 4) and of other non-vertebrate animals and *C. owczarzaki* (Supplementary Table 5). BUSCO scores and statistics were obtained via GVolante (https://gvolante.riken.jp/analysis.html).

Our *E. muelleri* resource was annotated using the genome mode to contain 83.83% of the eukaryotic set, with 49 genes (16.17%) noted as missing from our dataset. This is represents less comprehensive recovery than that observed in most genomes, but is nonetheless a high level of recovery. However, when the protein set derived from AUGUSTUS (see Supplementary Note 5.1 below) was used in protein mode on the GVolante server, only 30 (9.90%) of genes were noted as missing.

Compared to other sponge resources (Supplementary Table 4), the *E. muelleri* resource exhibits better recovery than *O. carmella* and *S. ciliatum*, approximately the same level as *T. wilhelma,* and slightly inferior metrics compared to *A. queenslandica,* and *L. baikalensis.* The number of genes recovered with the transcriptome is much higher, however (Supplementary Table 4). This could be a result of incomplete assembly or gene loss in *E. muelleri,* or a divergent gene set not recognised by the BUSCO algorithm. When compared to the genomes of other organisms (Supplementary Table 5) the level of missing genes seen in *E. muelleri* is also slightly higher than those seen in other datasets. To check whether these genes had simply not been annotated, we checked the published *E. muelleri* transcriptome [27] to see whether any missing BUSCO representatives could be noted there (using coding/transcribed (nucleotide) mode on the GVolante server). In that resource, only 16 genes, 5.28% of the total set, were missing (with 279 complete genes, and 8 partial genes). This represents 94.72% recovery, better than almost any genome set listed in the tables below. It therefore seems likely that at least 14 of the 30 missing BUSCO genes do exist in *E. muelleri,* although they are not recognised in our protein models.

**Supplementary Table 4:** Comparison of BUSCO Scores among sponge genomes.

| | ***E. muelleri* (genome mode)** | ***E. muelleri* (Augustus aa set)** | *A. queenslandica* | *T. wilhelma* | *L. baikalensis* | *O. carmella* | *S. ciliatum* |
|---|---|---|---|---|---|---|---|
| Total # of core genes queried | **303** | **303** | 303 | 303 | 303 | 303 | 303 |
| # of core genes detected | **243 (80.20%)** | **254 (83.83%)** | 274 (90.43%) | 258 (85.15% ) | 263 (86.80%) | 207 (68.32%) | 240 (79.21% ) |
| Complete + Partial | **254 (83.83%)** | **273 (90.10%)** | 281 (92.74%) | 275 (90.76% ) | 277 (91.42%) | 237 (78.22%) | 259 (85.48% ) |
| # of missing core genes | **49 (16.17%)** | **30 (9.90%)** | 22 (7.26%) | 28 (9.24%) | 26 (8.58%) | 66 (21.78%) | 44 (14.52% ) |
| Average # of orthologues per core genes | **1.12** | **1.22** | 1.04 | 1.05 | 1.15 | 1 | 1.07 |
| % of detected core genes that have more than 1 ortholog | **10.7** | **19.6** | 4.38 | 4.26 | 12.93 | 0 | 6.67 |

**Supplementary Table 5:** Comparison of BUSCO scores of non-vertebrate animals and *C. owczarzaki.*

| | ***E. muelleri* (genome mode)** | ***E. muelleri* (AUGUSTUS aa set)** | *A. queenslandica* | *C. owczarzaki* | *M. leidyi* | *N. vectensis* | *T. adherens* | *B. floridae* |
|---|---|---|---|---|---|---|---|---|
| Total # of core genes queried | **303** | **303** | 303 | 303 | 303 | 303 | 303 | 303 |
| # of core genes detected | **243 (80.20%)** | **254 (83.83%)** | 274 (90.43%) | 88 (95.05%) | 278 (91.75%) | 274 (90.43%) | 270 (89.11%) | 277 (91.42%) |
| Complete + Partial | **254 (83.83%)** | **273 (90.10%)** | 281 (92.74%) | 289 (95.38%) | 293 (96.70%) | 283 (93.40%) | 274 (90.43%) | 286 (94.39%) |
| # of missing core genes | **49 (16.17%)** | **30 (9.90%)** | 22 (7.26%) | 14 (4.62%) | 10 (3.30%) | 20 (6.60%) | 29 (9.57%) | 17 (5.61%) |
| Average # of orthologues per core genes | **1.12** | **1.22** | 1.04 | 1.01 | 1.01 | 1.03 | 1.01 | 1.05 |
| % of detected core genes that have more than 1 ortholog | **10.7** | **19.6** | 4.38 | 1.04 | 0.72 | 2.19 | 0.74 | 4.69 |

The *E. muelleri* genome contains a higher level of duplication in the BUSCO set of genes than seen in most genomes. The average number of orthologues per core gene, 1.12 (1.22 in our protein models), and the percentage of core genes with multiple orthologues, 10.7%, (19.6% in protein models) is higher by some margin than any other genome or the draft genome resource for *L. baikalensis,* which as an incomplete resource may retain heterozygous portions collapsed in better assemblies. This data, both absence and expansion, indicates that *E. muelleri* therefore may exhibit a more labile gene complement than other species, as even these highly conserved genes seem to be subject to loss and duplication in our resource.

Gene duplications are prevalent in our gene set, and can be seen frequently in our genome browser, with related genes, the product of local duplication events, often lying adjacent to one another. An example of these duplications is shown in Supplementary Figure 6 below, showing three *integrin alpha* genes on scaffold 4, taken from a region with large numbers of interspersed *integrin alpha* and *integrin beta* genes. These duplicates are sometimes, but not always, identical in intron/exon structure. The high level of duplication is commented upon more generally in Supplementary Note 7.

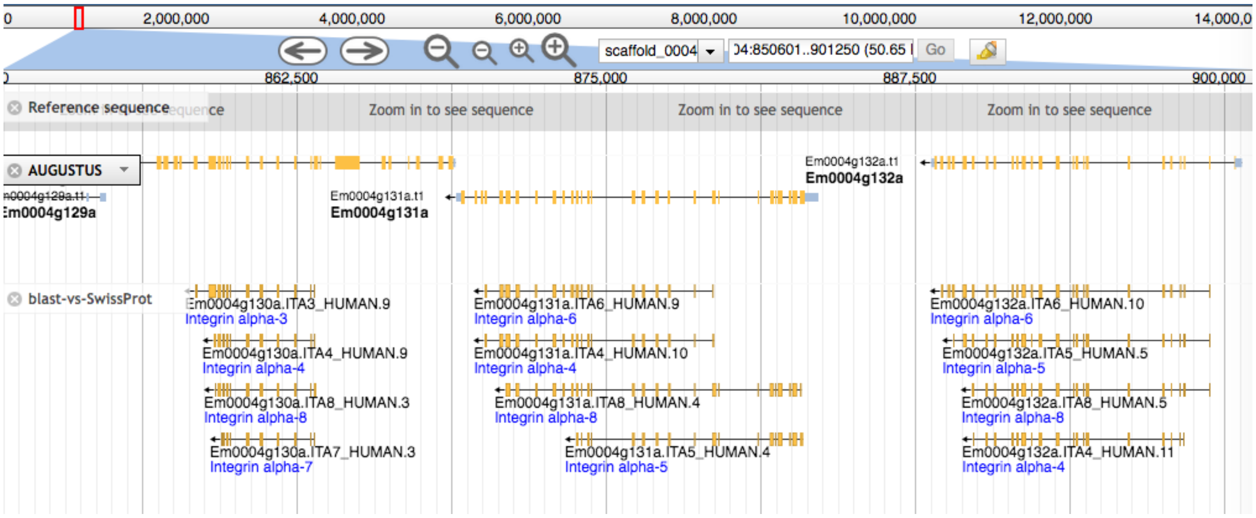

**Supplementary Figure 6.** Example of the prevalent duplication of genes. A screenshot from the genome browser shows three *integrin alpha* genes from a region where integrins are duplicated on scaffold 4 (Em0004g129a, Em0004g131a and Em0004g132a). The structure of these genes (and support from RNAseq data) is similar but not identical to genes located in *cis*, indicating that these are real genes, and not the product of genome assembly artifacts.

#### 3.3 Repeat content

To ascertain repeat content from both nuclear elements and simple repeats in the genomes of *Ephydatia muelleri* and other sequenced sponge genomes, RepeatModeler 2.0 and RepeatMasker 4.1.0 were used sequentially [28]. This process first generates libraries of repetitive sequences *de novo* from input genome sequences, and then identifies, quantifies and masks these sequences. As sponges do not have well-curated libraries of repeats, this process is the first step to generating a strong understanding of the repeat content of sponge genomes.

A number of patterns can be seen in the four sponge genomes examined. In all species, Unclassified repeats are the most common group of repeats (19.27 - 31.61% of the genome, Supplementary Table 6 below). This reflects the fact that almost all the repeat element families seen in sponges are yet to be formally classified, and as such they are not yet represented in existing libraries, at least to the level of conservation that can be recovered by blastn (performed by RepeatModeler). As sponge genomic resources become more available, it will be possible to track the gain and loss of these elements across poriferan phylogeny. Already from our data it is evident that some repeat elements are more common in individual species, and absent from others. SINES, for example, are present in *Sycon ciliatum* and *Tethya wilhelma,* making up 0.12% and 0.23% of these genomes, but are completely absent from *Amphimedon queenslandica* and *E. muelleri.*

*Ephydatia muelleri* in particular has many LTR and DNA elements compared to the other sponges examined here. It is possible that these expansions are not uniform across all *E. muelleri* populations, and differ in number between different sub-populations. *E. muelleri* also contains around three times more simple repeats than the other sponge genomes examined (6.72%, cf. 1.96-2.43%). However, in part this will be an artifact of the higher quality of genome assembly of *E. muelleri.* In past work, simple repeats have been difficult to assemble [29], whereas the *E muelleri* assembly is so contiguous that repeats are incorporated into the scaffolding.

It is worthwhile considering genome sizes, in light of the quantity of repeat elements contained in each genome. *Tethya wilhelma* has a small genome (125Mb) and a small number of repeats (30%). *Amphimedon queenslandica* has a small genome but a medium-to-high number of repeats (43%) within its notably compact genome structure. *Sycon ciliatum* has a large genome relative to other sponges (357 Mb), and many already masked regions (22.1% 'N', largely inserted as a consequence of scaffolding). When added to the 27.8% of repeats noted by RepeatMasker, the total sequence masked in the *S. ciliatum* dataset is 49.1%, although it should be noted that many of these Ns are a result of scaffolding rather than masking. The genome of *Ephydatia muelleri* is also large (325 Mb) and the number of repeats also high (47.04%).

Genomic expansion therefore seems linked to repeat content. Larger genomes of sponges examined here have more repeats, both proportionally and as a total number of base pairs within the genome. The sampling size is however very small, and wider sampling across the sponge tree of life is likely to be informative. Larger genomes commonly exhibit repetitive element expansions (e.g.[30]). While not universal, larger genome sizes and number of repetitive sequences often go together [31]. The size of the *Ephydatia muelleri* genome may be partly due to its larger number of repetitive sequences.

This percentage of repeat content is higher than some animals, but in general unsurprising. Repeat content of eukaryotes can range from around 10% to around 80%, with plants (e.g. *Helianthus annuus* (sunflower)) often exhibiting higher numbers [32]. Within non-bilaterian animals, placozoans are known to have low numbers of repetitive sequence [33]. Cnidarians possess between 26.2% and 57.64% transposon content, with larger genomes (such as *Hydra vulgaris*) possessing larger numbers of transposable elements, notably non-LTRs[33].

**Supplementary Table 6:** Repeat content of the sponges *Ephydatia muelleri*, *Tethya wilhelma, Amphimedon queenslandica,* and *Sycon ciliatum*. Repeats were annotated using RepeatModeller and RepeatMasker.

***Ephydatia muelleri***

| Element | Number | Length Occupied (bp) | % of Genome |
|---|---|---|---|
| SINES | 0 | 0 | 0 |
| MIRs | 0 | 0 | 0 |
| LINES | 24516 | 10352233 | 3.21 |
| LINE1 | 0 | 0 | 0 |
| LINE2 | 0 | 0 | 0 |
| L3/CR1 | 1673 | 1079942 | 0.33 |
| LTR elements | 20404 | 13977147 | 4.33 |
| ERV- class 2 | 0 | 0 | 0 |
| DNA elements | 54056 | 22038945 | 6.83 |
| hAT-Charlie | 0 | 0 | 0 |
| TcMar-Tigger | 241 | 109076 | 0.03 |
| Unclassified | 296587 | 83250207 | 25.8 |
| Total interspersed repeats | | 129618532 | 40.18 |
| Small RNA | 0 | 0 | 0 |
| Satellites | 0 | 0 | 0 |
| Simple repeats | 166923 | 21695785 | 6.72 |
| Low complexity | 8714 | 923911 | 0.29 |
| **Total length:** | 322620753 | **Bases masked** | 151771384 (47.04%) |

***Tethya wilhelma***

| Element | Number | Length Occupied (bp) | % of Genome |
|---|---|---|---|
| SINES | 1926 | 150121 | 0.12 |
| MIRs | 751 | 40859 | 0.03 |
| LINES | 2093 | 836649 | 0.67 |
| LINE1 | | | |
| LINE2 | 234 | 76836 | 0.06 |
| L3/CR1 | 608 | 250675 | 0.2 |
| LTR elements | 4573 | 1834500 | 1.46 |
| ERV- class 2 | 1136 | 156065 | 0.12 |
| DNA elements | 12516 | 3203506 | 2.55 |
| hAT-Charlie | 45 | 10066 | 0.01 |
| TcMar-Tigger | 471 | 95248 | 0.08 |
| Unclassified | 121669 | 28488640 | 22.67 |
| Total interspersed repeats | | 35413416 | 27.46 |
| Small RNA | 96 | 16573 | 0.01 |
| Satellites | 93 | 27930 | 0.02 |
| Simple repeats | 51427 | 2463746 | 1.96 |
| Low complexity | 2362 | 110446 | 0.09 |
| **Total length:** | 125670620 | **Bases masked** | 37073370 (29.50%) |

***Amphimedon queenslandica***

| Element | Number | Length Occupied (bp) | % of Genome |
|---|---|---|---|
| SINES | 0 | 0 | 0 |
| MIRs | 0 | 0 | 0 |
| LINES | 12893 | 2691209 | 1.86 |
| LINE1 | 795 | 459019 | 0.32 |
| LINE2 | 9026 | 886177 | 0.61 |
| L3/CR1 | 1011 | 500038 | 0.35 |
| LTR elements | 7811 | 3244519 | 2.24 |
| ERV- class 2 | 0 | 0 | 0 |
| DNA elements | 23925 | 6031236 | 4.16 |
| hAT-Charlie | 162 | 106189 | 0.07 |
| TcMar-Tigger | 91 | 20613 | 0.01 |
| Unclassified | 157693 | 45782581 | 31.61 |
| Total interspersed repeats | | 57749545 | 39.87 |
| Small RNA | 0 | 0 | 0 |
| Satellites | 3503 | 1054596 | 0.73 |
| Simple repeats | 71376 | 3521583 | 2.43 |
| Low complexity | 8538 | 496037 | 0.34 |
| **Total length:** | 144846769 | **Bases masked** | 62637881 (43.24%) |

***Sycon ciliatum***

| Element | Number | Length Occupied (bp) | % of Genome |
|---|---|---|---|
| SINES | 8911 | 818717 | 0.23 |
| MIRs | 0 | 0 | 0 |
| LINES | 36327 | 7195867 | 2.01 |
| LINE1 | 65 | 16178 | 0.00 |
| LINE2 | 7741 | 1379382 | 0.39 |
| L3/CR1 | 2145 | 503854 | 0.14 |
| LTR elements | 9406 | 2148292 | 0.6 |
| ERV- class 2 | 751 | 109488 | 0.03 |
| DNA elements | 101467 | 11680109 | 3.27 |
| hAT-Charlie | 2840 | 402092 | 0.11 |
| TcMar-Tigger | 1943 | 304745 | 0.09 |
| Unclassified | 558858 | 68900829 | 19.27 |
| Total interspersed repeats | | 90743814 | 25.38 |
| Small RNA | 1292 | 136954 | 0.04 |
| Satellites | 404 | 106033 | 0.03 |
| Simple repeats | 147020 | 7965529 | 2.23 |
| Low complexity | 4266 | 231015 | 0.06 |
| **Total length:** | 357509570 | **Bases masked** | 98589239 (27.58%) |

#### 3.4 Contiguity and scaffolding

In comparison to previous sponge genomic resources, the *Ephydatia muelleri* genome is exceptionally well-scaffolded. This can be seen clearly in Supplementary Figure 7A below, which shows the relative sizes of the scaffolds that make up these assemblies, as well as their number. The contig N50 of our assembly, 213.26 kb, in itself is higher than the scaffold N50 of previous poriferan resources, giving a firm basis for scaffolding efforts and leading to a very complete dataset.

The majority of the *Ephydatia muelleri* genome is concentrated on just 24 scaffolds. As the chromosomal *n* value of *E. muelleri* is 23 [24], these 24 large scaffolds likely represent these chromosomes, with one chromosome split into two parts. The additional smaller scaffolds will contain a variety of sequences that should be placed towards the ends of these larger scaffolds.

Vertebrate-type telomeres ($(TTAGGG)_n$ [24]) are known to be used by *Ephydatia muelleri.* However, no trace of these could be found in our assembly. If present, they should be found as long runs of (TTAGGG)n, repeated up to 3000 times. However, this motif is uncommon in our dataset. It occurs singly 7121 times across the entirety of our assembly (roughly 10% of what would be expected by chance (325Mb/4096=79,000). It occurs as a doublet twice, and never more than as a doublet, in this direction or in reverse complement.

Our assembly therefore is almost at chromosomal level, as we are unable to verify the position or quantity of telomeres in our assembly. However, our assembly cannot be far short of this standard, as the quantity of sequence within it (Supplementary Note 3.1) more than covers the predicted genome size. The *Ephydatia muelleri* genome contains 1,864,700 Ns all of which were intentionally inserted during the assembly process (1829 x 1000bp and 357 x 100 bp) and of these, 1,760,200 occur in the top 24 scaffolds. During the HiRiSE assembly process, 360 gaps were joined and 11 falsely joined portions of the input assembly were broken. This equates to 0.57% of the genome, a small percentage of gaps when compared to similar resources (Supplementary Note 3.1). We note that 434 scaffolds, totalling 5Mb, are AT repeat-rich, posing problems for assembly algorithms, and thus not incorporated well into our final genome assembly (Supplementary Figure 7B).

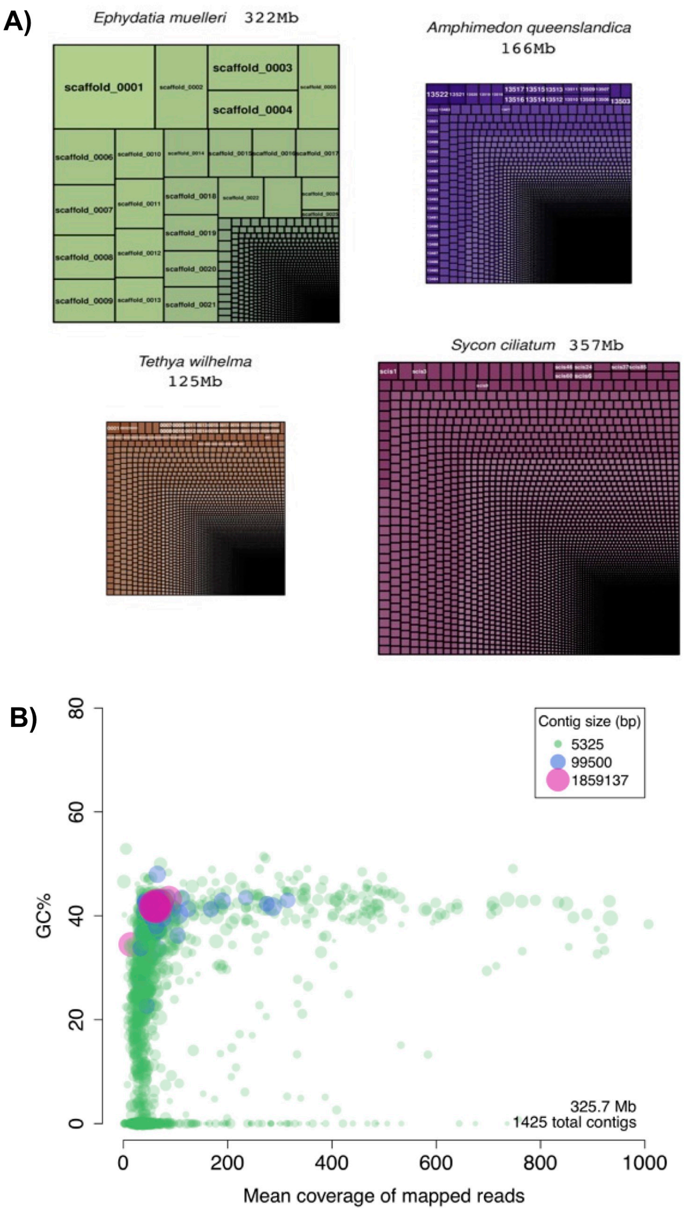

**Supplementary Figure 7:** A) Relative scaffold size in the genome of *Ephydatia muelleri*, compared to available sponge genomic resources. Figures drawn using the TreeMap package in R. B) GC content across the contigs in the *Ephydatia muelleri* genome assembly. Note a large number of small, AT rich contigs, representing repetitive areas of the genome not well-integrated into the assembly.

#### 3.5 Checks for contamination

It is possible for genome sequencing projects to contain sequences derived from a variety of exogenous sources. This can include, among other things, DNA taken up by the target organism (e.g. gut contents), contamination in the process of DNA extraction and sequencing (particularly human sequence), and microorganisms found on and alongside the target species, and gathered in the process of collection and DNA extraction. These sequences can be difficult to exclude, and can both hinder assembly and interfere with analysis by adding erroneous information derived from external sources.

**Supplementary Table 7:** Top 10 blastx hits between *E. muelleri* assembly and the swissprot bacterial database, after removal of Scaffold 25.

| Scaffold | Blast hit | % identity | length | mismatches | Gaps | Hit start | Hit end | E value |
|---|---|---|---|---|---|---|---|---|
| scaffold_0263 | sp\|A5EEQ0\|DXS_BRASB | 68.0 | 600 | 182 | 4 | 35034 | 33238 | 4.7e-229 |
| scaffold_0567 | sp\|Q9I3S3\|GBUA_PSEAE | 49.3 | 371 | 115 | 7 | 14485 | 13388 | 1.80E-90 |
| scaffold_0701 | sp\|Q1LQ00\|ADE_CUPMC | 53.4 | 283 | 130 | 1 | 11641 | 10793 | 1.90E-82 |
| scaffold_0785 | sp\|A1UUC3\|DNAK_BARBK | 53.3 | 291 | 123 | 5 | 2218 | 1364 | 2.20E-71 |
| scaffold_0696 | sp\|B3E600\|ADE_GEOLS | 59.3 | 199 | 81 | 0 | 11658 | 11062 | 4.70E-65 |
| scaffold_0786 | sp\|O54068\|UDG_RHIME | 39.4 | 363 | 210 | 6 | 2124 | 1042 | 1.40E-57 |
| scaffold_0808 | sp\|Q2LVL0\|MSBA_SYNAS | 44.6 | 267 | 121 | 3 | 11202 | 12002 | 1.10E-56 |
| scaffold_0904 | sp\|Q0AAW0\|ACSA_ALKEH | 40.8 | 287 | 88 | 7 | 16029 | 15169 | 4.50E-51 |
| scaffold_0216 | sp\|P25526\|GABD_ECOLI | 49.2 | 193 | 61 | 2 | 37761 | 37183 | 1.90E-43 |
| scaffold_0531 | sp\|C3MIE4\|BETA_SINFN | 30.3 | 587 | 319 | 20 | 21624 | 19939 | 2.90E-43 |

In sponges, most bacterial contamination is normally derived from the microorganisms that inhabit these species, and is difficult to exclude. To mitigate this, we grew gemmules from a single clone of *Ephydatia muelleri* in sterile conditions, preventing the horizontal transfer of bacteria to our specimen. In this way, we excluded a major origin point of contamination from our DNA source. However, contamination could have been introduced from a variety of other sources. To exclude this, we checked our assembly for contamination by comparison to the swissprot databases, containing sequences of known provenance. We were unable to retrieve any sequence of human origin within our assembly. As noted in the main text, one scaffold, (Scaffold_153_HRSCAF_192), was assembled as a nearly complete bacterial genome, and was clearly evident within our dataset originating from a bacterium. However, no other contamination was obvious in our dataset. Supplementary Table 7 shows the 10 best hits (blastx) in our dataset to the swissprot bacterial complement. In all cases, these hits are positioned within larger scaffolds, are short, and contain large numbers of mismatches. The *E* values recorded drop off in significance rapidly. In no case, apart from scaffold 25 above, does a single scaffold contain multiple hits to bacterial sequences. These hits are therefore more likely to represent chance similarity than they are to represent contaminating sequence. We are therefore confident that with the removal of scaffold 25, contaminating sequence from bacteria has been meaningfully excluded from our assembly.

#### 3.6 Synteny analyses

We examined synteny of *E. muelleri* with five other animals, and the choanoflagellates *Monosiga brevicollis* and *Salpingoeca rosetta*. Here we defined synteny as regions of a scaffold or chromosome derived from a common ancestor, resulting in linked genes not necessarily in the same order. Dot plots, also called synteny plots, were generated using a combination of custom Python and R scripts to display unidirectional BLAST hits showing the position of the matches on the respective scaffold (all are available in. Supplementary Data 4). AUGUSTUS gene models from *E. muelleri* were used, and five other genomic datasets for comparison. Four of these genomes were assembled from Sanger sequencing reads, resulting in highly contiguous scaffolds. However, the original annotations were de novo predictions following training by a small set of Sanger-sequenced ESTs, thus we made use of newer annotations for all four of them. For *Trichoplax adhaerens*, we used the AUGUSTUS re-annotation generated by Eitel et al [34], as the original Triad1 annotation [35] was found to contain many incomplete genes, and missing approximately 1000 others. For *Nematostella vectensis*, we used the v2 annotation [36]. For both *Monosiga brevicollis* and *Branchiostoma floridae*, both original annotations contained large numbers of falsely fused genes (see examples for *B. floridae* in [37]), therefore we used the AUGUSTUS re-annotation set from [38]. For *Salpingoeca rosetta*, we used the original genomic dataset. The genomes of the sponges *Amphimedon queenslandica* (scaffolds v1, annotations v2.1) and *Sycon ciliatum* (v1) were also used, though these were too fragmented to visualize, and thus were excluded from further analysis. We also compared the two placozoans *Hoilungia hongkongensis* and *T. adhaerens*, as the high degree of synteny [34] could serve as a positive control for the analytical strategy.

For each pair of genomes, proteins from the query species (typically *Ephydatia*) were aligned against proteins of the target genome using blastp, with an e-value cutoff of 0.001. The scaffolds of each genome, the position of protein-coding genes on the scaffolds, and the tabular blast results (for a summary see Supplementary Table 8) were all used as inputs for a custom Python 3.7 script (scaffold_synteny.py, Supplementary Data 4), which compiled a table of matches by the relative gene position on each genome. This table was used as input for the R script (draw_2d_synteny.R) to create the dot plots. Proteins were excluded from either genome if they had greater than 20 matches (i.e. large protein families or transposons), as these would likely cause random observations of synteny.

To examine the significance of the observed density of protein matches on each scaffold, we used Fisher's exact test, as was done by Srivistava et al. [35]. For scaffold *i* on genome G1 and scaffold *j* on genome G2, Fisher's exact test considers the number of matches of proteins from *i* on *j*, matches from *i* to all scaffolds on G2, matches from *j* to all scaffolds on G1, and the total number of all remaining matches.

To calibrate these results, gene order was randomized globally, or locally within each scaffold (for both genomes) using the scaffold_synteny.py script, with the options -R, or -S --double-randomize, respectively. As our downstream calculations were concerned with match density within scaffolds, the latter randomization only had apparent effects on the dot plot for the two placozoan species with highly conserved linear gene order. That is, local randomization within each scaffold disrupted the collinearity (visible in the dot plot) but not the overall significance of matches for that scaffold.

The randomized p-values were found to be highly dependent on the genomes and assemblies used in the analysis. When examining *H. hongkongensis* against *T. adhaerens*, the lowest random p-value (p=7.1e-64) was found comparing the longest scaffolds on each genome, while for *M. brevicollis* against *S. rosetta*, the lowest random p-value was 1.26e-101, also for the longest scaffold in each. This shows that scaffolds with a large number of genes are still likely to contain many matches by chance. Thus, we cautiously used a p-value threshold taken from the minimum p-value of the globally randomized set.

**Supplementary Table 8:** Synteny between *E. muelleri* and various species. For *E. muelleri,* the 24 longest scaffolds were compared against a subset of scaffolds from each target species, yielding m total comparisons (the number of hypotheses for multiple-test correction). A one sided Fisher's exact test was used for testing, with correction for multiple testing. N blocks refers to the number of significant blocks when using the minimum randomized p-value as a threshold for significance. N blocks by scaffold refers to the number of significant blocks when the threshold is calculated uniquely for each query-target scaffold pair based on the randomized p-value for that scaffold pair, rather than globally.

| Species | Target scaffolds compared | Total protein matches | Total scaffold comparisons (m) | Min p value | Min p value randomized | N blocks | N blocks by scaffold |
|---|---|---|---|---|---|---|---|
| *Monosiga* | 37 | 5968 | 888 | 1.37e-81 | 9.71e-84 | 0 | 208 |
| *Salpingoeca* | 35 | 6109 | 840 | 3.03e-73 | 5.54e-64 | 2 | 246 |
| *Trichoplax* | 31 | 6883 | 744 | 0 (1e-300) | 2.60e-113 | 14 | 103 |
| *Nematostella* | 31 | 2180 | 744 | 5.73e-120 | 7.25e-28 | 30 | 112 |
| *Branchiostoma* | 40 | 3716 | 960 | 3.02e-193 | 6.65e-46 | 21 | 157 |

### Comparison between placozoan species as a control

Within the subset of species examined, two placozoan species, *T. adhaerens* and *H. hongkongensis*, were compared because these two species are closely related and were shown to have large syntenic blocks of genes [34]. The randomized p-values were much greater than all of the syntenic blocks identified by Eitel et al. [34] and suggested that 104 blocks could be identified this way (versus 162 with lower p-values compared to the same scaffold pair when randomized). The assembly of *H. hongkongensis* is comparatively fragmented relative to *T. adhaerens*, so some of these blocks may be subparts of chromosomes that are not scaffolded in the assembly. For example, *T. adhaerens* scaffold_2 had significant matches to several contigs from *H. hongkongensis* (contig_002, 007, 008, 009), evident on the dot plot from collinear regions found across the scaffold. These contigs probably belong to a much larger scaffold that would be syntenic with the *T. adhaerens* scaffold, though it is possible that substantial translocations have occurred.

**Comparison of *E. muelleri* to choanoflagellates**

We also checked whether we could observe evidence of conserved syntenic arrangements between *E. muelleri* and each of two choanoflagellate species, *M. brevicollis* and *S. rosetta* (Supplementary Figure 8), as well as between the two choanoflagellate species themselves. When *E. muelleri* was compared to either *M. brevicollis* or *S. rosetta,* little evidence of conserved syntenic relationships was observed in our dot plots. However, clear syntenic relationships can be seen when these two choanoflagellate species are compared with one another. Given the syntenic relationships known to be preserved between *E. muelleri* and other metazoan species, and between other metazoan species more generally, this seems to indicate a clear difference between the genomes of metazoans and those of their choanoflagellate ancestors, with strong conservation within metazoans themselves. Two hypotheses may explain this. Either the observed synteny within metazoans and the absence within choanozoa reflects some biological change in metazoans; in the context of the dosage mechanisms discussed already, this could mean that some metazoan innovations require precise dosage in a way that was not constrained in the choanozoan ancestor, or the few scaffolds that were identified as having significant matches between *E. muelleri* and a choanoflagellate represent bona fide synteny, and that most other regions are simply below the detection limit of this approach. In the latter case, the processes controlling synteny would not be different between choanoflagellates and animals, and a relationship may be evident if more species with equally contiguous genomes were studied.

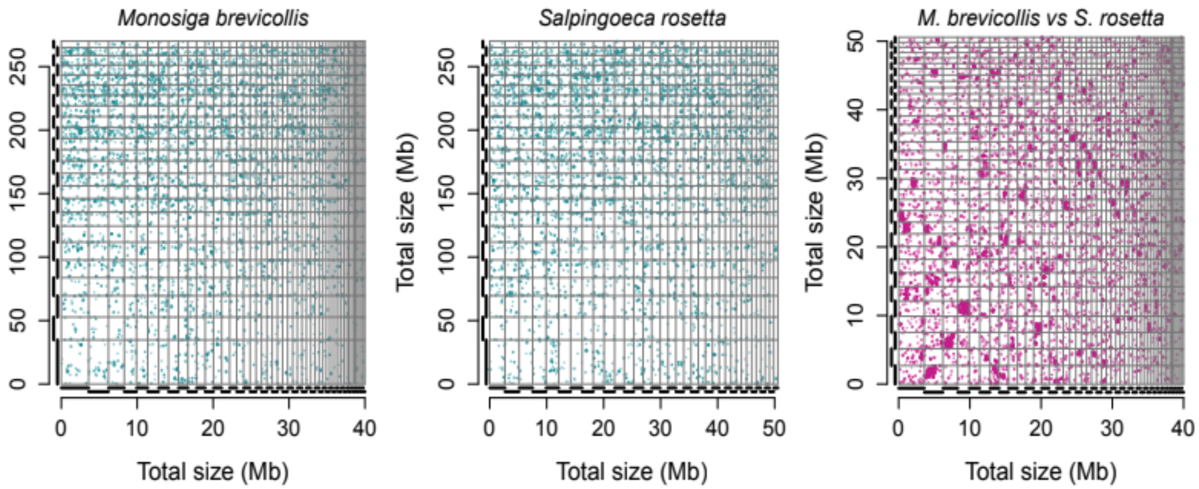

**Supplementary Figure 8:** Dot plot matrices showing gene arrangements in *Ephydatia muelleri* compared individually to two species of choanoflagellate (left and middle), and (right) these choanoflagellate species compared to one another.

### Supplementary Note 4: Analysis of scaffold 25: *Flavobacterium* sp. Genome

As noted in Supplementary Note 3.5 above, one scaffold, #25, was found to be assembled into a large piece of a bacterial genome. This was investigated as described below, and was found to belong to *Flavobacterium* sp., an as yet undescribed species. It has a genome size of 3.09Mb, a GC content of 0.3456, and when re-annotated was found to contain 3,811 genes (2,433 with an annotation), with coding density = 85.2% (Supplementary Data 1).

#### 4.1 Taxonomy assignment

CheckM 1.0.12 [39], run using the genome sequence, indicated that this belongs to a bacterium from the family Flavobacteriaceae. The genome completeness was estimated to be 89.38%, with 0.29% contamination. This was calculated using 81 genomes with 511 markers (database used by CheckM). The 16S rRNA sequence of *E. muelleri*, *Flavobacterium* sp., was blasted against the total NCBI 16S database with the following results:

- *Flavobacterium succinicans* (BLASTN similarity ~97%)
- *Flavobacterium fluvii* (BLASTN similarity ~97%)
- *Flavobacterium flevense* (BLASTN similarity ~96%)

##### 4.1.1 Average nucleotide identity (ANI)

**a. 16S ANI**

Average nucleotide identity (ANI) of the flavobacterium was analysed vs related species based on the 16S rRNA gene phylogeny, and using the OrthoANIu algorithm (https://www.ezbiocloud.net/tools/an). The ANI results were as follows: (>98.5% would be considered to be the same species, * indicates the genome is available):

1. NR_159121.1 — *Flavobacterium luteum* — 97.3%
2. NR_108537.1 — *Flavobacterium myungsuense* — 96.40%
3. NR_157632.1 — *Flavobacterium soyangense* — 96.85%
4. NR_042999.1* — *Flavobacterium weaverense* — 95%
5. NR_043000.1* — *Flavobacterium segetis* — 95.7%
6. NR_116173.1* — *Flavobacterium fluvii* — 96.10%
7. NR_108535.1 — *Flavobacterium yonginense* — 94.0%
8. NR_104712.1* — *Flavobacterium flevense* NBRC 14960 — 94.70%
9. NR_114992.1 * — *Flavobacterium flevense* DSM 1076 — 95.10%
10. NR_134036.1 — *Flavobacterium oryzae* — 95.5%

**b. Genome ANI**

Average nucleotide identity of the flavobacterium was also analysed using fastANI v1.1 [40] with all the genomes from the family Flavobacteriaceae (n = 385). Results were as follows: (* = not in 16S rRNA gene phylogeny):

- **Flavobacterium alvei* ASM292089v1 80.4224 (16S 94.85%)
- **Flavobacterium fluvii* DSM 19978 80.2876 (Similar to *NR_116173.1)
- **Flavobacterium* sp. GS13 ASM435522v1 80.0374
- *Flavobacterium flevense* DSM 1076 78.5074
- *Flavobacterium flevense* NBRC 14960 78.3723
- *Flavobacterium weaverense* 78.2615
- *Flavobacterium segetis* 78.1292

Hits to genomes not found in the 16S rRNA gene phylogeny were from species where the 16S data was not uploaded to the nucleotide database and only in the genome database, such as *Flavobacterium* sp. GS13. (Searches of all sequences of the same taxonomy found one hit with a 16S locus, MF685248.) The BLAST hit between the 16S sequence of *E. muelleri Flavobacterium* 16S and *Flavobacterium* sp. GS13 is 96.33% similar.

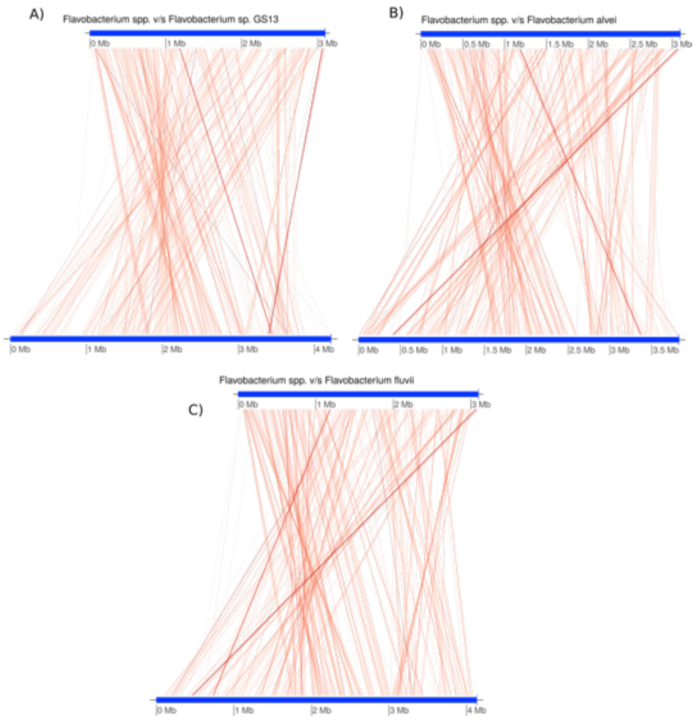

**Supplementary Figure 9:** Plot of the best hits on the genome ANI analysis. Each red line compares a region of 3000bp and calculates the ANI scores between the *Flavobacterium* sp. and the other genomes from the Family Flavobacteriaceae. A) Comparison to *Flavobacterium* sp GS13, B) Comparison to *Flavobacterium alvei*, C) Comparison to *Flavobacterium fluvii.*

The first three genomes were used to create a plot of conserved regions by reciprocal mapping of ANI values with fastANI. They were also checked with the ANI calculator from http://enve-omics.ce.gatech.edu/ani/. These plots (Supplementary Figures 9 and 10) show that while the genomes have high levels of ANI, there has been considerable rearrangement, and our *Flavobacterium* sp sequence does not correspond exactly to these known sequences, although it is related to them.

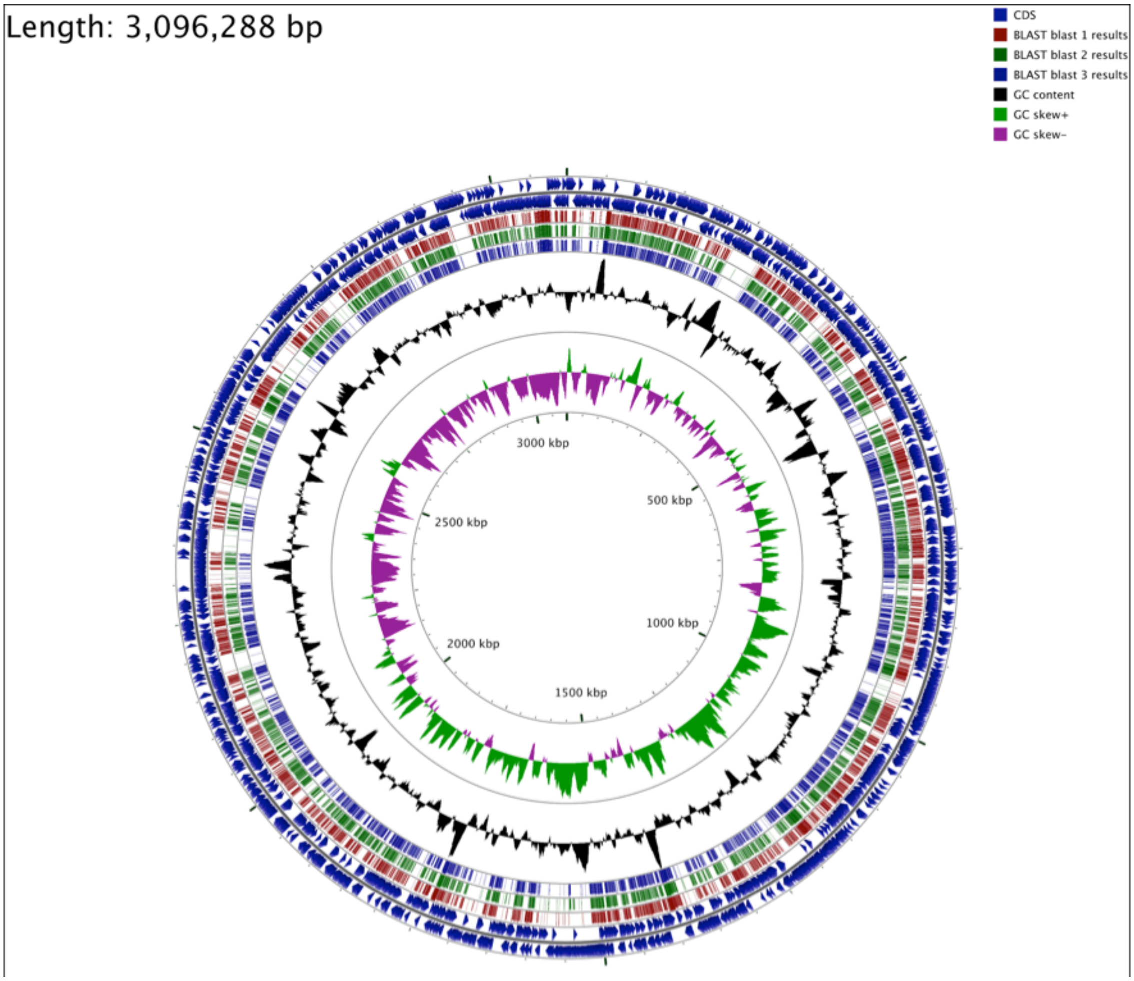

**Supplementary Figure 10:** Genome comparison of *Flavobacterium* sp. with the first three most related species as suggested by the ANI analysis (see Supplementary Note 4.1.1). Outer two circles are the CDS (forward and reverse). Next three layers are the blast (identity_cutoff=75) results of the genomes: Blast 1 - *Flavobacterium* sp. GS13, Blast 2 - *Flavobacterium fluvii*, and Blast 3 - *Flavobacterium alvei.* Internal circles show the GC content and the GC skew analysis.

#### 4.1.2 - Microbial genome atlas - MiGA

Analysis using the Microbial Genome Atlas (MiGA [41]) suggests that the *Flavobacterium* sp. genome is of high quality. MiGA was used to conduct a number of automated analyses using the standard pathways on that platform. The closest relatives found by MiGA in the database were:

- *Flavobacterium* sp. GS13 NZ CP037933 (76.82% AAI)
- *Flavobacterium gilvum* NZ CP017479$^{T}$ (72.75% AAI)

According to MiGA, the sequence most likely belongs to the family *Flavobacteriaceae* (p-value: 0), probably belongs to the genus *Flavobacterium* (p-value: 0.078), and possibly even belongs to the same subspecies of *Flavobacterium* sp. GS13 NZ CP037933 (p-value: 0.33). These data are presented in Supplementary Table 9 below. The results corroborate our independent ANI results presented above.

**Supplementary Table 9:** Average amino acid identity and genome relatedness: Average sequence identities to reference datasets in the database, as calculated by MiGA. Only the top-6 values are displayed.

| Dataset | AAI (%) | Std Dev (AAI%) | Fraction of proteins shared (%) |
|---|---|---|---|
| *Flavobacterium* sp. GS13 NZ CP037933 | 76.82 | 18.58 | 52.26 |
| *Flavobacterium gilvum* NZ CP017479 | 72.75 | 18.14 | 51.46 |
| *Flavobacterium commune* NZ CP017774 | 71.93 | 18.64 | 53.29 |
| *Flavobacterium crassostreae* NZ CP017688 | 71.69 | 17.86 | 60.32 |
| *Flavobacterium sp.* HYN0086 NZ CP030261 | 70.87 | 18.78 | 46.86 |
| *Flavobacterium faecale* NZ CP020918 | 69.43 | 0 | (estimated) |

***MyTaxa scan***

A MyTaxa scan [42] was also conducted within the MiGA framework. MyTaxa scans use each individual gene within an unknown sequence as a classifier, allowing evidence for taxonomic relationships to be gained from a variety of locations on the genome, which can be useful in cases of HGT. MyTaxa also weights each gene based on “its (predetermined) classifying power at a given taxonomic level and frequency of horizontal gene transfer”. This combination of evidence can allow stronger inference of identity to be gained in marginal cases. In our scaffold, however, it is clear that the best matches for the majority of genes are *Flavobacterium* sequences. These can be seen in Supplementary Figure 11 below, where *Flavobacterium* matches are shown in light blue.

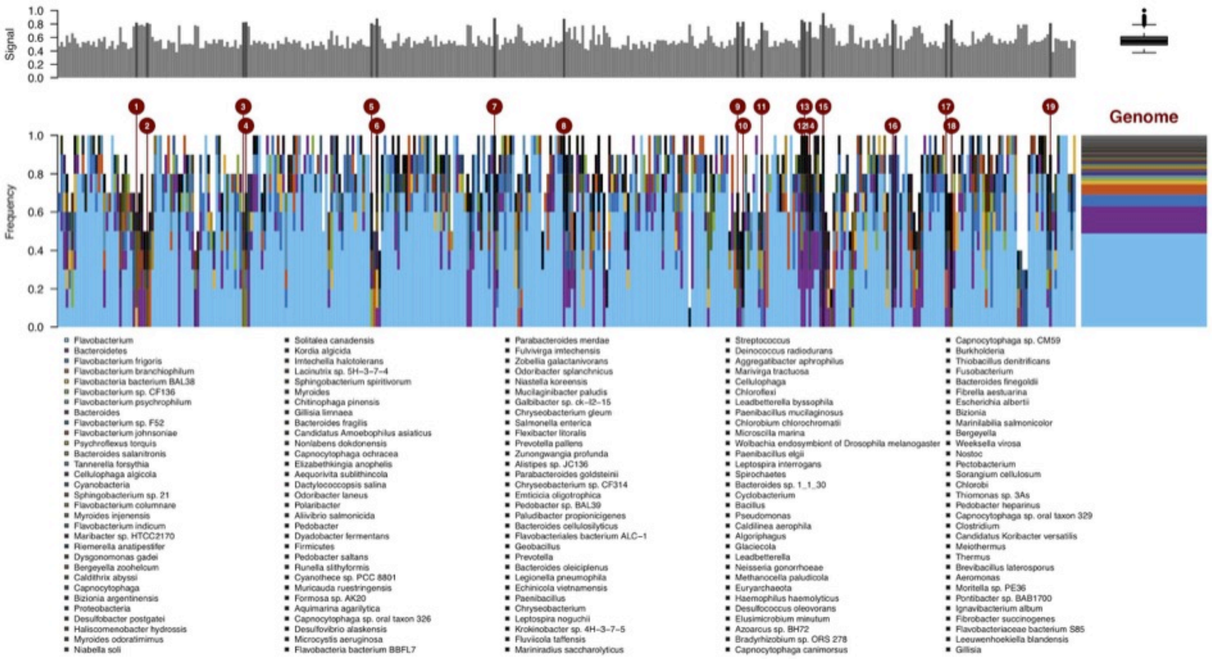

**Supplementary Figure 11:** MyTaxaScan output. Light blue represents regions that matched *Flavobacterium* sequences in this analysis. For box and whisker plot top right, boxplot centre line is median of signal plot top left, box limits are quartiles 1 (Q1) and 3 (Q3), whiskers are 1.5× interquartile range and points are outliers. The 3,811 genes on the identified *Flavobacterium* sp. scaffold were used as input for this signal plot.

Using the genes catalogued by MiGA and MyTaxa scan, we were able to extract metrics related to gene set quality and recovery. These metrics were excellent. Of the *essential* gene complement listed in MiGA, we found: 90/111. Our completeness metric is 81.1%, which is "very high", and our contamination metric was calculated to be 3.6% (very low). The overall quality of our assembly is given as 63.1% (high). This assembly will therefore be a useful dataset if this bacterium is found regularly across the *Ephydatia muelleri* radiation, with all the data readily available to use *Flavobacterium* sp. as a genomic model for the study of symbiosis in this species.

**16S rRNA gene cladogram**

A cladogram (Supplementary Figure 12) was generated using 16S sequences of *Flavobacterium* and related taxa (1300-1510bp in size) downloaded from NCBI, with a total of 225 sequences. Reads were aligned with MAFFT v7.407[43] using default options. The alignment was trimmed with TrimAL v1.4 (automated mode) [44]. IQ-Tree v1.6.10 [45] was used to generate the cladogram. The model of substitution was selected automatically by IQ-Tree, and SYM+I+G4 was chosen. Analysis was done using non-parametric bootstraps (1000 replicates), and is displayed below (Supplementary Figure 12), with the place of sampling (freshwater, soil or seawater) indicated with colours as noted on the figure legend. Our sequence was placed in a highly nested internal node, and is the 6th taxon from the lower edge of the tree displayed below. It is shown in red to aid viewing, but may require enlargement to visualise (or can be viewed in high quality in the high quality figure files downloadable from Ephybase, https://spaces.facsci.ualberta.ca/ephybase/).

On the basis of ANI, MyTaxa scan and phylogenetic evidence, we are very confident of the assignation of this bacterium into the *Flavobacterium* genus. It also seems highly likely to be a novel species, although it is related to some previously observed and sequenced organisms. The near-complete and high quality genome seen here will be useful for further investigations of the potentially symbiotic role of this bacterium with *E. muelleri.*

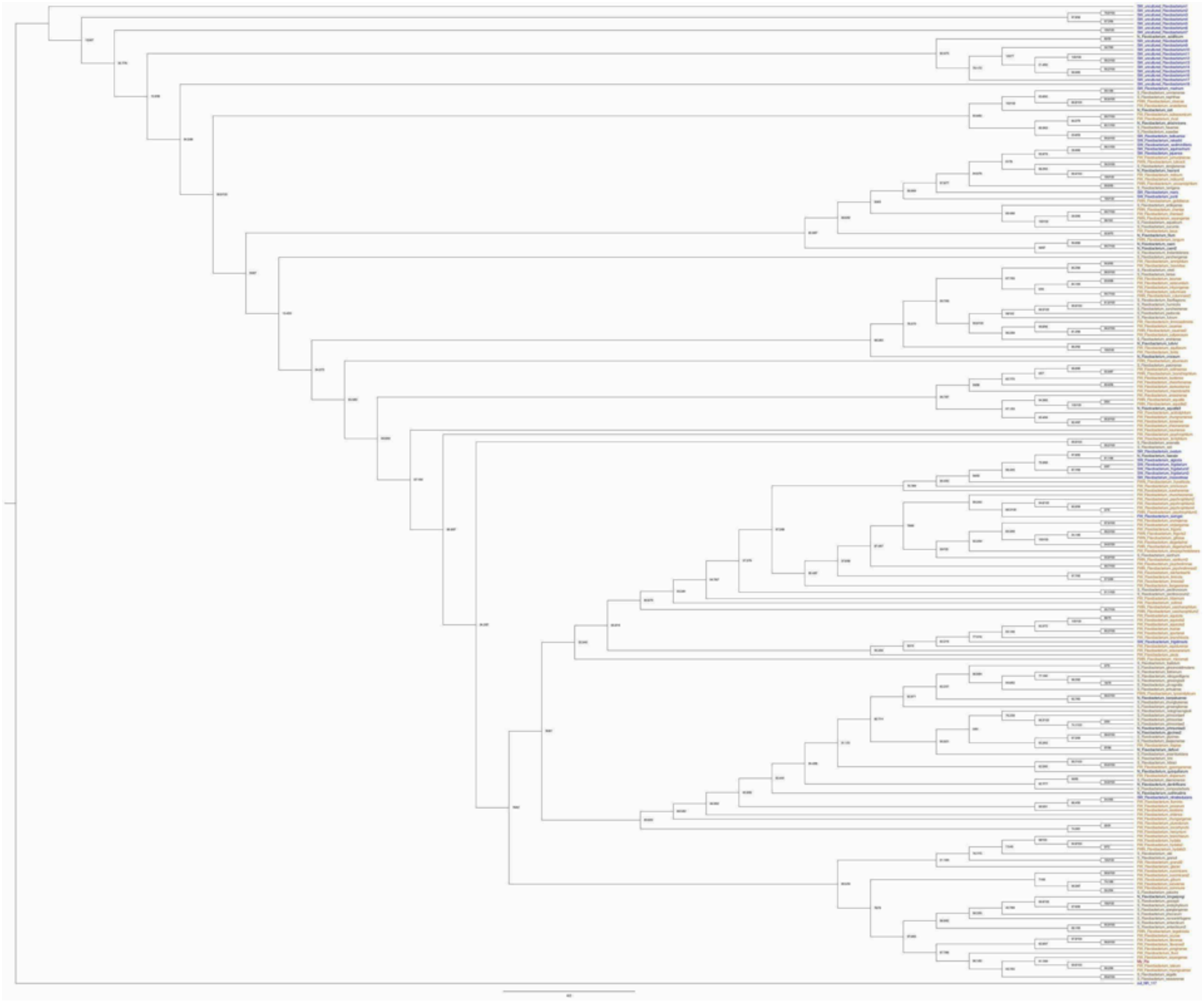

**Supplementary Figure 12:** 16S rRNA gene cladogram of *Flavobacterium* sp. (our bacterial sequence) and related species, recovered using IQ-Tree and the SYM+I+G4 model. Numbers at base of nodes are proportions of 1,000 bootstrap replicates. Samples starting with FW are freshwater (shown in orange), starting with S were recovered from soil (green), SW from seawater (blue) and N (black) have no accession information. *Flavobacterium* sp. (our bacterial sequence) is named My_Fla (red), and is found 6th from the bottom of the phylogeny at lower right. This figure is included in high resolution in Ephybase, https://spaces.facsci.ualberta.ca/ephybase/.

### 4.2 Genomic Islands

IslandViewer4 [46] was used to scan the *Flavobacterium* sp. genome and look for genomic islands. Nineteen putative genomic islands were found (Supplementary Figure 13 below). While they are generally distributed throughout the genome of this species, there are some small clusters of putative island sequences picked up by our analysis. These could represent areas of the genome where insertion is more straightforward and has happened repeatedly, or the remains of insertion events which have been fragmented by evolutionary processes in the genome.

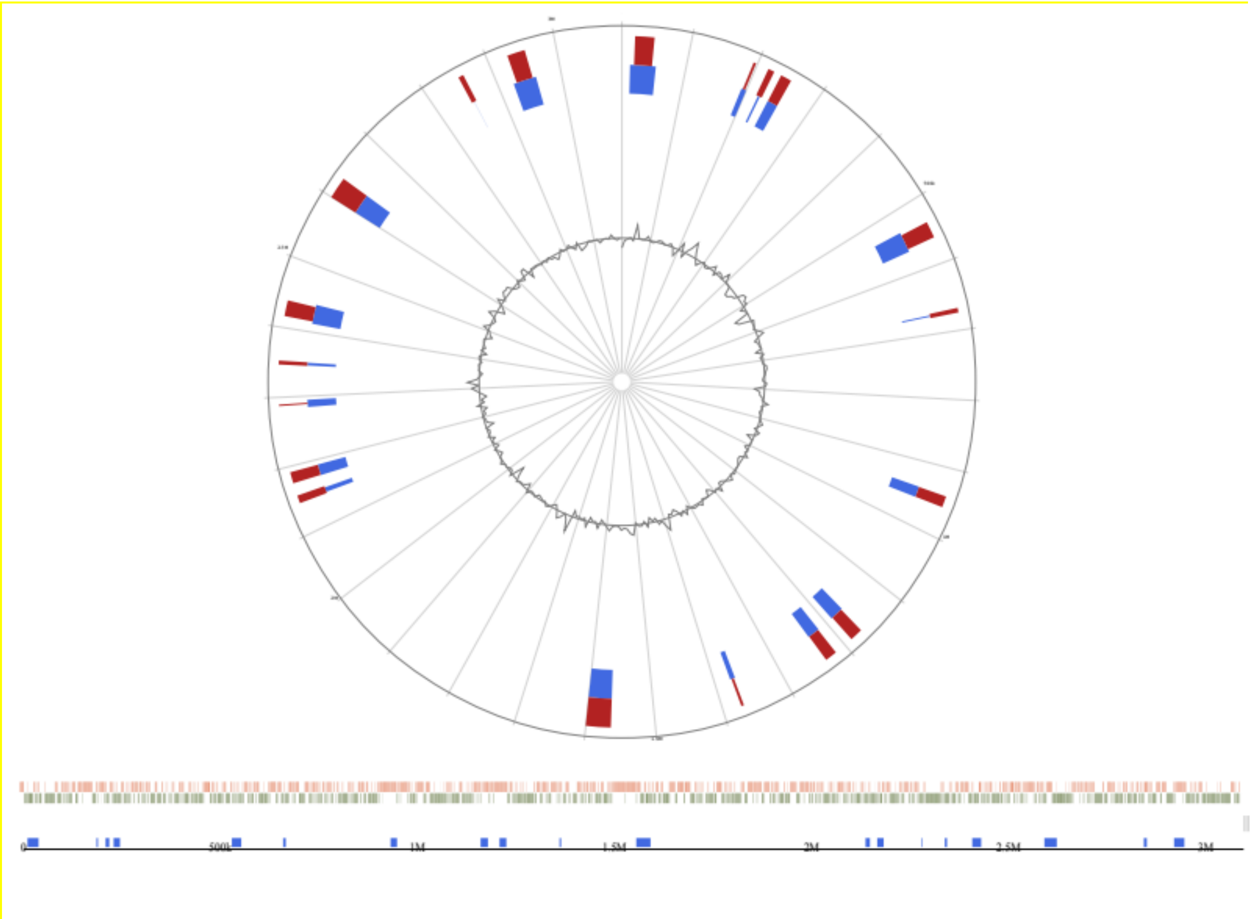

**Supplementary Figure 13:** IslandViewer4 output, showing the results of prediction of regions of possible origin by horizontal transfer in the *Flavobacterium* sp. sequence found in the *Ephydatia muelleri* assembly.

#### 4.3 Secondary metabolites

Secondary metabolites can be key components of symbiotic relationships between microorganisms and their hosts. They can also impact hosts negatively, and be informative regarding the taxonomy of, and environments inhabited by, micro-organism species, providing vital clues as to their biology  To search for secondary metabolites antiSMASH v5.0 [47] was run. Two hits were found.

- a terpene cluster, at Location: 1,304,271 - 1,325,110 nt. (total: 20,840 nt). This was most similar to a carotenoid (42% similarity).
- A *Type-III Polyketide Synthase* cluster (incomplete). Location: 1,978,058 - 2,019,092 nt. (total: 41,035 nt)

These clusters represent groups of genes that could aid in the production of specialised biomolecules, although whether this benefits *E. muelleri* is unclear. The arrangement of genes within these clusters closely matches the arrangement of these gene complexes in related *Flavobacterium* sequences (Supplementary Figures 14, 15 below).

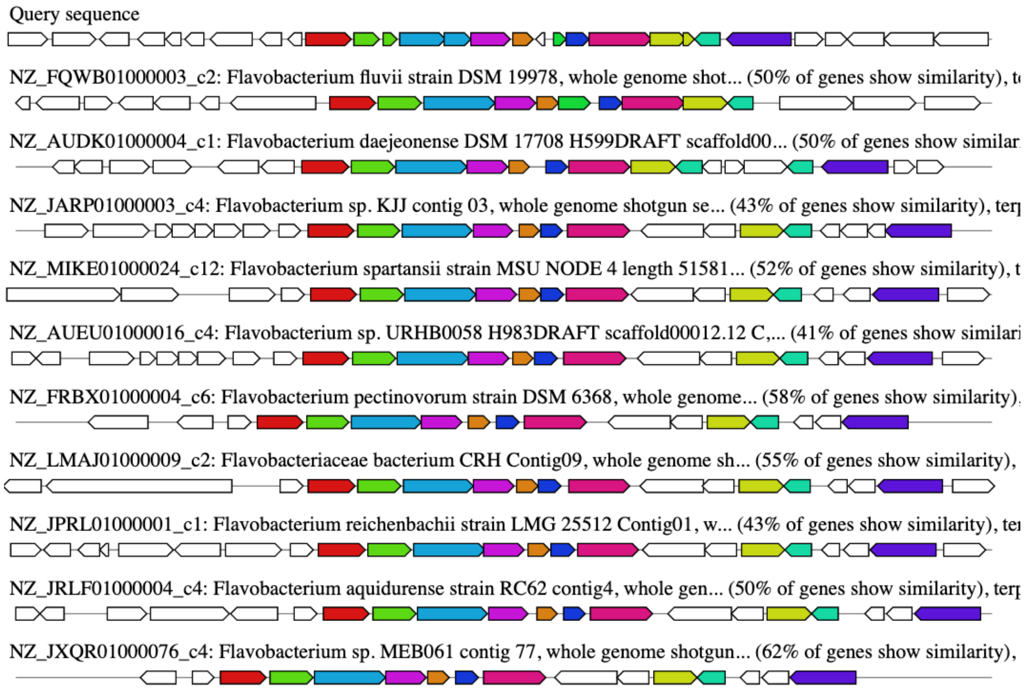

**Supplementary Figure 14**: Terpene-related gene cluster in our species (top, query sequence) and related species (below). Genes with the same colour between species are inferred to be homologous by antiSMASH.

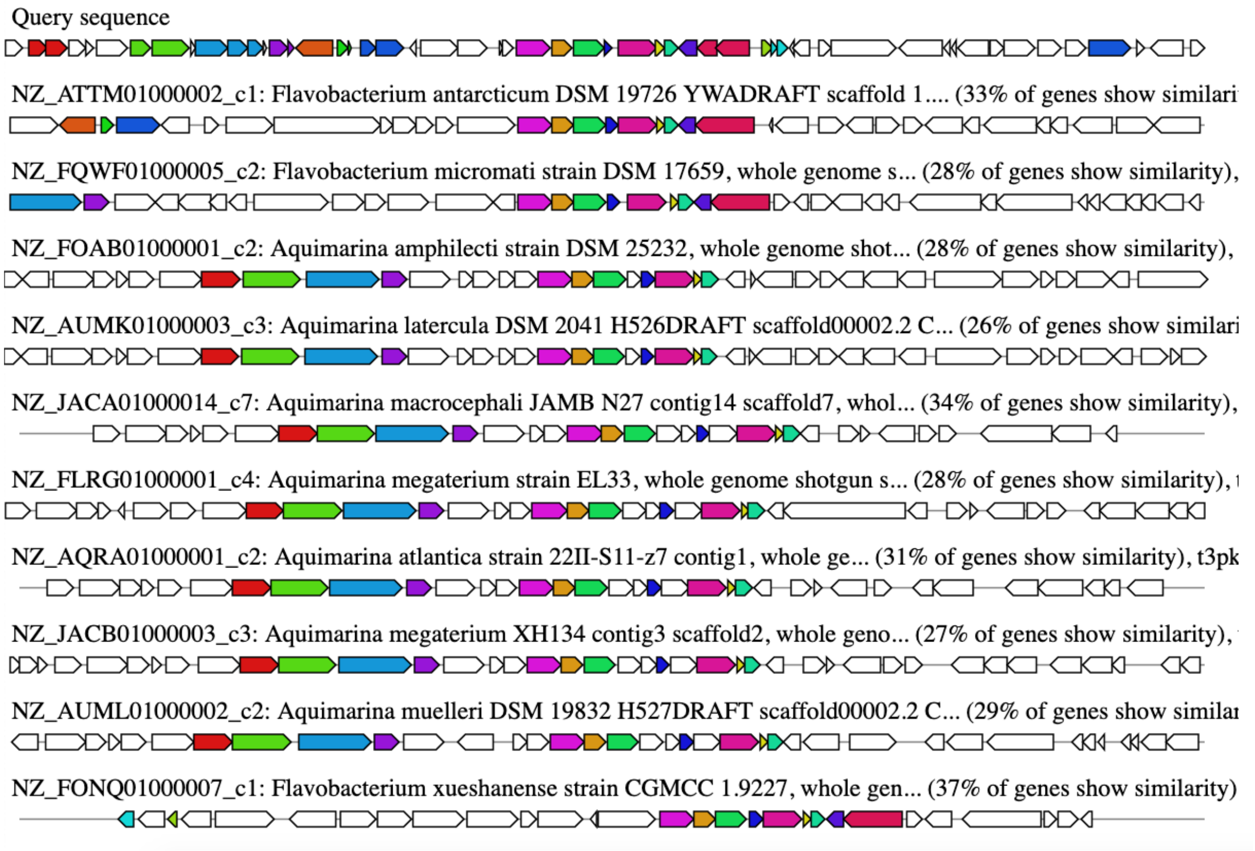

**Supplementary Figure 15:** Type-III Polyketide Synthase gene cluster in our species (top, query sequence) and related species (below). Genes indicated with the same colour between species are inferred to be homologous by antiSMASH.

### Supplementary Note 5: Gene models and annotation

#### 5.1 Gene prediction and metrics

The gene predictions used as our final gene set were generated using AUGUSTUS 3.3.2 [48] on the online tool available at http://bioinf.uni-greifswald.de/webaugustus/. Both training and annotation was performed using this tool, with a *de novo* assembly of the *Ephydatia muelleri* transcriptome used for training [27]. The following settings were allowed for prediction: UTRs allowed, both stranded reporting, no alternative transcripts, whole genes only (but note that UTR prediction in non-model organisms is difficult, and few UTRs were predicted for our dataset).

The results of this prediction were the basis for all further analyses performed here. A total of 39,245 genes were predicted (after the removal of 84 genes found on scaffold #25, Scaffold_153_HRSCAF_192, which is noted as a bacterial sequence, see Supplementary Note 4). Of these, 38,962 had canonical start codons, 38,247 canonical stop codons and 37,974 both start and stop codons. The usage of stop codons was not uniform - 10,904 gene models ended in TAA, 10,136 in TAG, and the plurality, 17,207, used TGA. This is consistent with similar ratios seen in many eukaryotic species [30] although some invertebrates favour TAA.

To ensure these predictions were robust, we also assayed a number of alternative gene prediction strategies. We annotated a masked version of the genome, based on the output of RepeatModeler/Masker (Supplementary Note 3.3). This is not recommended for complete gene predictions, as it often truncates complete gene models, but gives an overview of the number of genes that may have been inserted by repetitive elements into the genome, and the number of genes found outside repetitive regions (which can be silenced by epigenetic mechanisms). From this, 23,696 genes were annotated, a subset of the complete prediction, and these can be downloaded from our bitbucket site (AlternativeInferiorGenePredictions.tar.gz, https://bitbucket.org/EphydatiaGenome/ephydatiagenome/downloads/AlternativeInferiorGenePredictions.tar.gz).

We repeated our use of AUGUSTUS, using the gene sets of a diverse range of other metazoans to make gene predictions and test them against our annotation. We used AUGUSTUS-3.3.2 locally, trained AUGUSTUS with hints, and sequentially used the eukaryotic species present in the AUGUSTUS training set (config/species/ within the AUGUSTUS folder) to generate gene models (settings: --UTR = on --extrinsicCfgFile = extrinsic.M.RM.E.W.cfg --alternatives-from-evidence = true --allow_hinted_splicesites = atac). In no case was the recovery of BUSCOs superior to our own gene set, with generally fragmented results (e.g., the most complete metazoan recovery was gained with the pea aphid *Acyrthosiphon pisum*, with the following metrics: C:51.6%[S:41.5%,D:10.1%], F:19.4%, M:29.0%, n:978). We therefore did not proceed further with these gene sets, although they may contain genes not annotated by our approach and are available from our bitbucket site (AlternativeInferiorGenePredictions.tar.gz, https://bitbucket.org/EphydatiaGenome/ephydatiagenome/downloads/AlternativeInferiorGenePredictions.tar.gz).

MAKER [49] was also tried as an alternative annotation pathway. This resulted in a total of 36,520 gene models, with marginally improved BUSCO results (207 Complete BUSCOs, (167 single-copy, 40 duplicated), 60 fragmented BUSCOs, 36 missing BUSCOs, of 303 total BUSCO groups searched) compared to our automated AUGUSTUS results (88.1% vs 80.20% recovery). However, these genes were markedly truncated compared to the results of our automated annotation, with many missing several complete exons (median length = 208 amino acids, vs 348 in online AUGUSTUS-derived dataset, mean 307.74 vs 497.66 amino acids). Combined with the smaller total number of gene models in this resource, we decided against the use of the MAKER set in our further work, but it is also available to interested researchers.

**Supplementary Table 10:** Statistics, Final gene set used for further investigation, alongside those of previously published resources.

| | *Ephydatia muelleri* | *Amphimedon queenslandica* (V2.1) |
|---|---|---|
| Number of gene models: | 39,245 | 43,615 |
| Mean gene length (aa residues) | 497.66 | 337.84 |
| Maximum gene length (aa residues) | 37,921 | 18,893 |
| Minimum gene length (aa residues) | 10 | 5 |
| Mean number of introns per gene: | 3.009 | 2.01 |
| Mean intron size: (bp) | 361.49 | 327.62 |
| Median intron size: (bp) | 138 | 70 |
| Mean gene size: (bp) | 4507.53 | 2425.68 |
| Mean inter-gene distance: (bp) | 3524.79 | 2145.72 |

To determine whether genes could be missing from this figure, we compared our predicted gene models with the deep *E. muelleri* transcriptome used for gene prediction, and taking into account whether orthologous sequences were also found in the genomes of other sponge species (see Supplementary Note 7 for further details). A total of 1393 total orthogroups were recovered in our transcriptome that were not seen in our genome, and potentially shared with other sponge species. However, this total could include shared contamination or chimeric sequences not included in the *E. muelleri* genome, as well as sequences present in the genome but not mapped to a complete gene model. This therefore gives a likely upper estimate for missing genes not found in the gene model complement of around 3.5% (1393/39,245) although the true figure is likely slightly less than this quantity.

It is true that 39,245 genes is a higher number than that found in most metazoan genomes, but is similar to that predicted for *Amphimedon queenslandica* (30,327 [35], revised to 40,122 in [50], and 43,615 in the current V2.1 build on Ensembl). Intron and gene size were calculated using gtfstats.py [38]. The average intergene distance was calculated as 3524.79 bp. Average intron, exon and intergenic region length were highly similar in both sponges (*A. queenslandica* and *E. muelleri*), although exons were more numerous in *E. muelleri* and introns in *A. queenslandica* (Supplementary Table 10, Supplementary Figure 16). The number of intergenic regions in both sponges is far more abundant than in the rest of the metazoans we used for comparison (Supplementary Figure 16) although this will be a direct consequence of the larger number of genes seen in sponges.

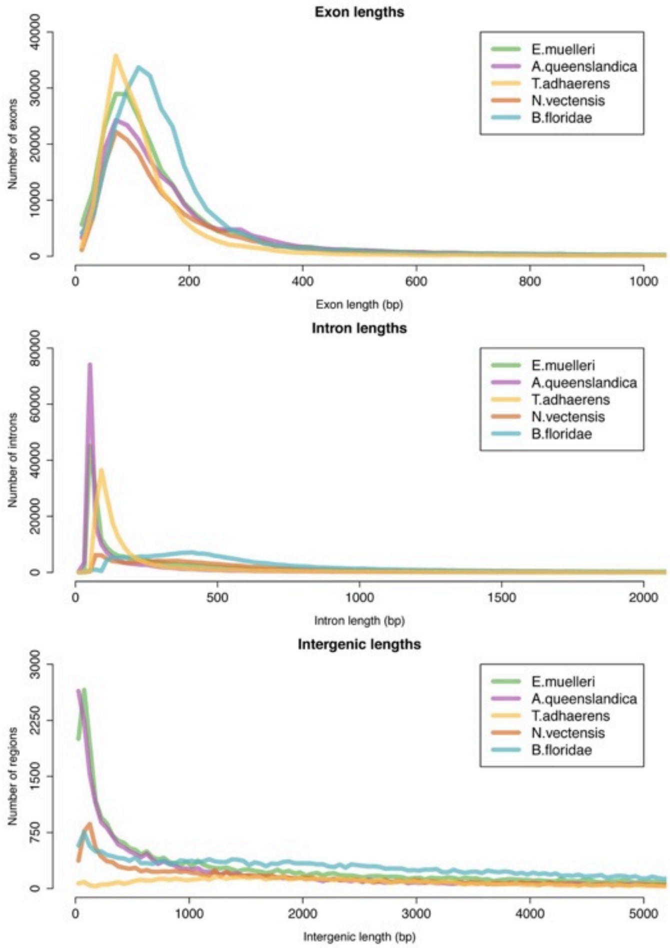

**Supplementary Figure 16.** Histograms of intron, exon and intergenic region numbers and sizes in *Ephydatia muelleri* and other animal taxa for comparison.

#### 5.2 Automated annotation results

We obtained the initial annotations for our gene models using BLAST 2.10.0 [51] and DIAMOND 0.9.31 [52] against two different databases: NCBI *nr* and Swiss-Prot (last accessed in September 2019), reporting the best hit with an *E*-value threshold of $10^{-5}$ in both cases. Our blast IDs obtained against these two different databases, resulted in 29,171 of our gene models with a hit against nr, and 19,736 with a hit against swissprot. Of the 29,571 genes with a blast hit against nr, 13,579 blasted against sponge sequences: 12,939 against *Amphimedon queenslandica* sequences, 330 against the freshwater sponge *Lubomirskia baikalensis*, while only 286 against *Ephydatia* species contained in nr (Supplementary Figure 17A). In addition, we obtained 79 hits against different *Flavobacterium* species. The results against Swiss-Prot show that only 19,736 obtained a blast hit (mostly to human sequences).

#### 5.3 Functional annotation

We annotated our dataset further by comparison to functional databases, using KEGG [53] and BLAST2GO PRO [54] to obtain the gene ontology (GO) terms associated with the blast hits obtained against swissprot for Biological Process, Molecular Function and Cellular Component, with the GOSlim function. KEGG annotation was carried out using the KEGG KAAS server [55]. The bidirectional best hit (BBH) method was used, with a BLAST bit score threshold of 30. The eukaryotic organism list, with the addition of placozoan, poriferan and cnidarian genes, was used as the comparison dataset. Of the 39,329 *E. muelleri* protein sequences, 13,166 were mapped to pathways in the KEGG database. These results can be found in Supplementary Data 2.

KEGG pathways were well recovered by this analysis, indicating that our dataset contains the expected gene complement of a eukaryotic organism. By way of example, the Glycolysis / Gluconeogenesis pathway has 45 proteins mapped to it, and is missing a maximum of 15 genes, although not all genes are expected to be found in every species. The reference KEGG map for *Amphimedon queenslandica*, by way of comparison, is missing 27 genes from this pathway. Similarly, 31 genes are recovered in the Basal transcription factors category, with only 4 genes missing (*TFIIB, TAF13, TAF14* and *CCNH*). The reference resource for *Amphimedon queenslandica* is also missing 4 of these genes (overlap: *TAF14*). These KEGG maps will therefore be useful for discerning how these pathways have changed in the course of poriferan evolution, and as an alternative reference to *A. queenslandica,* for making inferences about gene pathway content in sponges.

Our BLAST2GO results also show relatively high functional annotation recovery for the *Ephydatia muelleri* genome, with 19,362 genes recovering a functional annotation when using the swissprot database (release release-2019_08) for blastx, and 11,421 genes when using nr (2019-10-21) as the reference database for blastx. As can be expected, the annotation obtained from swissprot is much deeper than that obtained with nr (Supplementary Figure 17 B,C). Annotation with swissprot delivers a deep and meaningful GO annotation for many genes.  Full BLAST2GO results in Supplementary Data 2.

**Supplementary Figure 17:** A. The percentage of sequences assigned to different taxonomic groups using the nr database, with additional detail on Porifera shown in the inset at right. B. Score distribution of GO terms in Biological process using the nr database. C. Score distribution of GO terms in Biological process using the Swiss-Prot database.

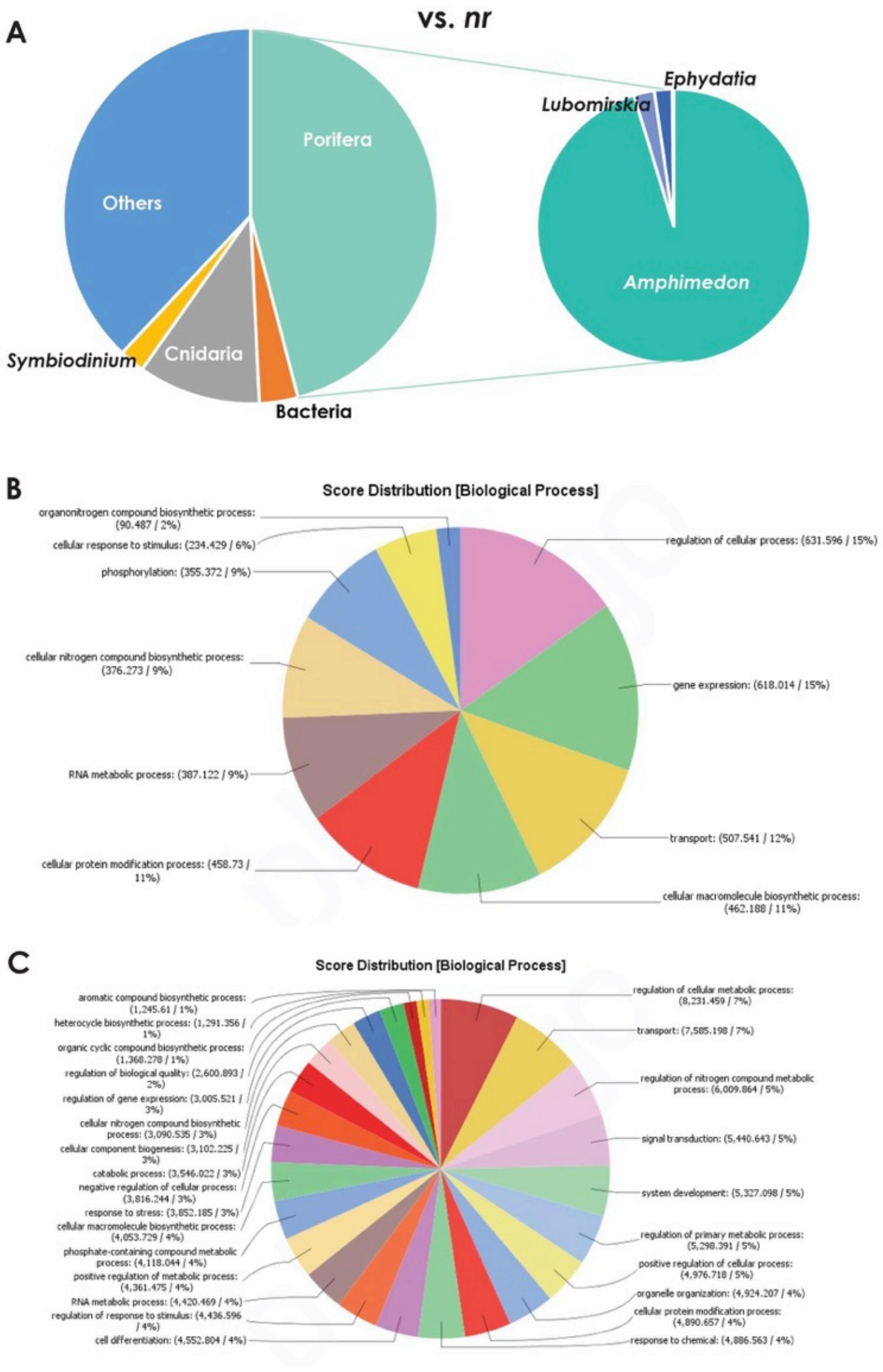

### Supplementary Note 6: Genome Architecture and Insights into Regulatory Structures

#### 6.1 Insights into longer-range gene regulation and chromatin interaction

The HiC data sequenced as part of genome assembly efforts also gave us the opportunity to visualise structural features of the genome, especially topologically associating domains (TADs) and loops. Chromatin loops are comprised of two regions of the genome that interact at a high frequency, while TADs are larger areas of the genome that show more interactions with one another than found in surrounding regions, or across the genome on average. Loops appear as dots on a HiC contact map, while TADs appear on these plots as darker shaded triangular regions, although these can be difficult to discern from background without highlighting.

Both TADs and loops were identified using Bowtie2 [56] and HOMER v4.11 [57]. Bowtie2 was used to index all scaffolds longer than 1 Mb in length. homerTools was used to trim reads (MboI/DpnII (GATC) sequence removed, with -mis 0 -matchStart 20 -min 20 settings). Bowtie2 was then used to map all HiC reads to the genome index. homerTools was used to create a tag directory and juicer output files with makeTagDirectory (-tbp 1, to remove PCR duplicates) and tagDir2hicFile.pl (-juicer auto, -p 20). analyzeHiC was used to visualise HiC maps, and the findTADsAndLoops.pl script (-res 3000 -window 15000) to find TAD and loop regions. This works by generating relative contact matrices for each scaffold, and then searching for areas with a significantly higher level of interactions relative to the surrounding region.

The average TAD FPKM coverage was 3.67 (across 1042 regions total) and the average loop anchor FPKM coverage was 3.34 (across 2754 regions total). HOMER assigned 492/1042 (47.217%) of the TADs as *good*, with 550/1042 (52.783%) excluded for insufficient coverage. Similarly, 1153/1378 (83.672%) loops were retained, with insufficient coverage excluding = 225/1378 (16.328%) of these. We were therefore able to note the presence of 492 TADs and 1153 loops in this genome. Supplementary Figure 18 below shows an example of the presence and absence of these features across the entirety of Scaffold 1 (Chromosome 1), from which the figure in the main text has been taken. TADs and loops do not always co-occur in our data, but the presence of loops in areas free of TADs may indicate that further TADs could be revealed with additional sequencing depth.

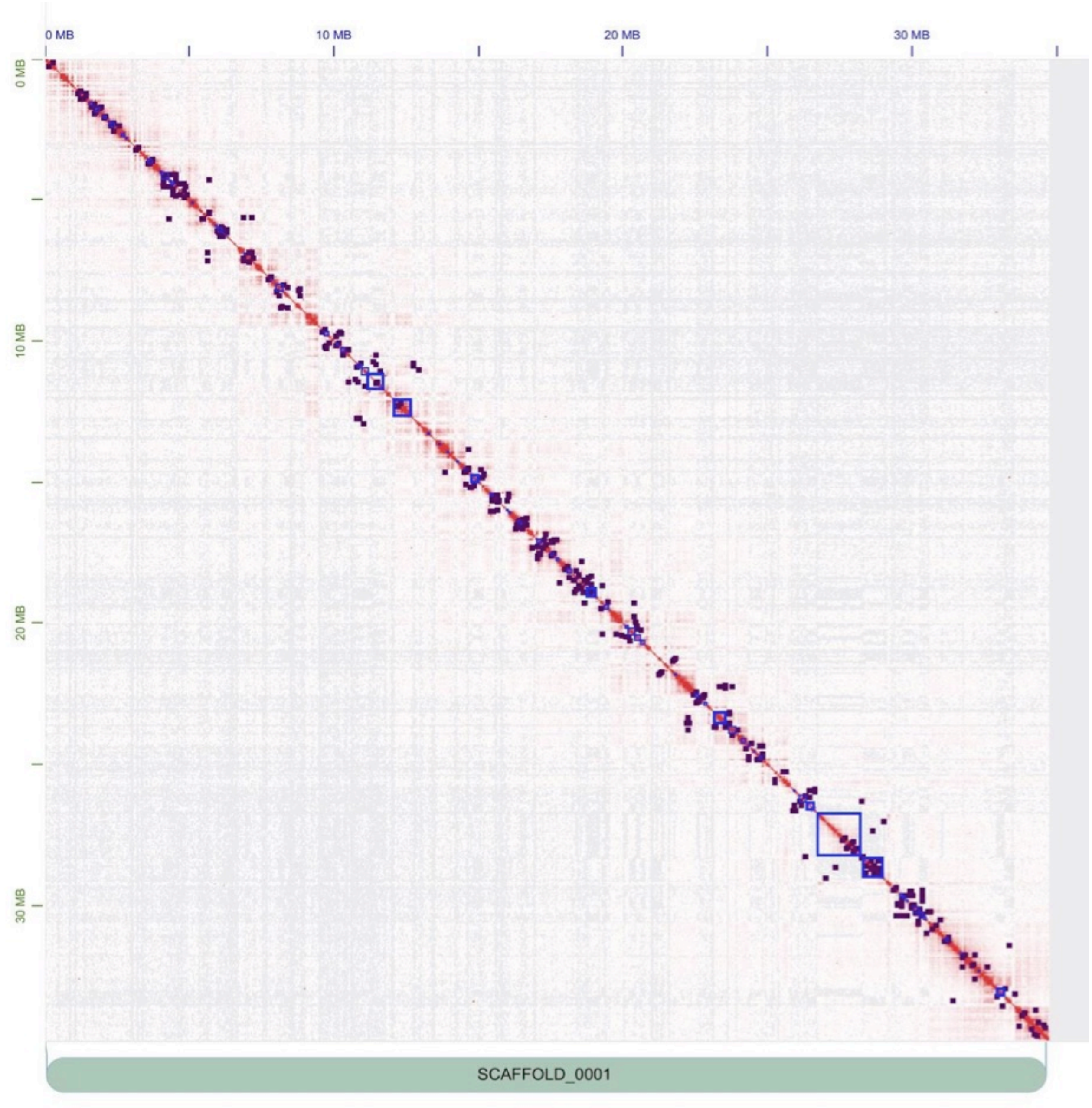

**Supplementary Figure 18:** Distribution of topologically associating domains (TADs) and chromatin loops on Scaffold (=Chromosome) 1 of the *Ephydatia muelleri* genome assembly. TADs and loops both identified using HOMER [57]. TADs indicated with blue squares. Loops indicated with purple dots. Red indicates HiC contacts and quantity.

### 6.2 Cytosine DNA methylation

Whole Genome Bisulfite Sequencing was performed on tissue from a fully developed freshwater sponge (stage 5) *E. muelleri* genomic DNA sample. We used the MethylC-seq protocol [58], combining 300 ng of *E. muelleri* DNA with 1 ng of spiked in unmethylated Lambda genomic DNA as a control to determine the bisulfite non-conversion rate (1.9%, false positives), and the sequencing was performed on an Illumina NovaSeq 6000 (37.5 million 50 bp paired end reads). We mapped the bisulfite converted reads using BS-Seeker2 [59].

Vertebrate and *A. queenslandica* genomes are highly depleted for CpG dinucleotides, which is attributed to methylation tending to cause deamination resulting in detrimental mutations [60].We then tested whether *E. muelleri* shows a biased ratio of CpGs versus the expected abundance (1 = equilibrium, <1 CpG loss) and we observed a less biased genome in *E. muelleri* than in *A. queenslandica* (Supplementary Figure 19A) as described in the main text. Supplementary Figure 19B shows that the mCG/CG ratio surrounding highly expressed genes is lower on average around the transcriptional start site, although it rises to above normal levels further from this point. Supplementary Figure 19C shows methylation signal across protein coding gene bodies and transposable elements, as revealed by Whole Genome Bisulfite Sequencing, can be seen.

The *E. muelleri* genome encodes two paralogs of DNMT1 and one of DNMT3, which are largely conserved with those of other sponges and animals. Interestingly, we found that the PWWP domain in the *E. muelleri* DNMT3 orthologue lacks a unique insertion, found in *A. queenslandica* (Supplementary Fig 19F), which suggests that the insertion is a secondary state not common to demosponges. Given that the PWWP domain is known to influence the targeting of DNMT3, it is possible that this innovation contributed to the hypermethylated state of the *A. queenslandica* genome. However, *S. ciliatum* also lacks the PWWP insertion and has higher methylation levels than *E. muelleri*; therefore, as discussed previously [60], we believe that hypermethylation is not the by-product of a methyltransferase change, but a long-term evolutionary process.

**Supplementary Figure 19 (Overleaf): Cytosine methylation and CpG composition of *E. muelleri* genome.** A) Proportion of CpG dinucleotides in the genome assemblies of various sponges, normalised by the expected number of CpGs (given the GC %). B) Mean CpG methylation level on gene bodies classified as per transcriptional level. 1-10th decile are the top expressed genes, whereas non expressed are defined as having <1 Reads Per Kilobase of transcript per Million (RPKM). C) Heatmap showing methylation levels on protein coding gene bodies and transposable elements (RepeatModeler insertions spanning > 200 bp), classified by their relative position regarding a protein coding gene body. Only genes and transposons where Whole Genome Bisulfite Sequencing coverage was higher than 4X are represented to avoid excess of missing data. TSS: Transcriptional Start Sites; TES: Transcriptional End Site; TE: Transposable element. D) Distribution of methylation levels on transposable elements found in intergenic and genic regions, classified as per RepeatModeller annotations. Many LTR retrotransposons show similarly high methylation levels irrespective of the genomic position, whereas most DNA transposons show lower methylation levels throughout and more acute differences. Boxplot centre lines are medians, box limits are quartiles 1 (Q1) and 3 (Q3), whiskers are 1.5× interquartile range and points are outliers. For numbers of transcriptional elements, see Supplementary Table 6. 37.5 million 50 bp paired end reads from a single sample were mapped. E) Methylation level distribution linked to transposable element divergence. Kimura divergence for each transposable element family was obtained by using the calcDivergenceFromAlign script in RepeatMasker. The total distribution of divergences was then subdivided in 10 deciles, which are ordered from less divergent to more divergent. Boxplot centre lines are medians, box limits are quartiles 1 (Q1) and 3 (Q3), whiskers are 1.5× interquartile range and points are outliers. For numbers of transcriptional elements, see Supplementary Table 6. 37.5 million 50 bp paired end reads from a single sample were mapped, F) Amino acid multisequence alignment representing the PWWP domain from DNMT3 orthologues. *E. muelleri* DNMT3 corresponds to Em0006g900a gene model, yet it is a truncated gene model within an assembly gap, therefore we obtained the full-length protein from a Trinity transcriptome assembly. Proteins were aligned using MAFFT.

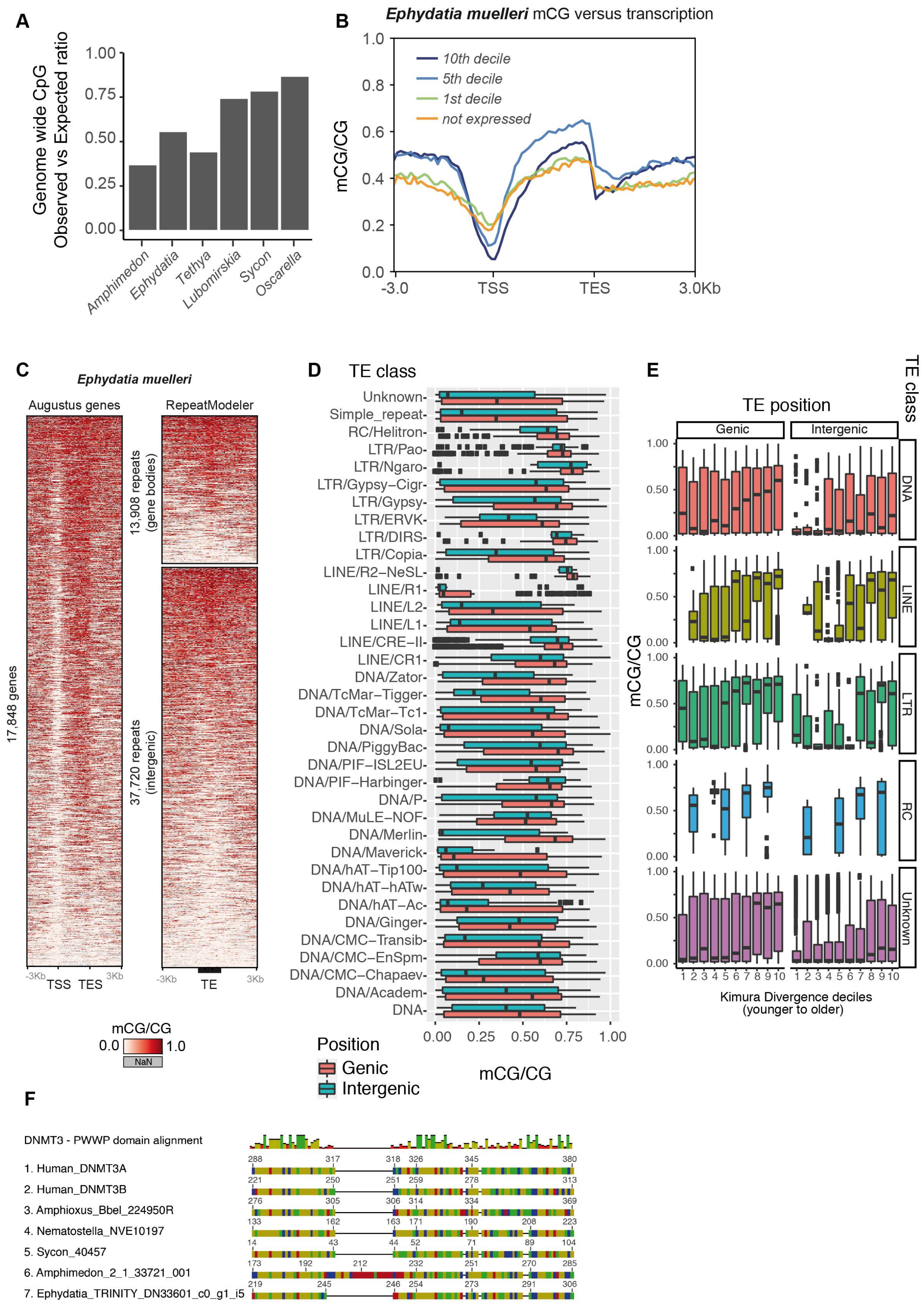
A
Genome wide CpG
Observed vs Expected ratio
1.00
0.75
0.50
0.25
0.00
Amphimedon
Ephydatia
Tethya
Lubomirskia
Sycon
Oscarella
B
Ephydatia muelleri mCG versus transcription
10th decile
5th decile
1st decile
not expressed
mCG/CG
1.0
0.8
0.6
0.4
0.2
0.0
-3.0
TSS
TES
3.0Kb
C
Ephydatia muelleri
Augustus genes
RepeatModeler
13,908 repeats
(gene bodies)
37,720 repeats
(intergenic)
17,848 genes
-3Kb
TSS
TES
3Kb
-3Kb
TE
3Kb
mCG/CG
0.0
1.0
NaN
D
TE class
Unknown-
Simple_repeat-
RC/Helitron-
LTR/Pao-
LTR/Ngaro-
LTR/Gypsy–Cigr-
LTR/Gypsy-
LTR/ERVK-
LTR/DIRS-
LTR/Copia-
LINE/R2–NeSL-
LINE/R1-
LINE/L2-
LINE/L1-
LINE/CRE–II-
LINE/CR1-
DNA/Zator-
DNA/TcMar–Tigger-
DNA/TcMar–Tc1-
DNA/Sola-
DNA/PiggyBac-
DNA/PIF–ISL2EU-
DNA/PIF–Harbinger-
DNA/P-
DNA/MuLE–NOF-
DNA/Merlin-
DNA/Maverick-
DNA/hAT–Tip100-
DNA/hAT–hATw-
DNA/hAT–Ac-
DNA/Ginger-
DNA/CMC–Transib-
DNA/CMC–EnSpm-
DNA/CMC–Chapaev-
DNA/Academ-
DNA-
0.00
0.25
0.50
0.75
1.00
mCG/CG
Position
Genic
Intergenic
E
TE position
TE class
Genic
Intergenic
DNA
LINE
LTR
RC
Unknown
mCG/CG
1.00
0.50
0.00
1 2 3 4 5 6 7 8 9 10
Kimura Divergence deciles
(younger to older)
F
DNMT3 - PWWP domain alignment
1. Human_DNMT3A
2. Human_DNMT3B
3. Amphioxus_Bbel_224950R
4. Nematostella_NVE10197
5. Sycon_40457
6. Amphimedon_2_1_33721_001
7. Ephydatia_TRINITY_DN33601_c0_g1_i5
288 317 318 326 345 380
221 250 251 259 278 313
276 305 306 314 334 369
133 162 163 171 190 208 223
14 43 44 52 71 89 104
173 192 212 232 251 270 285
219 245 246 254 273 291 306

### Supplementary Note 7: Analysis of *Ephydatia muelleri* novelties

Four sets of species were used for the analyses presented in this Supplementary Note:

- Supplementary Note 7.1: To determine patterns of gene gain and loss, in sponges compared to outgroups, we used only genomic datasets. These were derived from selected clades from across the metazoan tree of life, with 3 non-metazoan outgroups. (Total 13 species).
- Supplementary Note 7.2, 7.3: To assess what genes show signals of positive selection in *Ephydatia muelleri* we used previously published demosponge genomic (*T. wilhelma, A. queenslandica* and *X. testudinaria)* and transcriptomic (10 in total) resources and the *E. muelleri* protein set for orthogroup identification and curation. The above (demosponge-only) set was used to assess the gene complement of *E. muelleri,* especially in comparison to *T. wilhelma, A. queenslandica* and *X. testudinaria*. The sources of these datasets are listed in Supplementary Note 3.1.
- Supplementary Note 7.4 uses a larger dataset drawn from a previous study by Pett et al. 2019, to which the newly sequenced *E. muelleri* predicted proteome was added. The dataset included six Fungi, four Choanozoa, five Porifera, two Ctenophora and two Placozoa, three Cnidaria and 16 Bilateria. Proteome data obtained by whole genome prediction was used for all the species.This is the same dataset used for the orthofinder-based orthogroup phylogeny shown in Supplementary Note 1.3.
- Supplementary Note 7.5 concentrates only on clades where a freshwater adaptation event has taken place. Eight species from four phyla are represented, each with a freshwater and a marine representative.

In all cases except 7.1 (FastTree [61] used, with constrained trees as stated) and 7.4 (MCL [62] used for non-phylogeny analyses), the orthogroup matrices used for analysis were generated using Orthofinder2 [16], with IQ-TREE v1.6.12 [45], MAFFT 7.450 [43] and DIAMOND [52] blast options.

#### 7.1 Gains and losses

We compared the gene complements of sequenced sponge genomes with those of choanoflagellate outgroups, ctenophores, *Nematostella vectensis,* and several chordates, and checked for patterns of gene gain and loss, and how they differ across species and phyla. The results of this analysis can be seen in Supplementary Figure 20 below. The gains shown on this figure are total numbers, including paralogs, and are taken from the number of new genes determined from orthogroup trees by Orthofinder2. Losses shown are orthogroup counts, and were calculated using awk to determine the presence and/or absence of genes within orthogroups compared to presence within the ancestral state (from the Orthogroups.GeneCount.tsv).

These are presented first assuming sponges are the sister taxa to all other metazoans, and as a counter-example, with ctenophores shown as sister taxa. Full details of these results, and how they were determined, can be found in Supplementary Data 5.

*Gains in* E. muelleri *and beyond*

A strong pattern of gene gain is observed in all sponge species examined, with all species exhibiting an independent gain of over 12,000 genes regardless of assumed outgroup to the Metazoa. Only *Branchiostoma floridae* exhibits similar metrics, and in part this is due to the presence of incompletely curated gene models in the *B. floridae* v2 protein set (v1 filtered models fitted to the v2 genome) used for analysis. In general, other metazoan species show less than half the number of gene gains seen in the four sponge species examined here.

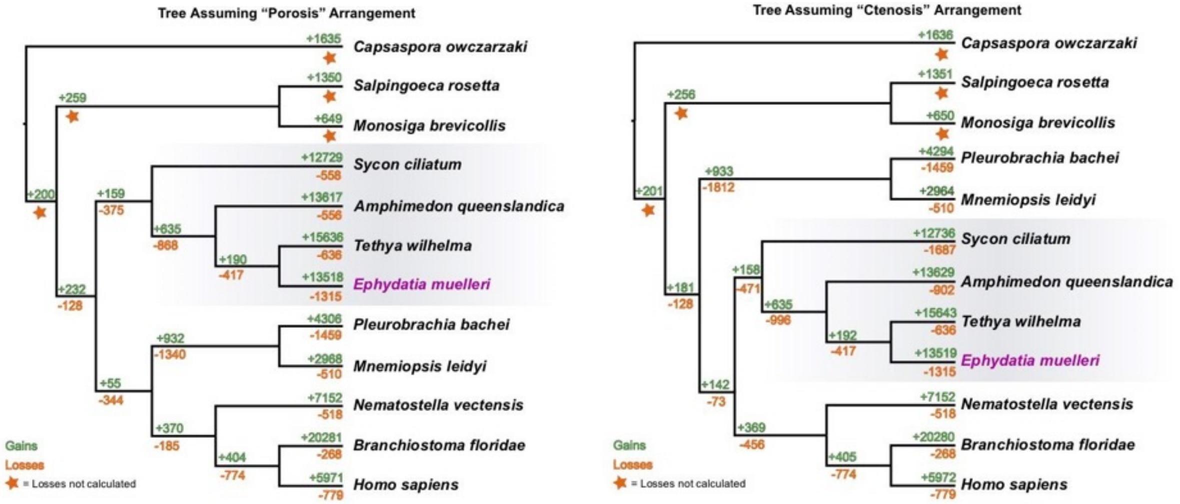

**Supplementary Figure 20:** Cladograms showing inter-relationships between a range of species used for analysis of gene gains through duplication, as well as losses when compared to ancestrally shared gene sets. Cladograms show results assuming sponge sister to all other metazoans at left (porosis), and ctenophores sister to other metazoans at right (ctenosis).

Sponges therefore seem to exhibit a uniformly strong tendency to generate extra gene copies, particularly by duplication, as will be predominantly measured by this analysis. *E. muelleri* shares this tendency to duplicate genes and this may represent a broadly shared adaptive trait in this phylum. The numbers of duplicated genes shared by multiple sponge species is, however, not markedly higher than observed at internal nodes in other parts of the phylogeny. These duplicated genes are therefore limited to individual species, and do not show any sign of being shared widely between sponges. The sponges sampled above are only distantly related, and further sampling of closer-related species may show patterns of retention. In Supplementary Figure 21 below, for example, deeper sampling of the Spongillida breaks down the patterns of orthogroup gain within that clade, and shows a large burst of duplications occurring at the base of the freshwater sponge radiation, and thus retained at least in that clade.

In short, sponges in general show a higher-than-background level of gene gain, as assessed here primarily by examining the addition of duplicated genes using Orthofinder2. *E. muelleri* is no exception to this rule, and, together with more nuanced sampling of Spongillida (Supplementary Note 7.2, Supplementary Figure 21), it is likely that gene gain plays an important role in the acquisition of novelty in the Porifera as a whole.

*Losses in* E. muelleri *and in comparison to other taxa*

Losses were only calculated for within the metazoan clade, given the necessity of a firm outgroup to discern these, and were calculated according to the tree shown in Supplementary Figure 20 above. A gene was only inferred as lost if it was present in at least two ancestrally located outgroup clades (e.g., a minimum of *C. owczarzaki* and one of (*S. rosetta* + *M. brevicollis*)), or in an ancestral clade and a sister clade to the sampled node, to exclude clade-specific novelties from consideration. All genomes examined exhibited some pattern of loss, either alone or shared ancestrally with other sequenced genome resources.

Within sponges, *E. muelleri* shows a higher level of terminal node loss, two-fold higher than any other sponge species examined. In independent tests not shown on the tree above, 1300 orthogroups were determined to be absent from the *E. muelleri* genome but present in at least one sponge and at least one non-metazoan (losses from the pre-metazoan ancestral cassette). An additional 15 orthogroups were noted as absent from the *E. muelleri* genome but present in at least one other sponge species and another metazoan (losses from the metazoan stem lineage set). These losses could be due to the changes necessary for adaptation to freshwater conditions, where many genes used for survival in marine environments may have been superseded by duplicate genes (see gains section above) or have proven broadly superfluous.

To an extent these absences from the genome are due to incomplete sampling. of these absences, 616 were independently found to be present in the *E. muelleri* transcriptome. This reduces the total gene loss in *E. muelleri* to only slightly higher loss than that observed in other sponges (699, cf 636, 558 and 556 in other sponges examined). In total, 1393 orthogroups (including the 616 above) were recovered in the transcriptome that were not seen in our genome. This genome resource is therefore missing some gene models, either through inadequate annotation or through incomplete recovery of the loci where these genes are found, and there is further discussion of this in Supplementary Note 5.1 above.

This genome will provide extra data for analysis of the ancestral sponge complement. In numerous previous studies, the complement of *A. queenslandica* has been taken as the representative of the sponge gene cassette. However, in our tests, there are 212 orthogroups lost in *A. queenslandica* that are found in our genome or that of another sponge species, which could falsely have been assumed absent from sponges in general based on that resource. Also, 53 orthogroups are present in *E. muelleri* and outgroup taxa, but absent in the other sponge species examined. These represent genes absent from previous genome sequencing projects, either through shared loss, lack of a gene model, or through non-recovery of the portion of the genome containing these genes. We also performed a similar analysis using the *E. muelleri* transcriptome and identified a further 18 genes matching this pattern. For inference of the ancestral condition in metazoans generally, or in sponges in particular, these "losses" can now be corroborated with an additional complete gene resource.

Large numbers of losses are also seen in ctenophores, and to a lesser extent in the human genome. A large number of orthogroups (1340) are absent from both sequenced ctenophore genomes examined under a constrained sponge outgroup to other metazoans, and an even higher number (1812) if ctenophores are specified as the outgroup. A similar number of orthogroups are also absent from the genome of *P. bachei* (1459). *M. leidyi* shows fewer losses (510 orthogroups). While high levels of loss in *P. bachei* could be down to incomplete genome sequencing in that species, the strong pattern of early, shared loss in the ctenophore lineage is less likely to be due to sampling error, and will reflect high rates of gene loss in this phylum.

Among the 616 genes lost in the genome of *E. muelleri* and also not recovered in the transcriptome, several were involved in epigenetic remodelling and gene regulation in other organisms, including *Histone acetyltransferase type B catalytic subunit, Histone deacetylase 11, Possible lysine-specific histone demethylase 1, ataxin-7 and -10, EEF1A lysine methyltransferase 1, Methyltransferase-like protein 13* and *9*, among others (full details, Supplementary Data 5). These losses are surprising, given the level of methylation found in this sponge, and point to interesting avenues of research. Interestingly, not all the orthologues related to those enzymes are lost, since *ataxin 2* and *3* were found in the genome, and other orthologues for the rest of enzymes mentioned as well. Equally intriguing are the losses of several *cyclins (D, H, and I)* and a *cyclin-kinase*, which are necessary to regulate cell proliferation. In the genome, we found over 20 *cyclin-kinases*, and 28 cyclins (*cyclin A*: 3, *cyclin B*: 11, *cyclin C*: 1, *cyclin D2*: 2, *cyclin E*: 2, *cyclin F*: 1, *cyclin G*: 1, *cyclin K*: 1, *cyclin J*: 2, *cyclin L1:* 1, *cyclin Q*: 1, and *cyclin Y:* 2).

In the ubiquitination machinery, several genes were also lost in *E. muelleri*: *E3 ubiquitin-protein ligase MGRN1, E3 ubiquitin-protein ligase RNF170, E3 ubiquitin-protein ligase Topors, E3 ubiquitin-protein transferase RMND5A, Probable E3 ubiquitin ligase complex SCF subunit sconB*, and several ubiquitin enzymes (full details, Supplementary Data 5). These losses might be related to other specific protein losses or divergent conformations. Perhaps more related to the adaptation to freshwater ecosystems, the loss of two vacuolar protein sorting-associated proteins and several solute carriers, which are related to adaptation and acclimation to freshwater systems [63,64],and also nucleoporins, which are important players in osmoregulation processes in vertebrates [65,66]. Changes in nucleoporins have been highlighted as important drivers of reproductive isolation [67], and here, the loss of some nucleoporin genes (*GLE1, NDC1,* and *Nup43*) might be related to potential hybridization potential long suspected for the genus. Interestingly, nucleoporin genes are consistently lost in other freshwater lineages (see Supplementary Note 7.4).

In summary, sponges in general do not show high levels of gene loss, and *E. muelleri* shows only a slightly higher level of loss than most Porifera. Ctenophores exhibit the highest rates of gene loss in the species sampled when compared to outgroup taxa, regardless of placement sister to Metazoa, or with sponges sister to Metazoa, in our analysis.

**7.2 Gene overlap and novelty in freshwater sponges**

From the 14 species included in this analysis, a total of 30,387 orthogroups were identified. Of these orthogroups 2,794 contained sequences from all 14 species and could be used for selection tests, as described in Supplementary Note 7.3 below. From our Orthofinder2 analysis, we were also able to investigate patterns of gene gain and loss across Spongillidae (freshwater sponges, 6 used here) and all outgroup non-spongillid taxa (8 here, from a number of demosponge species, including 3 genomes sequenced to a variety of depths.

Supplementary Figure 21 shows these groups together with a phylogeny showing the relationships of the species used in this analysis (Supplementary Figure 21A). Mapped onto this tree are the duplication and gain events inferred at nodes in the tree. It is notable that at the base of the Spongillida, and within this freshwater sponge clade, duplication is more prevalent than in other clades in the Heteroscleromorpha. Duplication is detected at relatively high rates within the Keratosa, but there are fewer sampled taxa in that group, and those duplications are expected to be ancestral, occuring at some point in the stem lineage of that relatively undersampled clade, rather than unique to those species sampled.

We used Orthofinder2 results to gain an understanding of the degree to which gene complements were shared between *E. muelleri* and other species of sponge. Supplementary Figure 21B shows these results, with the species with the highest overlap in orthogroup (and thus gene) complements shown in green, and those with the lowest shown in red. Of the species examined here, surprisingly *Dendrilla antarctica* and *Spongilla lacustris* have the greatest overall similarity, despite belonging to separate families within the Keratosa. However, in general all the members of the Spongillida are very similar in orthogroup content. Lake Baikal sponges are the most similar (which is not surprising, given their relatively recent diversification). *E. muelleri* itself is more similar to the Lake Baikal clade than it is to *S. lacustris* and *E. fragilis*, but the differences in shared content are slight.

Interestingly, other sequenced sponge genomes (*T. wilhelma* and *A. queenslandica*; note *X. testudinaria* is only sequenced to a low level of contiguity) match more sequences in non-spongillid species than *E. muelleri*. In all comparisons between these three genomic resources and non-Spongillid complements, *E. muelleri* has fewer matches than *T. wilhema* and *A. queenslandica* to orthogroups outside the Spongillida. This suggests a degree of loss within the freshwater sponge lineage, as noted in

Supplementary Note 7.1 above. However, these sequences could also be shared microbiome sequences found in marine species, but not in freshwater sponges.

Supplementary Figure 21C shows the percentage of genes from each species assigned to orthogroups. Those not placed into orthogroups could be novel genes or artifacts in gene prediction. *E. muelleri* has very similar numbers of genes placed in orthogroups, and of species-unique orthogroups, as other freshwater lineages, and those of the previously sequenced sponge genomes (*A. queenslandica* and *T. wilhelma).* Supplementary Figure 21C also shows the percentage of species unique orthogroups – those genes for which several copies are present , but only seen in individual species. Freshwater sponges are sampled in more depth than other clades included in this analysis, so the smaller numbers of unique orthogroups are not a surprise, as they are both representatives of a recent radiation, and have closely related species to compare complements with. However, due to the deeper sampling in *E. muelleri*, unique orthogroups can be reliably assumed to be specific to the lineage.

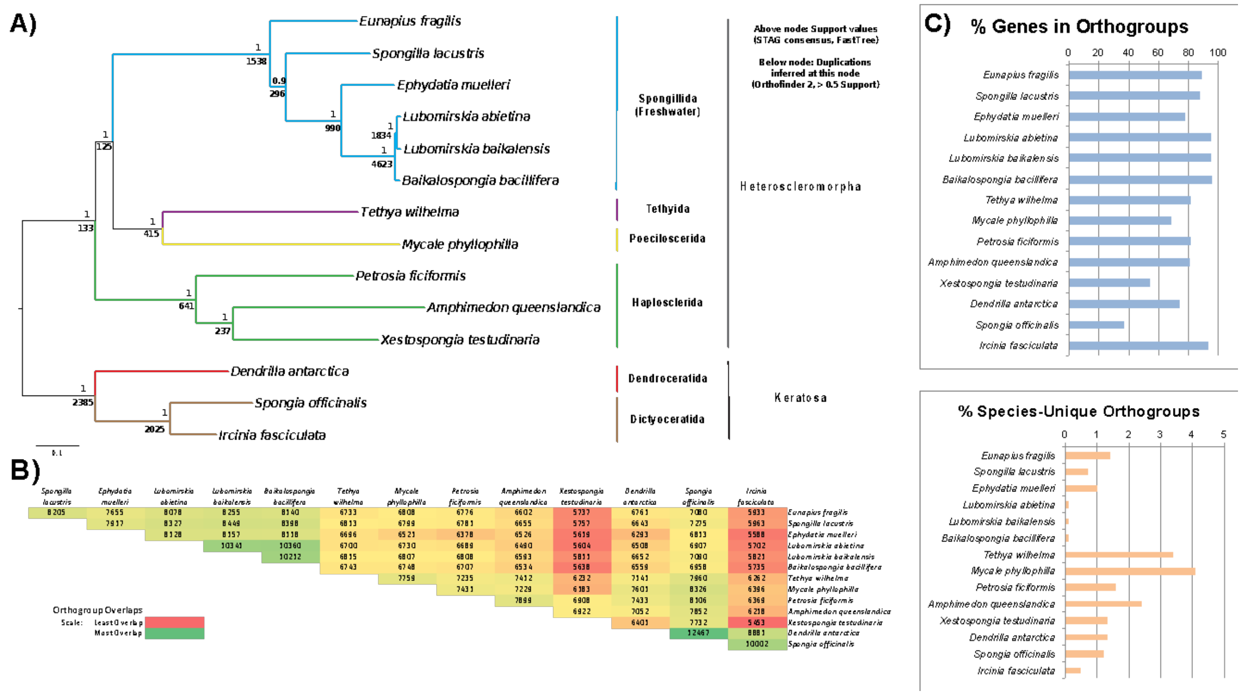

**Supplementary Figure 21:** A) The phylogenetic tree used for testing of selection, constructed within the Orthofinder2 framework, using the STAR method. Numbers at bases of nodes represent (above) support values inferred by STAR for each node, and (below) the number of gene duplications that were inferred to map to this nodal position on the tree (note - numbers for individual species not shown, but represented in C). B) Matrix showing numbers of orthogroups shared between species used in the analysis. Shading indicates the number of shared orthogroups for each species pair, red representing the least shared orthogroups and green the most. C) Upper: Percent of genes mapped to orthogroups for each species used in this analysis. Lower: Percent of orthogroups that were found to be unique to each species.

In short, at the base of the Spongillida, and within this freshwater sponge clade, duplication is more prevalent than in other clades in the Heteroscleromorpha. Duplication therefore plays a large role in the generation of novelty in freshwater sponges in general, and in *E. muelleri* in particular. Coupled with well-tested examples of species-unique orthogroups, which comparisons with closely related freshwater sponges show to be unique to *E. muelleri* itself, many novel sequences have been incorporated into the *E. muelleri* gene set, and it is these genes, along with those novelties shared with other Spongillida, that have likely allowed their success in freshwater environments worldwide.

**7.3 Positive selection**

Tests for positive selection were carried out twice, once with *E. muelleri* alone specified as the foreground taxon for selection tests, and once with all members of the Spongillida specified as foreground taxa. Selection tests were performed according to a scheme put forward by Santagata [68] and used in Kenny et al [23,]. In short, CODEML [69, 70] was used to test gene-level selection null vs alternative hypotheses, and differences in LnL used for $\chi^2$ tests of significance. Multiple comparison FDR was corrected [71]. Bayes Empirical Bayes (BEB) values used to identify specific sites under selection [72]. BUSTED, aBSREL and MEME tests of branch-level and site-level selection were run in HyPHy [73,74, 75, 76].

Of the 2,794 orthogroups represented in all 14 species, 1,751 were retained after curation with Phylotreepruner (removing sequences with out-paralogs, invariant sequences and orthogroups where alignments did not overlap in testable ways). These were subjected to tests for positive selection using both CODEML and the HyPhy suite. These were tested in two ways – once with *E. muelleri* alone noted as the foreground (test) taxon, and once with all freshwater sponges present in the test set noted in this way. A curated summary of these results, per gene, under each test, can be found in Supplementary Data 6. The full results of this analysis, along with alignments used and the json outputs of these tests, can also be found in this Supplementary Data 6.

From total results, we used the consilience of multiple tests to gain firm evidence for genes under positive selection. From our CodeML, Busted and aBSREL results (using a branch-site model, alignment-wide episodic diversifying selection, and adaptive branch-site random effects likelihood, respectively), we retained only genes with statistically significant evidence of selection under all three tests. For *E. muelleri* alone, this resulted in 117 orthogroups under positive selection with consilient evidence and full statistical support. These are shown in Supplementary Table 11.

When all freshwater species were tested as foreground taxa, only 33 orthogroups were found to be under positive selection. This lower number may be explained by the fact that extra lineages involved contribute noise in the form of extra data, as well as possibly contrasting selection pressures (Supplementary Table 12, with those genes overlapping with our '*E. muelleri* alone' test indicated at right). There is good, but not complete, overlap between these two data sets (Supplementary Figure 22A). Of the 117 genes with consilient evidence for positive selection when *E. muelleri* is tested alone, 23 overlap with the 'Spongillida as a whole' test. The larger number of genes found to be under positive selection in *E. muelleri* will be due to the increased clarity of the test without 'noise' in the form of varying data from other taxa, indicating the specific selection pressures that have influenced *E. muelleri* since its split from its last common ancestor with these species to be noted. From both of these tests, we took forward the 117 orthogroups noted as under positive selection in *E. muelleri* for further analysis. The 10 remaining genes significant in freshwater sponges as a whole cannot be said to have changed significantly in *E.muelleri,* and are not used in our further analysis.

*Genes under selection*

A diverse range of annotated orthogroups are noted in our tests for positive selection (Supplementary Tables 11 and 12). Annotation was made simple by the inclusion of the *A. queenslandica* genome in our dataset, with its history of use in a range of studies. Our orthogroups matched genes responsible for both housekeeping and more specialised roles within the cell, and agreed with previously published research into freshwater specialisation in sponges and beyond.

There are a number of examples of housekeeping genes annotated in our orthogroups. Some of these housekeeping genes are clearly structural, such as *actin-3-like,* several *lamin* genes, *kinesin-like protein KIF23, centromere/microtubule-binding protein CBF5-like isoform X3* and *septin-7-like isoform X1.* Others perform roles in maintaining normal cellular function, including genes such as *cytochrome P450 4F1-like* and numerous mitochondrially located genes. Of interest are *hematopoietic prostaglandin D synthase* and *prostaglandin reductase 1-like* which possibly perform inter-related roles. While many genes involved in housekeeping functions have been seen in previous tests of freshwater adaptation, the positive selection of prostaglandin-related genes may be specific to *E. muelleri* and possibly related to predator deterrence [77].

It is known that the move to freshwater conditions must be accompanied by a diverse range of changes to membrane functionality, particularly in intramembrane proteins and ion channels. Several genes known to perform roles in homeostasis and membrane function were noted in our dataset. Examples of this include *V-type proton ATPase subunit B*, three kinds of *sorting nexin*, *vacuolar-sorting protein SNF8* and *Multidrug and toxin extrusion protein 1*. It is easy to understand how these proteins may be under special pressure to evolve to better suit freshwater environments.

More specialised genes which may play a broader role in regulating homeostasis and cellular responses are also observed. Genes such as *bone morphogenetic protein receptor type-1B-like*, *ras-related protein O-Krev* and *dual specificity mitogen-activated protein kinase kinase 2-like (MAP2K2)* belong to

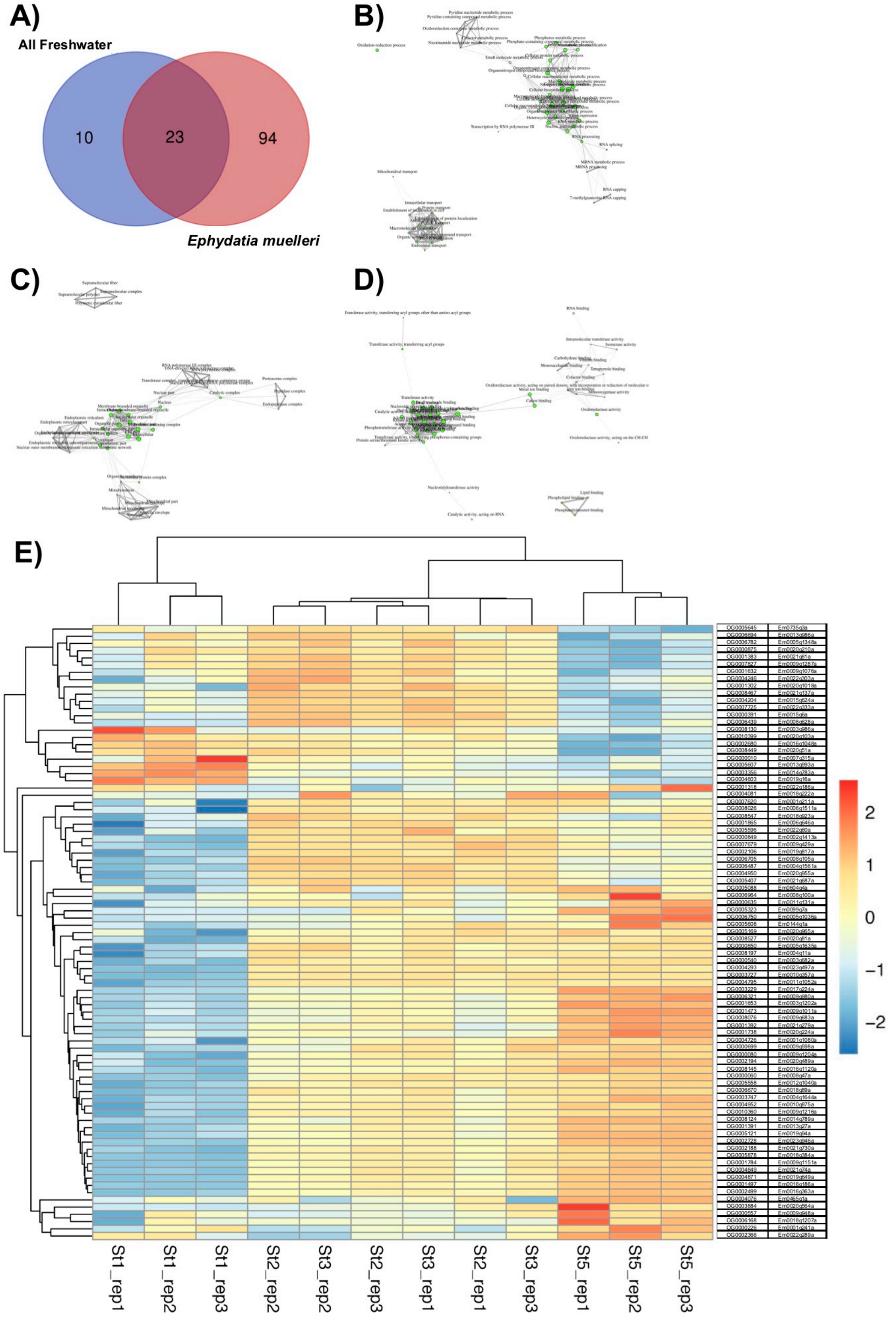

**Supplementary Figure 22:** Summary of results of selection tests. A) distribution of orthogroups with consilient support in tests of positive selection in the freshwater clade as a whole (left, blue) and *E. muelleri* alone (right, red). B-D) distribution of over-represented GO terms in genes identified as under positive selection in *E. muelleri.* These are separated by sub-ontology as follows: B) Biological Process, C) Cellular Component, D) Molecular Function. Note clustering of GO terms into groups of related identity. E) Expression profile of genes under selection. Stages of development are along the bottom, Stages 1, 2, 3 and 5 as St1, St2, St3 and St5 (for further information on stages, see Supplementary Note 11). Of the genes shown in Supplementary Table 11, 85 were differentially expressed in the course of development. Full numerical expression levels, annotations and further details provided in Supplementary Data 6.

signalling cascades which can control a variety of key processes in sponge growth and development. The presence of signatures of positive selection in these genes suggests that these pathways may play specialised roles in *E. muelleri* in particular.

When compared to previous work in freshwater sponges, there is a considerable overlap in genes found to be under positive selective pressure. The only previous investigation of this type in sponges [23] also found genes such as *actin-3-like, probable ATP-dependent RNA helicase DDX49,* and *T-complex protein 1 subunit* to be under positive selective pressure. More broadly, that resource and this found several representatives of apoptosis inducing factors, rho GTPases, serine/threonine protein kinases, vacuolar sorting proteins, dnaJ homologues and other protein families to be represented. It is useful but not surprising that the same genes were found in this analysis, as the same technique was used to identify genes in both.

*Differential expression of genes under selection*

Interestingly, almost all genes with signatures of positive selection were differentially expressed across the process of development (Supplementary Figure 22E, with full details listed in Supplementary Data 6). Of the 117 *E. muelleri* genes within the orthogroups under positive selection, 85 are differentially expressed at these time points (Time points and RNAseq are described in Supplementary Note 8.1). This includes orthogroups such as OG0004726 (Em0001g1080a), which encodes the *RAP1* gene, which is vital for cell adhesion and junction formation [78]. The protein encoded by this gene would experience markedly different conditions in freshwater when compared to marine environments. Other genes included in this differentially expressed complement include nucleoporins (OG0004246/Em0022g303a), vacuolar sorting proteins (OG0008197/Em0004g11a) and the solute carrier called the 'multidrug and toxin extrusion protein' (OG0000060/Em0008g47a).

The process of development is a clearly crucial time in any organism's life. Genes expressed at this time point will be under constant pressure from natural selection, and the process of adapting to a new environment (such as freshwater, in the case of *E. muelleri)* will require a number of changes in their sequence. It is therefore no surprise that we note numerous genes with signals of positive selection in the gene set to be differentially expressed at this time. The presence of numerous housekeeping genes, and particularly those known to perform roles in homeostasis and membrane function, as noted in Supplementary Note 7.2, is entirely congruent with this finding.

These genes, and the specific changes they have accumulated in the process of freshwater adaptation, are therefore ideal candidates for future investigation in *E. muelleri* and in other species that have convergently adapted from a marine habitat. They are of clear functional utility (as evidenced by their differential expression) and under strong selective pressure. If these genes were also observed to change convergently in other freshwater-adapted lineages, this would suggest that these conditions impose common constraints, which natural selection solves in similar ways.

*GO enrichment analysis*

To gain a holistic view of the links between these apparently unconnected genes, we used the ShinyGO v0.61 data visualisation tool [79]. This calculates over-represented GO terms within a gene set, and displays the GO terms of these, and their links to other GO terms in the same dataset. Accession numbers of *A. queenslandica* orthologues within the orthogroups were input with 0.05 P-value cutoff for FDR. ShinyGO automatically retrieved GO terms, determined enriched GO terms within this set, corrected for FDR, and displayed links between GO terms.

These results are displayed (separated by sub-ontology) in Supplementary Figure 22B-D. Note that while the gene names shown in Supplementary Tables 11 and 12 are disparate at first glance, under all three sub-ontologies GO terms are generally clustered, rather than evenly spread. This indicates that the genes mapped to these GO terms perform inter-linked roles within the cell.

To note easily which kind of genes are over-represented in our dataset, we have shown the most common GO terms (and the degree to which they are over-represented) in Supplementary Table 13. The most enriched terms in our dataset are cellular components (coded under a number of GO terms) and metabolic pathways. Notably among metabolic pathways, Nitrogen Compound Metabolic Processes, along with those related to cyclic/aromatic compound processing, are highly significant.

Less obviously significant, but with 2/5 components present in our list of genes under positive selection, the RNA polymerase III complex is noted as over-represented. This polymerase is responsible for the transcription of ribosomal 5S, tRNA and other small RNA genes [80]. As such its operation is intimately related to cell growth and the cell cycle. It also is responsible for the transcription of some regulatory RNAs in other species, including miRNA [81]. Changes to the sequence and operation of the RNA polIII complex could therefore be having a range of downstream effects on the biology of *E. muelleri.*

We should note that 1,751 orthogroups likely represent 10% or less of the gene complements of the sponges tested. There are many other genes that will be important for adaptation to freshwater conditions that were not tested by our analysis, due to absence in one or more of the resources used. As more contiguous genome assemblies become available from the Porifera, this could be re-visited to ensure that the entire set of genes used in the process of adaptation can be catalogued and understood.

Not discussed here, but presented in detail in Supplementary Data 6, are the specific sites under selection in each of these orthogroups, as assessed using both MEME and BEB methods. For all orthogroups studied here, the specific amino acid residues under positive selection have been noted, and this data will be useful for researchers interested in specific genes. Previous studies (as discussed in [23]) have found that these sites tend to be found in intramembrane domains, likely as an adaptation to freshwater conditions, and the markedly different osmotic effects of such an environment.

**Supplementary Table 11:** Genes identified as being under positive selection in *E. muelleri.* Accession numbers and annotations derived from best hit in the *nr* database (almost universally to *Amphimedon queenslandica).* Further details on individual sites under selection, changes compared to ancestral pattern, and P values associated are all available for download as Supplementary Data 6. Expression of genes across development, Supp Fig 22.

| Orthogroup | Accession number | Best hit gene | Orthogroup | Accession number | Best hit gene | Orthogroup | Accession number | Best hit gene |
|---|---|---|---|---|---|---|---|---|
| OG0000002 | XP_019856229.1 | uncharacterized protein LOC109584796 isoform X5 | OG0002499 | XP_003385030.1 | nicotinamide phosphoribosyltransferase-like | OG0005903 | XP_011409647.1 | DNA-directed RNA polymerase III subunit RPC6-like |
| OG0000010 | XP_011407636.1 | piggyBac transposable element-derived protein 4-like | OG0002502 | XP_003384737.3 | protein SON-like isoform X2 | OG0005910 | XP_003383283.1 | V-type proton ATPase subunit B |
| OG0000021 | XP_019861849.1 | uncharacterized protein LOC109590368 | OG0002629 | XP_019849717.1 | alpha-aminoadipic semialdehyde synthase, mitochondrial-like | OG0006167 | XP_011410392.2 | interleukin enhancer-binding factor 2 homolog |
| OG0000060 | XP_019856923.1 | multidrug and toxin extrusion protein 1-like isoform X1 | OG0002680 | XP_003391996.1 | methionine aminopeptidase 1-like, partial | OG0006168 | XP_003384418.1 | mitochondrial import inner membrane translocase subunit Tim21-like |
| OG0000080 | XP_003384376.1 | actin-3-like | OG0002728 | XP_019853346.1 | SH3 domain-containing kinase-binding protein 1-like isoform X2 | OG0006321 | XP_003386746.1 | sorting nexin-2-like isoform X1 |
| OG0000193 | XP_011410522.1 | tetratricopeptide repeat protein 38-like | OG0002786 | XP_019848782.1 | LIM domain-binding protein 3-like | OG0006357 | XP_019856000.1 | midnolin-like |
| OG0000226 | XP_019858579.1 | arrestin domain-containing protein 3-like | OG0002879 | XP_003383503.1 | methylmalonyl-CoA mutase, mitochondrial-like | OG0006439 | XP_011402894.1 | splicing factor 3B subunit 2-like |
| OG0000335 | XP_003386175.1 | probable 4-coumarate--CoA ligase 1 | OG0003229 | XP_019859992.1 | lethal(3)malignant brain tumor-like protein 1 | OG0006449 | XP_003385038.1 | grpE protein homolog 1, mitochondrial-like |
| OG0000391 | XP_003385851.1 | serine/threonine-protein phosphatase alpha-2 isoform | OG0003356 | XP_003390645.1 | dnaJ homolog subfamily B member 13-like | OG0006480 | XP_003383939.1 | protein farnesyltransferase/geranylgeranyltransferase type-1 subunit alpha-like |
| OG0000540 | XP_019860622.1 | bone morphogenetic protein receptor type-1B-like | OG0003727 | XP_003384468.1 | kinesin-like protein KIF23 | OG0006487 | XP_003383674.1 | glycylpeptide N-tetradecanoyltransferase 2-like |
| OG0000557 | XP_003391770.2 | prostaglandin reductase 1-like, partial | OG0003747 | XP_011403428.2 | serine/threonine-protein phosphatase 6 regulatory ankyrin repeat subunit A-like | OG0006500 | XP_003382982.1 | 26S proteasome non-ATPase regulatory subunit 6-like |
| OG0000635 | XP_019849740.1 | cytochrome P450 4F1-like | OG0003878 | XP_003385504.1 | arf-GAP with dual PH domain-containing protein 1-like | OG0006670 | XP_011406756.1 | WW domain-containing oxidoreductase-like |
| OG0000699 | XP_003386741.1 | hematopoietic prostaglandin D synthase-like | OG0003884 | XP_003385106.3 | protein ADP-ribosylarginine hydrolase-like | OG0006694 | XP_019854962.1 | trimethylguanosine synthase-like |
| OG0000747 | XP_003382696.1 | uncharacterized protein LOC100635379 | OG0004076 | XP_003384813.1 | peflin-like | OG0006705 | XP_019854114.1 | procollagen-lysine,2-oxoglutarate 5-dioxygenase 3-like |
| OG0000849 | XP_011408847.2 | carboxypeptidase A5-like | OG0004081 | XP_019849345.1 | vesicle-associated membrane protein-associated protein B/C-like isoform X1 | OG0006750 | XP_019851060.1 | EF-hand domain-containing protein 1-like |
| OG0000850 | XP_019862581.1 | mRNA-capping enzyme-like, partial | OG0004204 | XP_003386963.1 | T-complex protein 1 subunit zeta-like | OG0006782 | XP_019849559.1 | uncharacterized protein LOC109580618 |
| OG0000875 | XP_003391408.2 | probable ATP-dependent RNA helicase DDX49, partial | OG0004246 | XP_019853816.1 | glycine-rich cell wall structural protein 1-like | OG0006964 | XP_003391973.2 | protein kinase C iota type-like, partial |
| OG0001172 | XP_019860835.1 | LOW QUALITY PROTEIN: iron-sulfur protein NUBPL-like | OG0004293 | XP_003384065.1 | dual oxidase maturation factor 1-like | OG0007091 | XP_003385302.1 | uncharacterized protein LOC100634227 isoform X1 |
| OG0001263 | XP_003386218.1 | trans-1,2-dihydrobenzene-1,2-diol dehydrogenase-like | OG0004603 | XP_019858094.1 | serine/threonine-protein kinase D1-like | OG0007131 | XP_003383886.2 | ribose-5-phosphate isomerase-like |
| OG0001302 | XP_003385512.1 | ATP-dependent RNA helicase dbp2-like | OG0004619 | XP_019855522.1 | endoplasmic reticulum resident protein 44-like isoform X2 | OG0007146 | XP_003383305.1 | clathrin interactor 1-like |
| OG0001318 | XP_019864138.1 | uncharacterized protein LOC109593544, partial | OG0004726 | XP_003383013.1 | ras-related protein O-Krev | OG0007620 | XP_019862998.1 | lamin-B receptor-like, partial |
| OG0001383 | XP_019850665.1 | centromere/microtubule-binding protein CBF5-like isoform X3 | OG0004795 | XP_019853969.1 | sorting nexin-7-like | OG0007650 | XP_003389688.1 | methyltransferase-like protein 14 homolog |
| OG0001391 | XP_019864389.1 | exocyst complex component 2-like | OG0004849 | XP_003389005.1 | EF-hand domain-containing family member C2-like | OG0007679 | XP_011405655.2 | lamin-L(I)-like isoform X1 |
| OG0001392 | XP_011409565.2 | tyrosine 3-monooxygenase-like | OG0004871 | XP_003387642.3 | pleckstrin homology domain-containing family D member 1-like isoform X1 | OG0007725 | XP_003386357.1 | ribonucleoside-diphosphate reductase large subunit-like |
| OG0001473 | XP_019863757.1 | myotubularin-related protein 10-B-like | OG0004925 | XP_003384129.1 | tRNA-splicing ligase RtcB homolog | OG0007827 | XP_011404360.1 | 28S ribosomal protein S30, mitochondrial-like |
| OG0001487 | XP_019858049.1 | serine/threonine-protein kinase 26-like | OG0004950 | XP_003382862.1 | mitochondrial dicarboxylate carrier-like | OG0008019 | XP_003388223.1 | COMM domain-containing protein 2-like |
| OG0001497 | XP_003387043.1 | band 7 protein AGAP004871-like isoform X1 | OG0004952 | XP_003382930.1 | COP9 signalosome complex subunit 2-like | OG0008026 | XP_003389696.1 | 3-ketoacyl-CoA thiolase, mitochondrial-like |
| OG0001632 | XP_019863974.1 | asparagine--tRNA ligase, cytoplasmic-like, partial | OG0005088 | XP_019851983.1 | carnitine O-palmitoyltransferase 2, mitochondrial-like isoform X1 | OG0008076 | XP_019853162.1 | uncharacterized protein LOC100639992 |
| OG0001653 | XP_003387510.1 | probable ethanolamine kinase | OG0005121 | XP_011402690.1 | WD repeat-containing protein 20-like | OG0008124 | XP_003385793.1 | EF-hand calcium-binding domain-containing protein 2-like |
| OG0001738 | XP_019849243.1 | sorting nexin-29-like | OG0005169 | XP_019852937.1 | protein HID1-like | OG0008130 | XP_003385641.1 | MOB kinase activator 2-like |
| OG0001784 | XP_011406309.2 | uncharacterized protein LOC100641267 | OG0005323 | XP_019851245.1 | radial spoke head protein 3 homolog B-like | OG0008145 | XP_003385121.1 | carboxyl-terminal PDZ ligand of neuronal nitric oxide synthase protein-like |
| OG0001805 | XP_011410501.2 | RNA-binding protein 25-like | OG0005407 | XP_019850467.1 | 26S proteasome non-ATPase regulatory subunit 1-like | OG0008197 | XP_003383211.1 | vacuolar-sorting protein SNF8-like |
| OG0001865 | XP_003386492.1 | omega-amidase NIT2-like isoform X2 | OG0005558 | XP_019854586.1 | MAGUK p55 subfamily member 6-like | OG0008213 | XP_003382661.1 | dolichyl-diphosphooligosaccharide--protein glycosyltransferase 48 kDa subunit-like |
| OG0002105 | XP_019864424.1 | WD repeat-containing protein 11-like isoform X1 | OG0005596 | XP_003385522.1 | apoptosis-inducing factor 1, mitochondrial-like | OG0008449 | XP_003387651.2 | 28 kDa heat- and acid-stable phosphoprotein-like |
| OG0002106 | XP_019864272.1 | L-2-hydroxyglutarate dehydrogenase, mitochondrial-like | OG0005607 | XP_019850041.1 | uncharacterized protein LOC100639981 | OG0008467 | XP_019852743.1 | spliceosome-associated protein CWC15 homolog |
| OG0002188 | XP_003383659.1 | desumoylating isopeptidase 1-like | OG0005608 | XP_003385059.1 | glutaminyl-peptide cyclotransferase-like | OG0008527 | XP_011410455.1 | dual specificity mitogen-activated protein kinase kinase 2-like |
| OG0002194 | XP_003382830.3 | ethanolamine-phosphate cytidylyltransferase-like isoform X2 | OG0005645 | XP_019859018.1 | T-complex protein 1 subunit eta-like | OG0008547 | XP_003383789.1 | proteasome subunit alpha type-7-like |
| OG0002227 | XP_019849657.1 | vacuolar protein sorting-associated protein 27-like | OG0005794 | XP_019856999.1 | DNA-directed RNA polymerase III subunit RPC9-like | OG0010360 | XP_003386748.1 | protein AF-9-like |
| OG0002366 | XP_003386945.1 | fatty acid hydroxylase domain-containing protein 2-like | OG0005878 | XP_003384646.1 | septin-7-like isoform X1 | OG0010399 | XP_011410454.1 | galactokinase-like |

**Supplementary Table 12:** Genes identified as being under positive selection when the Spongillida as a whole is specified as the clade under selection. Further details on individual sites under selection, changes compared to ancestral pattern, and P values associated are all available for download as Supplementary Data 6.

| Orthogroup | Accession number | Best hit gene | Also under positive selection in *E. muelleri* alone? |
|---|---|---|---|
| OG0000010 | XP_011407636.1 | piggyBac transposable element-derived protein 4-like | Yes |
| OG0000021 | XP_019861849.1 | uncharacterized protein LOC109590368 | Yes |
| OG0000056 | XP_003391783.2 | receptor-type tyrosine-protein phosphatase F-like, partial | No |
| OG0000061 | XP_019852875.1 | TNF receptor-associated factor 2-like | No |
| OG0000193 | XP_011410522.1 | tetratricopeptide repeat protein 38-like | Yes |
| OG0000195 | XP_019854217.1 | protein F37C4.5-like | No |
| OG0000270 | XP_011407264.2 | autophagy-related protein 16-1-like | No |
| OG0000540 | XP_019860622.1 | bone morphogenetic protein receptor type-1B-like | Yes |
| OG0000572 | XP_003384736.1 | VPS10 domain-containing receptor SorCS1-like | No |
| OG0000730 | XP_019855106.1 | fatty aldehyde dehydrogenase-like | No |
| OG0000992 | XP_003391448.2 | ras-related protein Rab-22A-like | No |
| OG0001047 | XP_003385695.1 | vacuolar protein sorting-associated protein 4B-like | No |
| OG0005088 | XP_019851983.1 | carnitine O-palmitoyltransferase 2, mitochondrial-like isoform X1 | Yes |
| OG0005323 | XP_019851245.1 | radial spoke head protein 3 homolog B-like | Yes |
| OG0005407 | XP_019850467.1 | 26S proteasome non-ATPase regulatory subunit 1-like | Yes |
| OG0006100 | XP_019854052.1 | pleckstrin-2-like | No |
| OG0006321 | XP_003386746.1 | sorting nexin-2-like isoform X1 | Yes |
| OG0006670 | XP_011406756.1 | WW domain-containing oxidoreductase-like | Yes |
| OG0006694 | XP_019854962.1 | trimethylguanosine synthase-like | Yes |
| OG0006750 | XP_019851060.1 | EF-hand domain-containing protein 1-like | Yes |
| OG0006782 | XP_019849559.1 | uncharacterized protein LOC109580618 | Yes |
| OG0007131 | XP_003383886.2 | ribose-5-phosphate isomerase-like | Yes |
| OG0007620 | XP_019862998.1 | lamin-B receptor-like, partial | Yes |
| OG0007679 | XP_011405655.2 | lamin-L(I)-like isoform X1 | Yes |
| OG0007725 | XP_003386357.1 | ribonucleoside-diphosphate reductase large subunit-like | Yes |
| OG0008019 | XP_003388223.1 | COMM domain-containing protein 2-like | Yes |
| OG0008124 | XP_003385793.1 | EF-hand calcium-binding domain-containing protein 2-like | Yes |
| OG0008145 | XP_003385121.1 | carboxyl-terminal PDZ ligand of neuronal nitric oxide synthase protein-like | Yes |
| OG0008213 | XP_003382661.1 | dolichyl-diphosphooligosaccharide--protein glycosyltransferase 48 kDa subunit-like | Yes |
| OG0008449 | XP_003387651.2 | 28 kDa heat- and acid-stable phosphoprotein-like | Yes |
| OG0008467 | XP_019852743.1 | spliceosome-associated protein CWC15 homolog | Yes |
| OG0008547 | XP_003383789.1 | proteasome subunit alpha type-7-like | Yes |
| OG0008839 | XP_003387700.1 | aristaless-related homeobox protein-like | No |

**Supplementary Table 13:** Results of enrichment analysis of genes under positive selection in *E. muelleri.* Numbers of genes with given GO annotation are given both for the gene list determined for *E. muelleri,* and the figure in *A. queenslandica*'s complete gene complement.

| Enrichment FDR | Genes in list (of 117) | Total genes (*A. queenslandica*) | Functional Category |
|---|---|---|---|
| 9.65E-08 | 26 | 2596 | Intracellular part |
| 9.65E-08 | 15 | 757 | Metabolic pathways |
| 8.08E-07 | 28 | 3375 | Cell |
| 8.25E-07 | 27 | 3197 | Intracellular |
| 1.15E-06 | 33 | 4733 | Cellular metabolic process |
| 1.17E-06 | 27 | 3315 | Cell part |
| 2.12E-06 | 33 | 4922 | Nitrogen compound metabolic process |
| 3.53E-06 | 32 | 4794 | Ion binding |
| 1.26E-05 | 15 | 1236 | Cytoplasm |
| 1.94E-05 | 22 | 2714 | Anion binding |
| 2.86E-05 | 14 | 1176 | Protein-containing complex |
| 2.86E-05 | 17 | 1740 | Membrane-bounded organelle |
| 2.86E-05 | 19 | 2156 | Intracellular organelle |
| 3.44E-05 | 19 | 2194 | Organelle |
| 5.51E-05 | 19 | 2277 | Cellular aromatic compound metabolic process |
| 6.03E-05 | 19 | 2311 | Organic cyclic compound metabolic process |
| 6.03E-05 | 4 | 42 | Phosphatidylinositol binding |
| 6.35E-05 | 28 | 4554 | Macromolecule metabolic process |
| 6.93E-05 | 16 | 1712 | Intracellular membrane-bounded organelle |
| 6.93E-05 | 10 | 648 | Oxidoreductase activity |
| 0.00011769 | 10 | 696 | Oxidation-reduction process |
| 0.00011769 | 17 | 2018 | Transferase activity |
| 0.000122835 | 7 | 300 | RNA processing |
| 0.000131171 | 20 | 2736 | Small molecule binding |
| 0.000132098 | 19 | 2530 | Cellular nitrogen compound metabolic process |
| 0.000132098 | 18 | 2283 | Heterocycle metabolic process |
| 0.000132098 | 4 | 58 | Phospholipid binding |
| 0.000132098 | 13 | 1254 | Gene expression |
| 0.000132098 | 12 | 1061 | RNA metabolic process |
| 0.000148601 | 11 | 908 | Cytoplasmic part |
| 0.000167504 | 21 | 3078 | Organonitrogen compound metabolic process |
| 0.000234433 | 16 | 1968 | Drug binding |
| 0.000250455 | 12 | 1156 | Intracellular organelle part |
| 0.000261714 | 17 | 2223 | Nucleobase-containing compound metabolic process |
| 0.000281027 | 12 | 1177 | Organelle part |
| 0.000453414 | 2 | 5 | RNA polymerase III complex |
| 0.00050855 | 15 | 1898 | ATP binding |
| 0.000517631 | 15 | 1910 | Adenyl nucleotide binding |
| 0.000517631 | 15 | 1909 | Adenyl ribonucleotide binding |
| 0.000549412 | 18 | 2640 | Nucleotide binding |
| 0.000549412 | 18 | 2640 | Nucleoside phosphate binding |
| 0.000586385 | 3 | 35 | Proteasome |
| 0.000846852 | 6 | 313 | Endomembrane system |
| 0.001103084 | 4 | 112 | Lipid binding |
| 0.001325278 | 21 | 3651 | Cellular macromolecule metabolic process |
| 0.001550221 | 2 | 10 | RNA capping |
| 0.001550221 | 2 | 10 | 7-methylguanosine RNA capping |
| 0.001819522 | 5 | 244 | Amide transport |
| 0.001819522 | 5 | 243 | Establishment of protein localization |
| 0.001819522 | 14 | 1968 | Nucleic acid metabolic process |

### 7.4 Comparison with other freshwater lineages

To gain an insight into the genomic changes needed when occupying a freshwater niche, we compared the gene complement of *Ephydatia muelleri* to a range of other metazoan datasets, including species that have independently undergone a marine to freshwater transition. The taxa compared can be seen in Supplementary Figure 23 below, and include four phyla with one internal clade that independently moved into freshwater.

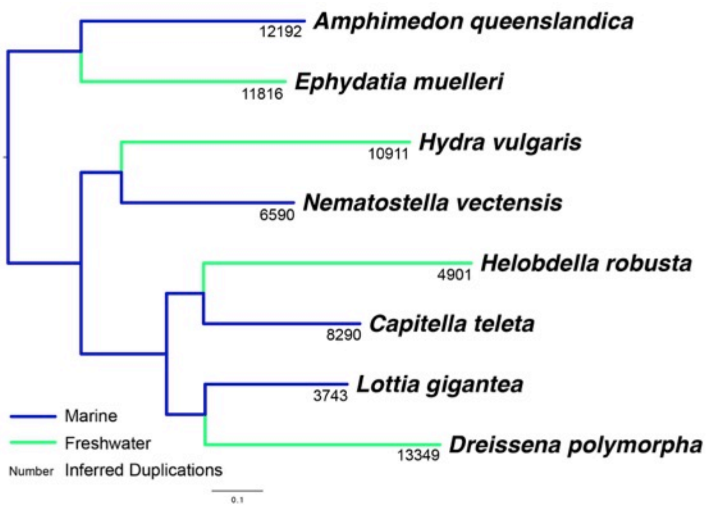

**Supplementary Figure 23:** Phylogeny showing inter-relationships between marine and freshwater species tested. Phylogeny inferred in Orthofinder2 using the STAR method. Numbers at tips of nodes are inferred duplication events in a lineage given the phylogeny shown.

Gene duplication in particular does not seem to be a universally contributing factor towards adaptation to freshwater environments. If this was universally useful for adaptation to freshwater, we would expect to see higher numbers of duplicates in all freshwater lineages. Deeper sampling-based analysis of freshwater sponge lineages (see Supplementary Note 7.2) shows a burst of retained duplication events at the base of the freshwater sponge radiation. However, in four pairwise comparisons, only two of four showed more duplications in the freshwater lineage. In the remaining cases, there are more duplications in marine lineages.

We used these results to see if there were any genes that were lost in every freshwater lineage. While some shared losses can be mathematically expected by chance, shared loss of certain categories of gene could be informative for understanding what is needed to evolve to freshwater conditions. There are 29 orthogroups that are lost in all freshwater lineages but present in all marine lineages. To estimate the role of chance in this result, we calculated the genes that are lost in all marine lineages but present in all freshwater lineages, and found only 4. The average loss rate for any 4 taxa was 16.9 genes.

Similarly, a large number of shared losses are seen spanning three of the four freshwater lineages. The four marine taxa are all present in orthogroups with only one of the freshwater species (only a single representative from *E. muelleri, D. polymorpha, H. vulgaris* or *H. robusta*) respectively 73, 22, 52 and 37 times (indicating near complete loss of these orthogroups in freshwater, but complete retention in marine species). By way of contrast, four freshwater taxa are all present in orthogroups with only one marine species 1, 2, 8 and 0 times (which would indicate near complete loss in marine lineages for these orthogroups). This difference indicates significantly higher ($p$=0.013, $t$ statistic) loss in freshwater lineages. Shared loss in freshwater lineages therefore seems to be a reasonably common phenomenon, whereas it happens very seldom across marine taxa.

**Supplementary Table 14**: The 29 orthogroups absent from all four freshwater lineages examined domain-containing proteins are also present (kishB, transmembrane protein 242, amino acid transporter AVT3B) alongside genes of unknown function.

| OG name | Best Blast Hit ID | PFAM domains |
|---|---|---|
| OG0010707 | WD repeat containing protein 18 | WD40,Nup160,Nucleoporin_N,DUF4331 |
| OG0009598 | TEPP (testis, prostate and placenta-expressed protein | RNA_pol_3_Rpc31 |
| OG0010693 | amino acid transporter AVT3B isoform X2 | Aa_trans,MotA_ExbB,DUF4079 |
| OG0010670 | dynein beta chain, flagellar outer arm-like isoform X14 | DHC_N1,DUF2408,DHC_N2,MT,AAA_8,AAA_7,AAA_6,Myosin_tail_1,TBLOP,DALR_1,DivIC,DUF4441 |
| OG0009648 | protein kish-B | DUF1242,Jag_N,GP46 |
| OG0010663 | glyoxalase domain-containing protein 5-like isoform X2 | Glyoxalase_2,Glyoxalase,Glyoxalase_4,tRNA_Me_trans,Glyoxalase_3,Ub-Mut7C,MR_MLE_C,MR_MLE_N,MR_MLE,Sigma70_r1_1 |
| OG0010682 | DNA polymerase delta subunit 4-like | DNA_pol_delta_4,Paramyx_P_V_C |
| OG0010717 | uncharacterized protein | |
| OG0003180 | metabotropic glutamate receptor 3-like | ANF_receptor,OrfB_Zn_ribbon,Peripla_BP_6,HSF_DNA-bind,HSP70,FtsA,MreB_Mbl,StbA,7tm_3,SUR7 |
| OG0007786 | viral helicase | SH3_9,SH3_2,SH3_1,SH3_3,YfkD,Peptidase_S24,NACHT,AAA_17,AAA_18,AAA_22,AAA,NB-ARC,IstB_IS21,NTPase_1,AAA_19,AAA_16,AAA_14,AAA_24,Viral_helicase1,MobB,RNA_helicase,AAA_28,hSH3,AAA_11,ABC_tran,AAA_33,RuvB_N,T2SE,DUF258,AAA_5,cobW,Mg_chelatase,Arch_ATPase,AAA_10,SRP54,AAA_30,AAA_25,ATP_bind_1,DSX_dimer,LptC,SH3_4 |
| OG0010714 | transmembrane protein 242-like isoform X2 | DUF1358,DUF3040 |
| OG0009642 | WD repeat-containing protein 88 | WD40,Zn-ribbon_8,Nup160,PQQ_2,Cytochrom_D1,DUF3312,SEFIR,DUF2084,BBS2_Mid,Nucleoporin_N |
| OG0008744 | WD repeat-containing protein 31-like isoform X2 | WD40,Nup160,Ricin_B_lectin,AA_permease_2,AA_permease,eIF2A,Nucleoporin_N,DUF2031,DUF3040,DUF4337,Coatomer_WDAD,DUF3312,PQQ_2 |
| OG0009640 | protein Churchill-like | Churchill,Ribosomal_L32p,PolC_DP2,Prok-RING_4,Kazal_1,TF_Zn_Ribbon,Lar_restr_allev,Terminase_GpA,Viral_NABP,zf-ribbon_3,Zn-ribbon_8,Elf1,Ribosomal_L37e,Baculo_LEF5_C,TFIIS_C,DUF2072,LIM,Cytochrome_C7,DUF3795,zf-C4pol,DZR,Cytochrom_c3_2,zinc_ribbon_2,Cytochrome_CBB3 |
| OG0009630 | uncharacterized LOC protein | Rhodanese,N1221,PTS_IIB |
| OG0010731 | peroxisomal 2,4-dienoyl-CoA reductase-like | adh_short_C2,adh_short,KR,Epimerase,F420_oxidored,Polysacc_synt_2,Shikimate_DH,3Beta_HSD,Saccharop_dh,3HCDH_N,NAD_binding_10,His_biosynth |
| OG0010653 | uncharacterized LOC protein | PDZ,PDZ_2 |
| OG0010652 | elongation of very long chain fatty acids protein 4-like | ELO,COX14,DUF2484,DUF3270 |
| OG0010744 | uncharacterized LOC protein | Nop14 |
| OG0008733 | basal body-orientation factor 1-like | DUF4515,DUF972,BLOC1_2,PilO,NPV_P10,FlxA,DUF4407,V_ATPase_I,IncA,DUF948,DUF3573,Atg14,GAS,EssA,DUF4239,DUF4618,PPO1_KFDV,RuvC,PCP,PNGaseA,TroA,Barstar |
| OG0010699 | uncharacterized LOC protein | EF-hand_1,EF-hand_8,EF-hand_7,EF-hand_6,EF-hand_5,EF-hand_4,Telomere_Sde2 |
| OG0009634 | N-sulphoglucosamine sulphohydrolase-like | Sulfatase,Phosphodiest,DUF229,DUF1501,Sulfatase_C |
| OG0009644 | mitotic-spindle organizing protein 2-like | MOZART2,MOZART1,DUF4159 |
| OG0008757 | uncharacterized LOC protein | LRR_4,LRR_8,LRR_9,LRR_1,LRR_7 |
| OG0010750 | ornithine carbamoyltransferase, mitochondrial-like isoform X2 | OTCace,OTCace_N,DUF2540,OCD_Mu_crystall,DUF3783 |
| OG0008756 | mitochondrial enolase superfamily member 1-like isoform X1 | MR_MLE_C,MR_MLE,MR_MLE_N,MAAL_C,PepSY,HTH_17 |
| OG0010746 | nicolin-1-like isoform X5 | DUF3770,CBM_11 |
| OG0009647 | ubiquitin carboxyl-terminal hydrolase 40-like | UCH,UCH_1,DNA_RNApol_7kD,Mu-like_Com,zf-C2H2_6,zf-BED,zf-C2H2,ubiquitin,DUF2369,Ubiquitin_2,YukD |
| OG0008726 | ubiquitin-conjugating enzyme E2 T | UQ_con,Prok-E2_B,RWD,UEV |

We examined the identity of the 29 genes that were lost in all four freshwater lineages, and these are given in Supplementary Table 14 above. They belong to a wide variety of protein families, and several of the genes represented are transmembrane proteins. This makes sense given the changes required to go from marine environments to freshwater locations will often require changes in membrane structure, functionality and form when moving to contrasting osmotic regimes.

Some of these missing genes are easier to identify and more obviously functionally-linked than others. There are four genes with homology to nucleoporins in this list, OG0010707, OG0009642, OG0008744 and OG0010744. The homology is occasionally obscured by best blast hit name, where the best hit is named in an unclear or misleading fashion, generally by automatic annotation. Nucleoporins make up the nuclear pore complex, and regulate the flow of molecules between the cytoplasm and the nucleus. Such genes could have a role in cell control of osmolarity.

The loss of an overlapping (but not completely identical) gene set therefore seems to be a characteristic feature of freshwater-adapted species. It is important to note that loss of genes is not necessarily itself adaptive, but could be a consequence of these proteins falling into misuse. These genes are therefore excellent targets for functional investigations aimed at exploring why they might be used in marine conditions, but lost in freshwater.

#### 7.5 Cluster expansion and novel *Ephydatia muelleri* genes

*Identification of* Ephydatia muelleri *novelties*

The Orthofinder2 results reported in Supplementary Notes 7.2 and 7.3 are excellent ways of recording the expansion of gene families where these are shared between multiple species. However, if genes are unique to *E. muelleri,* or represent unique paralogs*,* they may be overlooked by the above analyses, particularly if they are single copy novelties with no genes of similar sequence. To detect these genes, we used an alternative approach, and sampled widely across metazoan phylogeny to ensure that *E. muelleri* novelties were correctly curated.

The same 38 species-strong dataset used for phylogenetic reconstruction in Supplementary Note 1.3 was used for predicting the occurrence of unique genes within homologous gene families. Full proteomes were blasted against each other using BLASTp in DIAMOND v0.9.31 [52], with cutoff e-value of 1e-5. Homomcl [14] was used to convert blast output for further use with the Markov clustering algorithm (MCL) [62]. The output clusters were used for phylogenetic reconstruction as seen in Supplementary Note 1.3.

A total of 85,317 clusters of homologues were predicted (putative homologous gene families). From these, 6,335 species-unique clusters for *Ephydatia muelleri* were identified, containing 18,240 of *Ephydatia muelleri*'s 39,245 genes. Please note that as different search mechanisms and cut-offs for identity have been used in this analysis, these do not match the assignations found in Supplementary Notes 7.1-7.3. We also determined which genes represented likely Porifera and Demospongiae novelties using the same process as found above, as well as determining which homogroups were not found in any demosponges at all. These four categories (All Porifera Unique, *Ephydatia muelleri* unique, Demospongiae Unique, and Sponges Not Demospongiae (*Oscarella carmela* and *Sycon ciliatum*) Unique) were then analysed for content, to determine whether the makeup of these categories of unique genes was linked to the adaptations of these clades (Supplementary Table 15).

These species- or clade-unique clusters were annotated for GO terms using two approaches to raise accuracy and cover more options (different tools tend to show slightly different predictions for GO terms). First, we used the PANNZER web tool [82]. The output was downloaded and analyzed using a custom bash commands. For *Ephydatia muelleri,* the annotation process mapped 14,095 of the 18,240 genes to GO terms (Supplementary Data 7: 7A). Any cluster that received a GO term is sufficiently close in sequence identity to genes of known homology to receive an annotation, but sufficiently different in sequence identity to them that it is not clustered into combined groups by the MCL algorithm. Genes not mapped to a GO category and not clustered by MCL are likely to be true novelties to *Ephydatia muelleri*, 4,145 genes in this case, but determining their function and evolutionary history will require detailed analysis and is beyond the scope of this work. This analysis was repeated for all 4 categories under analysis, for all 3 high level GO categories (Biological Process = BP, Cellular Components = CC, Molecular Function =MF).

The three different categories were summarized and visualized using the REVIGO web tool (http://revigo.irb.hr/) and R studio v-3.5.3 (Rstudio.com) to create REVIGO Gene Ontology treemaps, plotting the different hits into higher relevant categories and showing the corresponding size (calculated dispensability value) from the total annotation per group of hits (CC, MF or BP). These results are shown in Supplementary Figure 23 below and overleaf.

Secondly, we used the same PANNZER2 tool this time to predict GO terms overlap for each of the four categories indicated before. These results are shown in Supplementary Figure 22 below. Those results indicate that while by granularity Porifera was found to share 550 *Porifera unique* genes with 2,022 GO terms, only 197 GO terms were shared by all five species and 205 GO terms were found to be *E. muelleri* hits (Supplementary Figure 24, Supplementary Data 7: 7B). Among the three Demospongiae included in our analysis 3,303 GO terms were found to be shared, and for the non-Demospongiae, 858 GO terms are shared; the full list of GO term names can be found in Supplementary Data 7: T7B and 7C.

Because the homologous groups predicted can contain orthologues, we analyzed our sequences from the predicted clusters for each category by providing the data to OrthoVenn2 web tool [83]. The results are shown in Supplementary Figure 25 and Supplementary Data 7: Tables 7A-E. Those results indicate that despite many GO terms found to be unique in sequence terms, the analysis of orthology detection in the web tool used classified many of the genes to shared clusters. The *Porifera unique* genes contained only seven clusters of orthologues unique to *E. muelleri* and 42 clusters of orthologues shared among all the five species (Supplementary Figure 25D). Demospongiae were found to share 774 orthologous clusters, but only 30 genes were found with Swiss-Prot annotations (Supplementary Figure 25 E). The full list of GO terms and Swiss-Prot annotations for those analysis is shown in Supplementary Data 7: Table 7B.

While there is a wealth of information in these GO categories, the most striking result is how little they overlap in content. Each of the groups has a distinct pattern of GO terms that reflects a very different set of unique genes for each clade. Concentrating on *Ephydatia muelleri,* within the genes noted as unique in this species, the most prominent MF categories are linked to catalytic activity (GO:0003824), binding (GO:0005488) and transferase activity (GO:0016740), the most prominent CC category (representing half of all CC terms) is intracellular components GO:0005622, and the most obvious BP categories are linked to biological regulation (GO:0065007) and response to stimulus (GO:0050896). Among the small categories of MF for *Ephydatia muelleri* unique can be found calcium ion binding (GO:0005509), potassium ion binding (GO:0030955) and copper ion binding (GO:0005507). To have a closer look at the *E. muelleri* unique GO terms over all, we made a summary of all GO terms during all the analysis above and found 2,196 hits. The results can be found in Supplementary Data 7: 7E-7H. The highly represented GO terms are transportation and regulation related. These illustrate specific ways in which *E. muelleri* is expanding its gene complement - while GO categories are illustrative of broad-scale patterns, *E. muelleri* is expanding families of genes which allow it to identify and respond to changes in external stimulus, in ways that differ from other clades.

**Supplementary Figure 24:** REVIGO plots of GO terms found to be over-represented in four categories of genes (A: All Porifera Unique, B: *Ephydatia muelleri* unique, C: Demospongiae Unique, and D: Not Demospongiae Sponge Unique (*Oscarella carmela* + *Sycon ciliatum)*). Parentheses indicate the number of GO terms found for each of the groups in the category (MF/CC/BP).

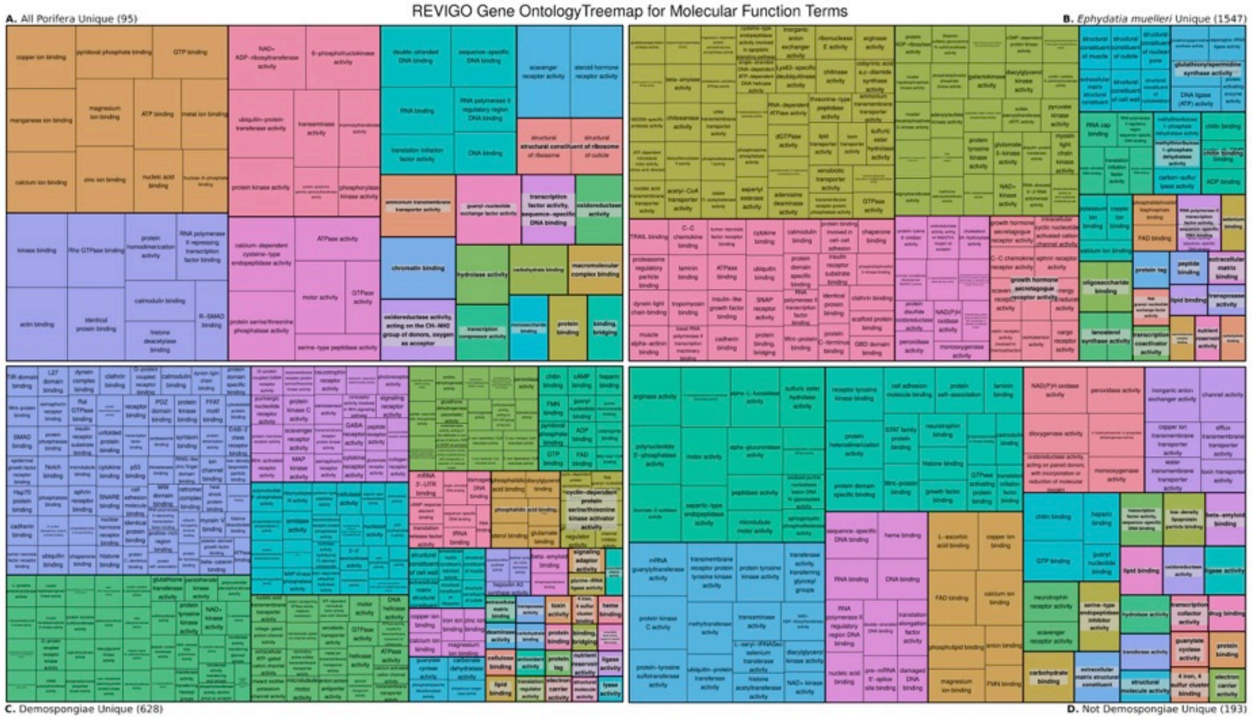

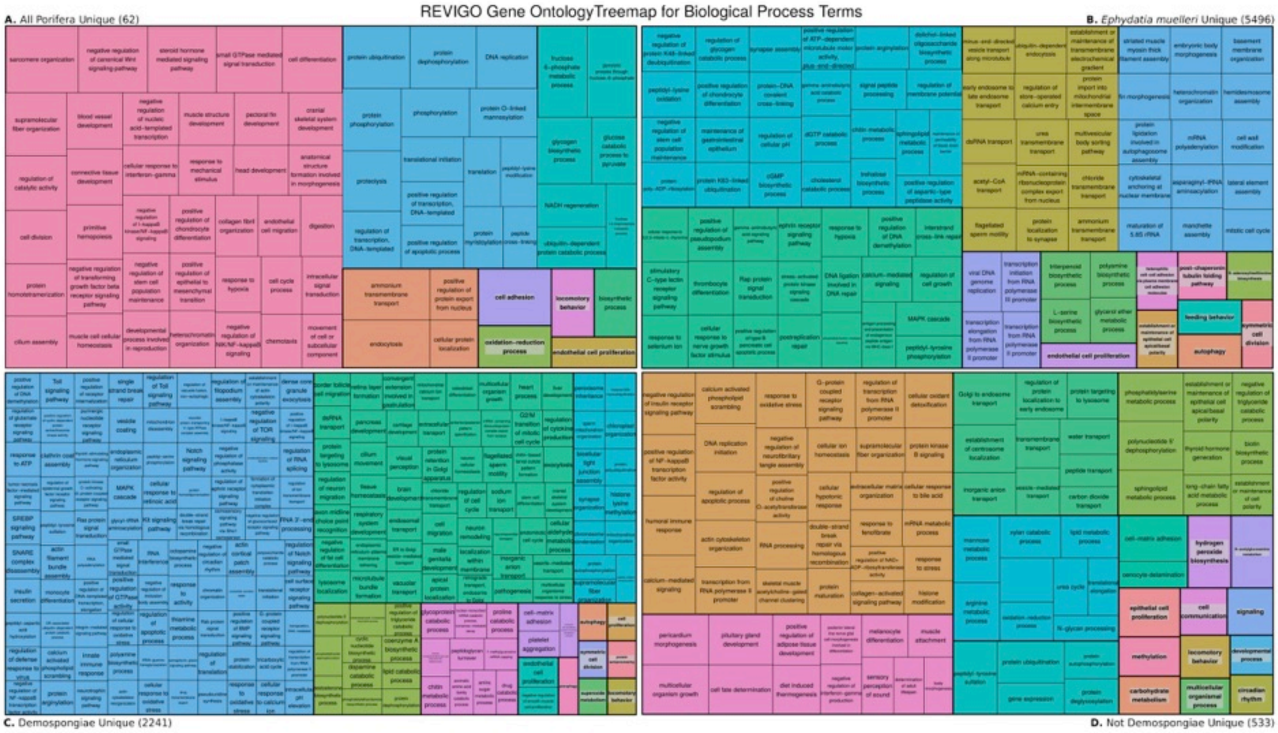
REVIGO Gene OntologyTreemap for Biological Process Terms
A. All Porifera Unique (62)
B. Ephydatia muelleri Unique (5496)
C. Demospongiae Unique (2241)
D. Not Demospongiae Unique (533)

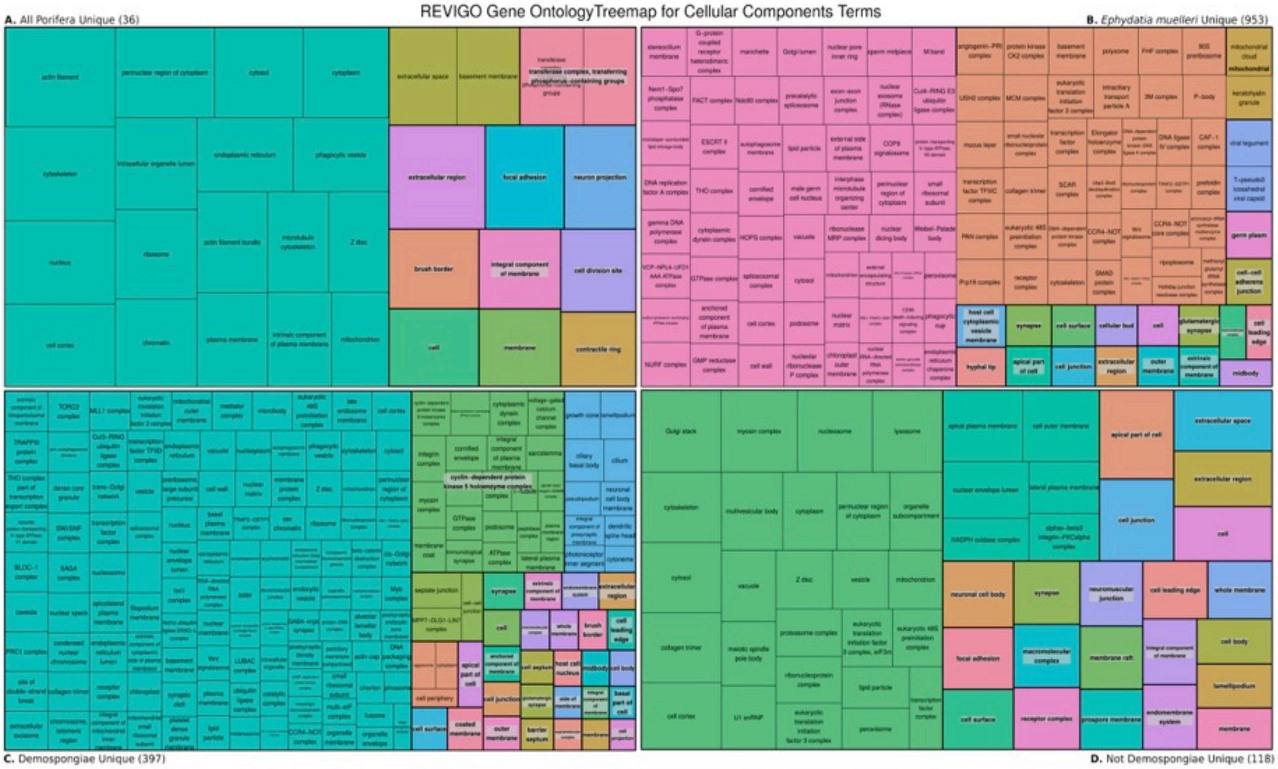
REVIGO Gene OntologyTreemap for Cellular Components Terms
A. All Porifera Unique (36)
B. Ephydatia muelleri Unique (953)
C. Demospongiae Unique (397)
D. Not Demospongiae Unique (118)

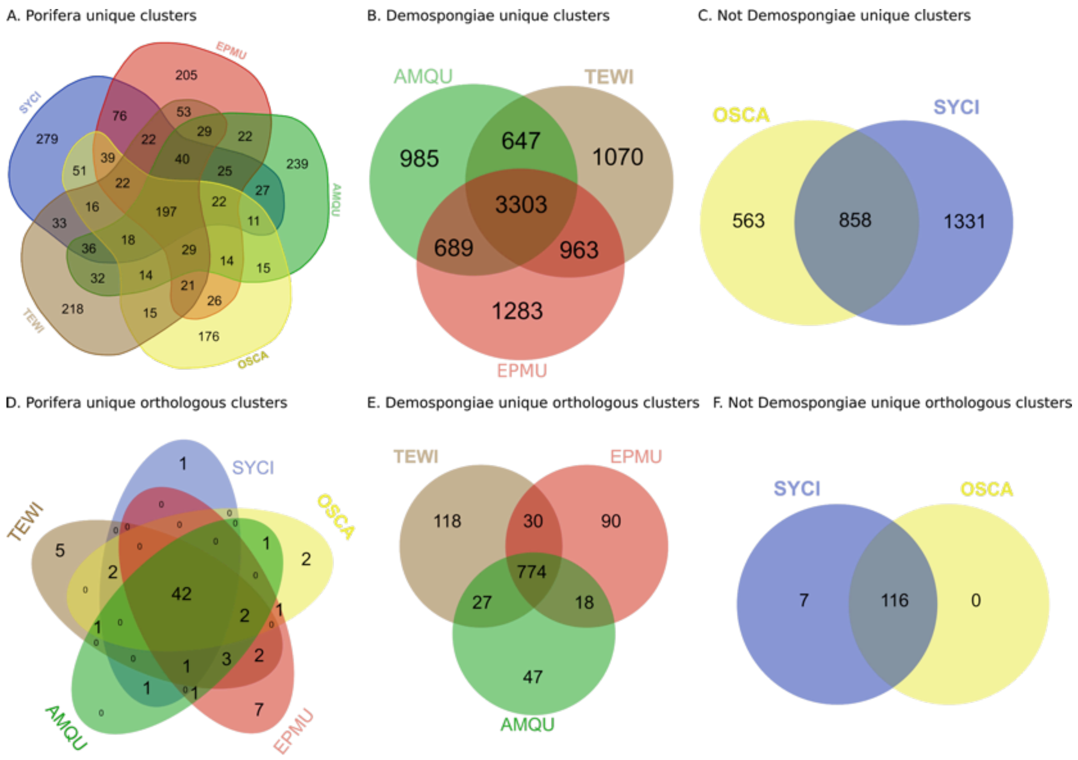

**Supplementary Figure 25:** Venn diagram plots showing representation for three categories analyzed for GO terms A) All Porifera Unique, B) Demospongiae Unique, and C) Not Demospongiae Sponge Unique (*Oscarella carmela* + *Sycon ciliatum))* and OrthoVenn2 clusters annotations of orthologous prediction for the same categories as A,B and C; D,E and F (respectively). Abbreviations: AMQU: *Amphimedon queenslandica*, EPMU: *Ephydatia muelleri*, TEWI: *Tethya wilhelma*, SYCI: *Sycon ciliatum*, OSCA: *Oscarella carmela*.

*Cluster expansion of positively selected gene sets*

In Supplementary Note 7.3 we noted 117 genes as showing evidence of being under positive selection in *Ephydatia muelleri.* We identified these genes in the MCL clusters described above, and used them to understand whether the copy number of these genes in particular had changed. If these genes were under selection following duplication, this could be because they are under decreased functional constraint. Similarly, if a gene proves particularly useful in freshwater conditions, it may be more likely to duplicate, both to increase transcriptional capacity, but also to act as a source of variation for future evolutionary changes.

The 117 genes under positive selection were mapped to 104 clusters, some clusters with more than one gene represented in the cluster (NB, some genes cluster in the same homology group with the granularity level used, see Supplementary Table 15). Using UGENE v.2 [84] we aligned these clusters with

MAFFT 7.450 [43] and exported the alignment as a distance matrix (by similarity %), with further details in Supplementary Data 7. These matrices were manually reviewed, and where *Ephydatia muelleri* sequences had higher similarity to one another than to any other sequences, these were noted as in-paralogues. Out of 117 genes with positive selection, 13 in total, can be marked as possibly selected and duplicated. Of the 104 clusters, 10 contain apparent in-paralogs, with three clusters containing two or more independent in-paralogous clusters of *E. muelleri* genes. Nine more duplicated genes of *Ephydatia muelleri* were found in those clusters, but were not identified as duplications of the genes with positive selection themselves. Among the 13 duplicated genes, five are part of the genes identified in the unique *Ephydatia muelleri*'s 18,240 genes: Em0001g3814a (MF;GO:0007166:cell surface receptor signaling pathway), Em0007g315a (MF;GO:0008270:zinc ion binding), Em0214g3a (MF;GO:0003676:nucleic acid binding), Em0001g241a (no PANNZER2 annotations with GO term or OrthoVenn2) and Em0020g955a (MF;GO:0015131,oxaloacetate transmembrane transporter activity). Three other genes found as positively selected in *Ephydatia muelleri* are in the list of genes unique to Demospongiae: Em0019g427a (Swiss;Q54P77: Probable 4-coumarate--CoA ligase 1, GO:0009698, P:phenylpropanoid metabolic process), Em0003g682a (Swiss; O46680: TGF-beta receptor type-1, GO:0003222, P:ventricular trabecula myocardium morphogenesis) and Em0014g783a (Swiss; Q5R8J8: DnaJ homolog subfamily B member 4 , GO:0006457, P:protein folding), Supplementary Data 7: 7I.

On average, there are 4.8 *Ephydatia muelleri* in-paralogs in these clusters (Median = 4, Mode =2). Genes found to be under selection therefore tend to be found in clusters which are prone to duplication. The full details of these clusters can be found in Supplementary Data 7: 7F.

**Supplementary Table 15:** Summary of the numbers of genes found in homologous cluster analysis. *Genes in total. Note that this clustering is independent of orthofinder-based clustering, and will more discretely cluster paralogs.

| **Unique homologous clusters of genes** | **Genes in cluster** | **In Supplementary Note** |
|---|---|---|
| Unique to Porifera | 57,916 | 7.5 Identification of novel *Ephydatia muelleri* genes |
| Unique to Demospongiae | 7,023 | 7.5 Identification of novel *Ephydatia muelleri* genes |
| Unique Demospongiae & positively selected | 3* | 7.5 Cluster expansion in positively selected genes |
| Unique to *Ephydatia* | 18,240 | 7.5 Identification of novel E*phydatia muelleri* genes |
| Unique Emu & positively selected | 5* | 7.5 Cluster expansion in positively selected genes |
| Unique to *Amphimedon* | 18,001 | - |
| Unique to *Tethya* | 4,299 | - |

#### 7.5.1 Specific gene cluster expansions: mGABA receptors

Metabotropic GABA receptors are one of 15 expanded gene clusters with over 120 predicted mGABA receptors in the *Ephydatia muelleri* genome (Supplementary Figures 26, 27). In detail, there are 127 mGABA receptor genes in 31 scaffolds, but 28% of the genes appeared in 3 scaffolds (scaffold 4 (14 genes), scaffold 13 (with 26 genes), and scaffold 22 (with 21 genes)). The rest of the scaffolds have from 1 to 8 genes, but most just have 1 or 2.

The sponge mGABAs align most closely with the GABA-B2 receptor of humans, and although the sequences show a shared conserved region in the LB1 portion of the fly trap domain, the different amino acids in the LB2 portion (Supplementary Figure 26) suggests that, if the proteins encoded by these genes are expressed like other mGABA-B2 receptors, then they likely bind different molecules than GABA. In freshwater there is an abundance of organic acids and so an exploration of what organic acids or other molecules potentially produced by microbes or other organisms in the water could bind these receptors would be a good next step.

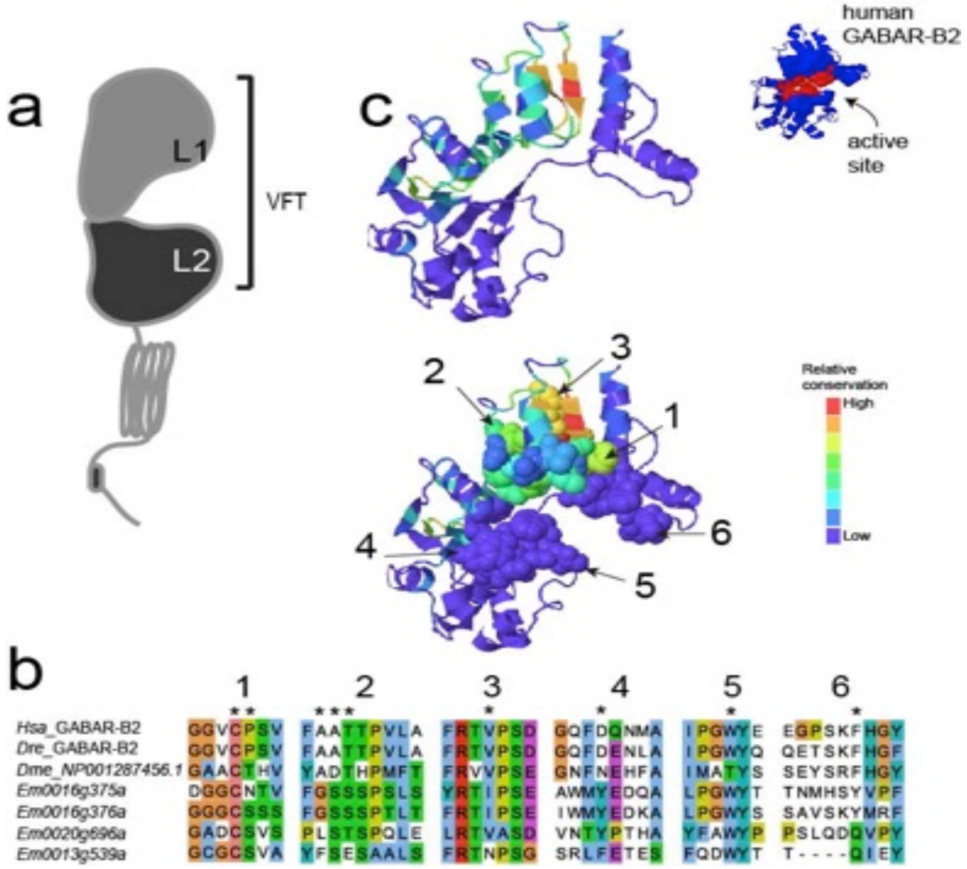

**Supplementary Figure 26**: Diagram of the mGABA receptor molecule showing the venus flytrap domain A) and the alignment of four of the *Ephydatia muelleri* mGABA-R sequences with human, zebrafish and fruit fly mGABA-B2 B). A phyre2 model of the human (inset) and *Ephydatia muelleri* mGABA-R showing the location of the conserved residues in the VFT domain.

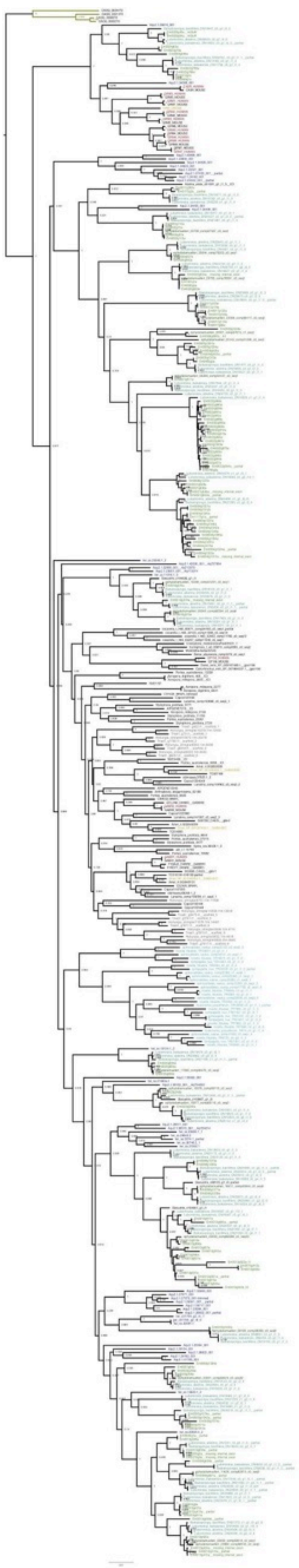

**Supplementary Figure 27:** Phylogenetic tree of mGABA receptors and related receptors across metazoans. Alignment was generated with MAFFT v7.450 (with option -linsi). Phylogenetic tree was generated using IQTREE v1.6.12 with the model WAG+F+R9 and 1000 pseudo-bootstrap replicates (-m WAG+F+R9 -bb 1000) . NB full size phylogeny available from Ephybase (https://spaces.facsci.ualberta.ca/ephybase/) in high resolution figure file download section.

#### 7.5.2 Specific gene cluster expansions: Aquaporins

Aquaporins are one of the best studied family of proteins involved in freshwater adaptation. Aquaporins are water and solute carriers that are present in all living organisms, and are involved in regulating osmotic change [85].They are divided into two main groups: one allowing the passage of water, ammonia and urea (AQP group), and the other group allowing glycerol, arsenite, and silicic acid among other compounds, usually called aquaglyceroporins or GLP group [85].We screened the genome of *Ephydatia muelleri* for aquaporin orthologues, and we aligned them with 131 sequences of diverse aquaporins from bacteria, protists, plants, fungi, and animals (see accession numbers on the figure between brackets) (Supplementary Figure 28) with MAFFT 7.450 [43]. Then we built our phylogenetic hypothesis for aquaporin evolution with RAxML 8.1.22 [86] with GTR-GAMMA as parameters of model evolution and 100 replicates for bootstrap sampling.

Whereas marine sponges have both types of aquaporin (Supplementary Figure 28), freshwater sponges only have aquaglyceroporins, a group with affinity to aquaporin 3 and 9 that allows intake of silicic acid [87] and another group, GLFPs, with high affinity to glycerol uptake facilitator proteins, which were previously only found in bacteria and plants (Supplementary Figure 28 [85, 88, 89]). The genome of *E. muelleri* has 9 paralogs of GLFP, with five of them located on the same scaffold (Em0009), and four of aquaporin 9 (Supplementary Figure 28). Orthologues of this GLFP are also present in marine sponges, including *Amphimedon queenslandica* [90] and other demosponges (Supplementary Figure 28). An interesting question is whether, as in plants [88], the presence of GLFPs in sponges may have happened via horizontal gene transfer.

Freshwater sponges have two rounds of duplication of AQU9 (Supplementary Figure 28). This gene encodes for a channel protein which is a highly permeable to glycerol and urea and has low permeability to water [91] . In contrast the closely related aquaporins 3 and 7 have a much higher permeability to water [91]. Phylogenetic analysis shows that sponge aquaglyceroporins are more similar to AQU9 than to AQU3 or AQU7, but their permeability ranges have never been tested. In freshwater systems, osmoregulation involves active ion uptake to compensate for passive ion loss to and water load from the environment [92]. In fishes, AQU3 helps re-establish osmotic balance in cells during acclimation to freshwater (e.g.[92,93]). Freshwater sponges have lost AQU8, which is used in vertebrates to compensate for different salinities (Supplementary Figure 28). Interestingly, AQU9 mediates silicon influx in the intestine of mice [87], and it would be interesting to explore whether it has a role in uptake of silicon in sponges. In sum, the differences in aquaporin complements between marine and freshwater sponges could account for the variability of the regulation of water homeostasis in such different environments, and highlights the use of *Ephydatia muelleri* as a useful model for studying molecular adaptations of marine invertebrates to freshwater ecosystems.

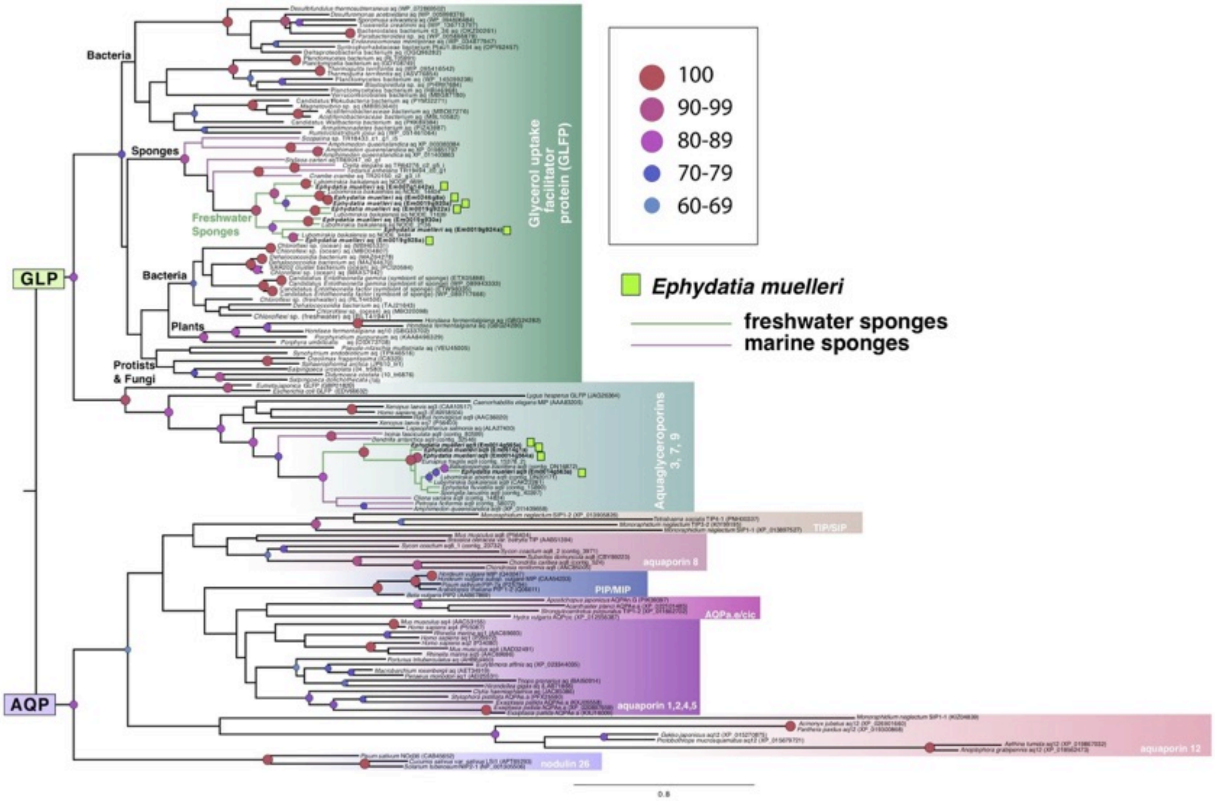

**Supplementary Figure 28:** Aquaporin (AQP) and aquaglyceroporin (GLP) phylogeny. Phylogenetic hypothesis generated was generated in RaxML with 100 bootstrap replicates (proportions given at base of nodes and colour coded following scale). *E. muelleri* sequences are indicated with green squares, freshwater sponges are indicated with green branches and marine sponges with blue branches.

#### 7.5.3 Analysis of Qbc-SNAREs in Holozoa

The high-quality genomic sequence data for *Ephydatia muelleri* reported here will support further investigation of early evolution of metazoan tissues via phylogenetic analysis of genes with potential tissue-specific functions. For example, one family of genes with tissue-specific functions are the SyNaptosomal-Associated Proteins (SNAPs) of vertebrates. These proteins are members of the Soluble N-ethylmaleimide sensitive factor Attachment protein REceptor protein superfamily [94,95]. These SNAP SNAREs function in membrane fusion at the cell surface. In mammalian cells, SNAP-25 mediates fusion of vesicles with the presynaptic membrane of neurons, while its paralogue SNAP-23 mediates fusion of vesicles in other regions of the cell surface [96,97,98]. The presence of genes encoding SNAP-25-like SNARE proteins has sometimes been inferred to indicate presence of neuron-specific protein machinery [35, 99]. However, homologues of SNAPs are widely conserved among eukaryotes without nervous systems, such as plants [100]. This raises the question of whether or not particular SNAP SNARE genes found in early-branching metazoan lineages indicate an early origin neuron-specific protein machinery. This question may be addressed via phylogenetic analysis to determine whether particular genes are orthologous to neuron-specific SNAP SNARE paralogues of other metazoans, and to investigate the timing of the origin of neuron-specific SNAP SNAREs found in mammalian cells.

**Methods**:

*Taxon selection*

As for other phylogenetic analyses reported, a representative set of holozoan genomes were selected for analysis in addition to *E. muelleri* predicted proteins and transcripts (see Supplementary Data 8A for summary of data sources).

*Similarity searching*

To identify homologues of the human SNAP SNAREs SNAP-23, 25, 29, and 47 among sampled genomic sequence data, we first aligned peptide sequences of representative metazoan homologues using MUSCLE [101]. Using HMMER v3.3 [102], this alignment was used to construct a hidden markov model and search for similar peptide sequences (http://hmmer.org). Only HMMER hits retrieved with an E-value less than or equal to 0.0005 were considered for downstream analysis. To exclude other types of SNARE proteins, a reciprocal-best-hit search strategy was used: Each hit sequence retrieved using HMMER was used as a query to search human peptide sequences with NCBI's Basic Local Alignment Search Tool. In each case, the subsequence of each hit showing similarity to the original query was used as a reciprocal search query, and only those retrieving SNAP-23, 25, 29, or 47 with an E-value less than or equal to 0.05 and five orders of magnitude less than any other human proteins retrieved were included for further analysis [103].

Also, several criteria were applied to identify representative sequences for phylogenetic analysis: 1) at least 55 amino acids in length or 15% of the length of the query; 2) no more than 98% identity with any higher-ranking hit sequence retrieved from the same genome; 3) overlap with a higher-ranking hit sequence such that at least 50 residues are aligned and are similar between the two sequences. These criteria were applied, and the similarity searches were run, using scripts similar to those described previously [104]. The source code for these scripts is available here: https://github.com/laelbarlow/amoebae.

*Phylogenetic analysis*

Identified representative amino acid sequences were aligned using MUSCLE 3.8.31 [101]. The resulting alignment was trimmed to include only those positions showing clear shared homology between the majority of sequences. Selection of a model of sequence evolution was performed using ModelFinder

[105]. Phylogenetic analysis was performed using IQ-TREE v1.6.12 for Maximum Likelihood analysis with nonparametric bootstrapping and MrBayes 3.2 for Bayesian analysis [45, 106]. MrBayes was run with four Markov chains with a sample frequency of 1000 until the average standard deviation of split frequencies reached 0.01, and a burnin fraction of 25% was applied for summarizing results. Analyses were run on the CIPRES webserver [107]. Several rounds of analysis were performed to remove redundant sequences and to identify orthologues of SNAP-23 and 25, excluding sequences more closely related to SNAP-29 or 47.

Similarity searches retrieved many homologous sequences in the sampled genomes (see Table 2 in Supplementary Data 8B for summary of all search results). Phylogenetic analysis of identified representative sequences allowed identification of those holozoan sequences most closely related to the human orthologues of SNAP-23, 25, 29, and 47 (not shown). Phylogenetic analysis of SNAP-23/25 homologues revealed that the vertebrate neuron-specific paralogue SNAP-25 arose from a duplication that occurred in an early vertebrate, while the two SNAP-23/25-like genes found in *E. muelleri* arose from an independent duplication that occurred in sponges (Supplementary Figure 29).

Homologues of the vertebrate neuron-specific SNARE SNAP-25 were identified in *E. muelleri* and other Porifera (Supplementary Figure 29). However, like the non-holozoan homologues, these genes are no more closely related to SNAP-25 than to the non-neuronal vertebrate paralogue SNAP-23. Therefore, the presence of these SNAP-25-like genes does not indicate the presence of neural-specific protein machinery, and is not suggestive of a capacity for synaptic transmission in Porifera.

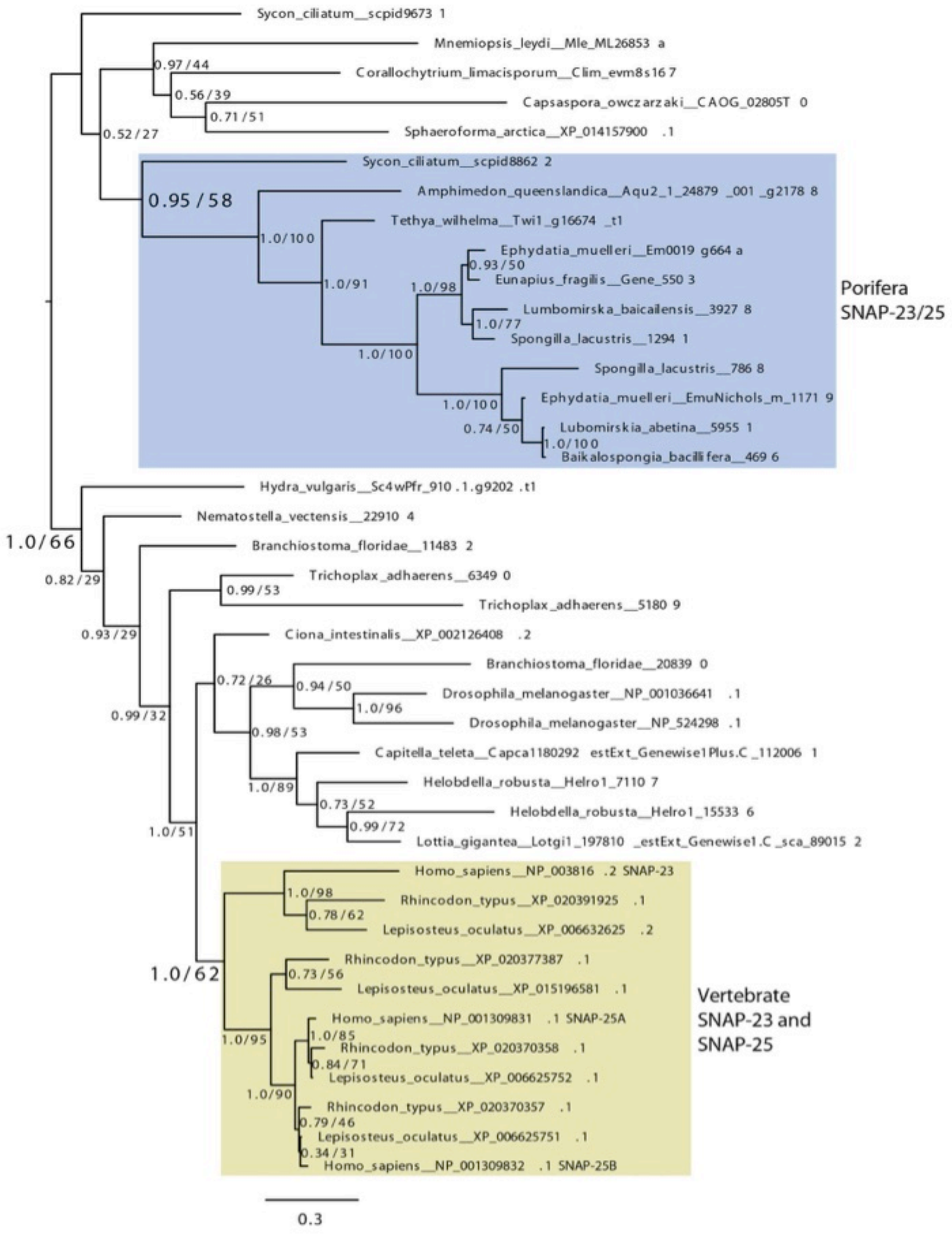

**Supplementary Figure 29:** Phylogenetic analysis of representative holozoan SNAP-23/25 related proteins. Bayesian analysis was performed with MrBayes, and Maximum Likelihood analysis with non-parametric bootstrapping was performed using IQ-TREE [45, 106]. The LG substitution matrix was used with gamma distributed rates using four discrete rate categories. The topology recovered by MrBayes is shown, and support values for each internal node are shown in the order MrBayes/IQ-TREE, where MrBayes supports are posterior probabilities (0.80 or greater is considered significant) and IQ-TREE bootstrap percentages (50 or greater is considered significant).

### Supplementary Note 8: Developmental RNAseq in *Ephydatia muelleri*

#### 8.1 RNAseq Methods

To understand which expressed genes are common to and which are distinct from other metazoans during the development of the filter-feeding body plan, we examined differential gene expression from hatching gemmules through to the formation of a filtering sponge. Five distinct stages in development of a freshwater sponge from the gemmule can be observed: Stage 1, differentiation of amoebocytes from thesocytes in the gemmule and emergence of the first amoebocytes with the first steps in formation of an epithelium to cover the differentiating cells; Stage 2, formation of choanocyte chambers, spicules, and the first lacunae or nascent canal structures; Stage 3, multiplication of choanocyte chambers to surround large lacunae that spread over a larger area; Stage 4, merging of lacunae into clear excurrent canals, and formation of an osculum; Stage 5, organization of the canals with a greater appearance of polarity, enlarged at the base of the osculum, and a more substantial osculum. Because the latter two stages are distinguished only by the reorganization of canals into their final shape, we considered these as one stage in our analysis. More detail on these stages is available in Supplementary Note 11.

These stages have been analysed by others previously in related species (e.g. *Ephydatia fluviatilis*) [108]. We find, as have other authors [108] that timing of development through these stages can vary with temperature, how long the gemmules have been dormant for, and the individual sponge from which the gemmules were obtained. Other factors may play a role in variability in hatching and timing of development, such as size of the gemmule (e.g. how many cells it contains), presence of algal symbionts, the type of culture medium hatched in (there are several recipes for freshwater media), exposure to light, and frequency with which the medium is changed, which alters the likely available oxygen and production of wastes by gemmules. Crowding of gemmules in a dish, or size of dish relative to the number of gemmules can also influence development. Most laboratories hatch gemmules at 'room temperature' in the dark, and change the culture medium every two days.

Gemmules from three individuals stored in 10% DMSO at -80ºC for 1-5 years were thawed and hatched. For Stage 1, gemmules were allowed to develop for only 12 hours before tissue was harvested for RNA. For Stages 2, 3, and 5, gemmules were allowed to hatch and sponges cultured in the lab as described [19]. Tissue was flash frozen in liquid nitrogen and either stored at -80ºC or processed immediately for RNA. Total RNA was extracted using the animal tissue RNA purification kit (Norgen Biotek, Thorold, Ontario, Canada) following the manufacturer's protocol. Total RNA quantity and purity were analysed by nanodrop, Qubit and Bioanalyzer 2100 (Agilent, CA, USA). RNA with RIN number >8.9 was stored as a precipitate in NaOH and ethanol.

Poly(A) selection and library preparation from mRNA was carried out by LC Sciences (Houston, Tx) as follows. Poly(A) RNA integrity was checked with Agilent Technologies 2100 Bioanalyzer. Poly(A) tail-containing mRNAs were purified using oligo-(dT) magnetic beads (Invitrogen) with two rounds of purification. After purification, poly(A) RNA was fragmented using divalent cation buffer in elevated temperature. RNA fragments were reverse-transcribed to create the final cDNA library in accordance with the standard protocol for the Truseq mRNA-Seq sample preparation kit (Illumina, San Diego, USA). the average insert size for the paired-end libraries was 300 bp (±50 bp). Quality control and quantification of the sequencing library were performed using Agilent Technologies 2100 Bioanalyzer High Sensitivity DNA Chip. Paired-ended sequencing was performed on an Illumina Hiseq 4000 sequencing system by LC Sciences. Cutadapt 1.10 [109] and proprietary perl scripts (LC Sciences) were used to remove reads containing adaptor contamination, low quality bases and undetermined bases. Sequence quality post-cleaning was verified using FastQC 0.10.1 (http://www.bioinformatics.babraham.ac.uk/projects/fastqc/).

HISAT 2.0 [110] was used to map RNAseq reads to the reference *E. muelleri* genome. StringTie 1.3 [111] and edgeR v3.14.0 [112] were used to estimate the expression levels (FPKM) of all transcripts across all replicate samples. mRNAs that had log2 (fold change) >1 or log2 (fold change) <-1 and with statistical significance ($p$-value < 0.05) were considered significantly differential mRNAs. Annotation was obtained using the gene IDs from the swissprot blast of the genome, and then automated through the Blast2GO PRO mapping and annotation pipeline [113] to obtain Gene Ontology (GO) terms and KEGG annotations. To understand the proportion of novel genes and those with eukaryotic origin, we used the blast annotation from swissprot and coded the genes as eukaryotic when the hits were from eukaryotic organisms (including sponges), sponge-specific when they only contained protein assignments to sponges, or *Ephydatia*-specific when they did not blast against any organism. Alternative splicing events were obtained using ASprofile [114].

**8.2 Expanded results of RNAseq**

*Sequencing and read cleaning*

The results per sample for our RNA sequencing can be seen in Supplementary Table 16 below, with results shown before and after read cleaning. Sequencing quality was excellent, with high (>98%) Q30% observed for all samples. The least well-recovered sample was Sp3St1_2, with 6.21 Gbp sequenced, and the most-sequenced sample, Sp2St3, contained 10.43 Gbp. In all cases, good sequencing depth was present for three samples per Stage.

**Supplementary Table 16:** Basic metrics related to RNAseq analysis

| Sample (Sp= specimen, St=stage) | Raw Data | | Valid Data, post-cleaning | | Q20% | Q30% | GC content % |
|---|---|---|---|---|---|---|---|
| | Reads | Bases | Reads | Bases | | | |
| Sp2St1 | 65384222 | 9.81G | 63424534 | 9.51G | 99.98 | 98.81 | 50 |
| Sp3St1_2 | 42657906 | 6.40G | 41381358 | 6.21G | 99.98 | 98.81 | 50 |
| Sp5St1a | 60690692 | 9.10G | 58317434 | 8.75G | 99.98 | 98.24 | 50.5 |
| Sp2St2 | 71413884 | 10.71G | 69402662 | 10.41G | 99.98 | 98.88 | 50 |
| Sp3St2 | 53437384 | 8.02G | 51954310 | 7.79G | 99.98 | 98.8 | 50 |
| Sp5St2 | 58653672 | 8.80G | 56964766 | 8.54G | 99.97 | 98.27 | 50 |
| Sp2St3 | 71617156 | 10.74G | 69513422 | 10.43G | 99.98 | 98.74 | 50 |
| Sp3St3 | 55205926 | 8.28G | 53676614 | 8.05G | 99.97 | 98.18 | 50 |
| Sp5St3 | 64282994 | 9.64G | 62852894 | 9.43G | 99.97 | 98.1 | 50 |
| Sp2St5 | 68103604 | 10.22G | 66444460 | 9.97G | 99.97 | 98.82 | 50 |
| Sp3St5 | 53605480 | 8.04G | 52574334 | 7.89G | 99.97 | 98 | 49.5 |
| Sp5St5 | 57330788 | 8.60G | 56142162 | 8.42G | 99.97 | 98.11 | 50 |

**Supplementary Table 17:** Statistics related to mapping of reads to reference genome.

| Sample | Valid reads | Mapped reads | Unique Mapped reads | Multi Mapped reads | PE Mapped reads |
|---|---|---|---|---|---|
| Sp2St1 | 63424534 | 54034771 (85.20%) | 34434234 (54.29%) | 19600537 (30.90%) | 48568276 (76.58%) |
| Sp3St1_2 | 41381358 | 31893750 (77.07%) | 21328580 (51.54%) | 10565170 (25.53%) | 28148154 (68.02%) |
| Sp5St1a | 58317434 | 46948931(80.51%) | 30730068 (52.69%) | 16218863 (27.81%) | 41844802 (71.75%) |
| Sp2St2 | 69402662 | 58016625 (83.59%) | 37537781 (54.09%) | 20478844 (29.51%) | 53095824 (76.50%) |
| Sp3St2 | 51954310 | 42725176 (82.24%) | 27783460 (53.48%) | 14941716 (28.76%) | 38787960 (74.66%) |
| Sp5St2 | 56964766 | 44732428 (78.53%) | 29459451 (51.72%) | 15272977 (26.81%) | 40000168 (70.22%) |
| Sp2St3 | 69513422 | 54350710 (78.19%) | 35560176 (51.16%) | 18790534 (27.03%) | 48554822 (69.85%) |
| Sp3St3 | 53676614 | 41986038 (78.22%) | 27498767 (51.23%) | 14487271 (26.99%) | 37583766 (70.02%) |
| Sp5St3 | 62852894 | 51631663 (82.15%) | 33202176 (52.83%) | 18429487 (29.32%) | 46929530 (74.67%) |
| Sp2St5 | 66444460 | 53430271 (80.41%) | 33999367 (51.17%) | 19430904 (29.24%) | 47806086 (71.95%) |
| Sp3St5 | 52574334 | 43332773 (82.42%) | 26966951 (51.29%) | 16365822 (31.13%) | 39488430 (75.11%) |
| Sp5St5 | 56142162 | 43070516 (76.72%) | 27828868 (49.57%) | 15241648 (27.15%) | 38223518 (68.08%) |

*Mapping to genome*

For all replicate samples, excellent mapping results were observed to our reference genome. In any sample, no fewer than 76.72% of all reads could be mapped to the genome (Supplementary Table 17). We were also able to note where these mapping results occurred relative to coding sequences in the genome (Supplementary Figure 30). No fewer than 90.09% of the reads mapped were placed in exonic regions in any sample. Between 1.36% and 2.82% of our RNAseq reads mapped intergenically. These reads could be mapping to genes not recovered in our gene models, however, they will also represent non-coding RNA contained stochastically in our libraries,

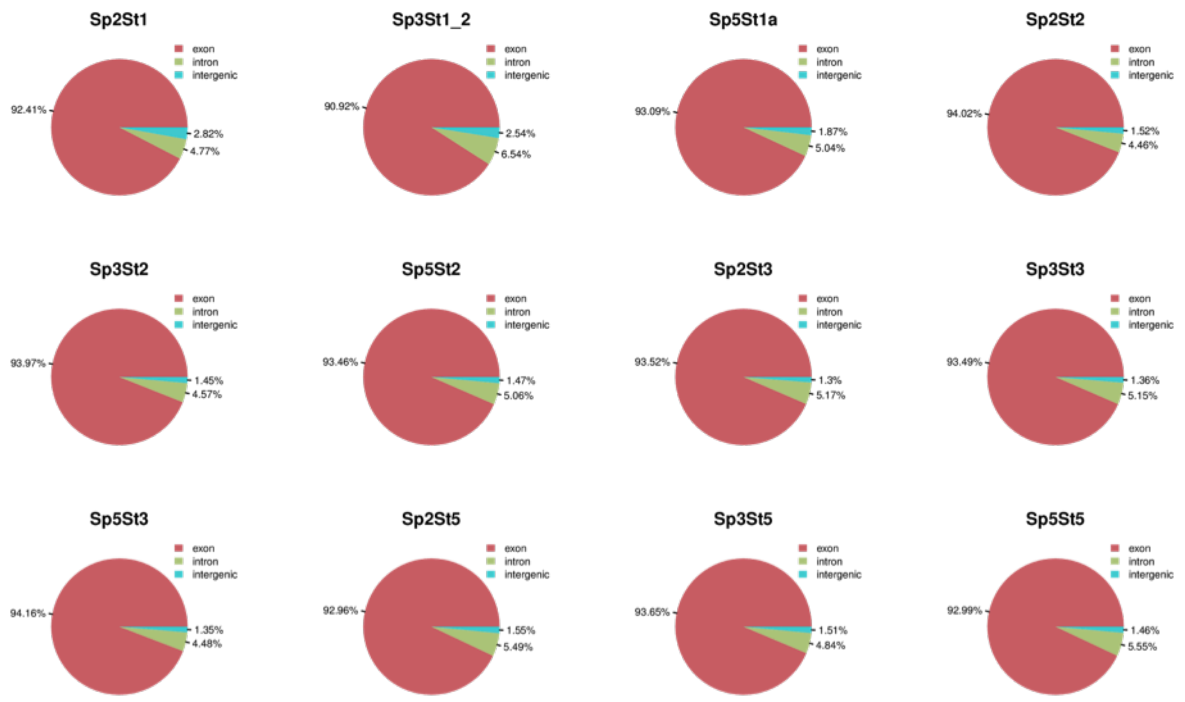

**Supplementary Figure 30:** Mapping locations of reads from each sample. Red (largest portion in all cases) shows exonic, green intronic, and blue intergenically mapped proportion of reads per sample, with percentages as indicated on figure.

*Differential gene expression, Gene Ontology and KEGG enrichment*

To understand the evolution and deployment of the aquiferous body plan in sponges, we examined differential gene expression from hatching gemmules through to the development of filtering sponges. Remarkably, 13,285 genes were differentially expressed across the gemmule-hatching process (Fig. 5, Supplementary Figure 31). Among the few genes differentially expressed during Stage 1 (pre-hatching), most were orphan genes or *Ephydatia*-specific and those genes that were known were involved in processes like stem cell proliferation (*E3 ubiquitin-protein ligase TRIM17*), oxidation-reduction (specifically arachidonate pathways for glycogen breakdown and arachidonate pathways for fatty acid metabolism), apoptosis and glutamine metabolism (Fig. 5 and Supplementary Figure 31). During Stages 2 and 3 when the canal system is becoming organized (Fig. 5 and Supplementary Figure 31), the genes *noggin-2* and *hedgling*, involved in developmental patterning [115,116] are upregulated, as well as genes involved in cell motility and adhesion such as *myosin regulatory light chain* and genes with roles in extracellular matrix formation developmental patterning including *collagen alpha 1* and *fibroblast growth factor receptor-like*.

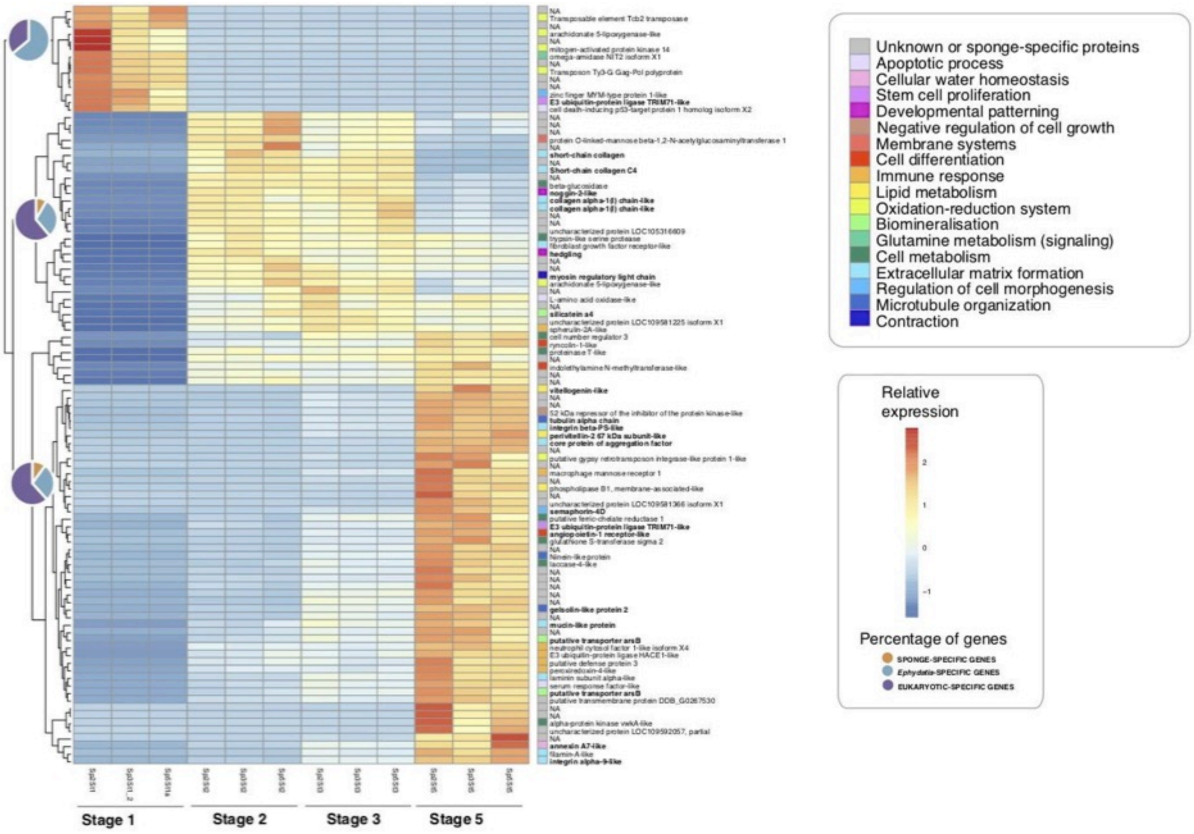

**Supplementary Figure 31:** Heatmap of the 100 most significantly differentially expressed genes in our RNAseq analysis. On the right, gene IDs and their affiliation to different biological categories (in colours). On the left, pie charts depicting percentage of differentially expressed genes with an eukaryote, sponge, or *Ephydatia* origin.

Sponge-specific genes, such as *silicatein,* involved in forming the silica-based sponge skeleton [117], were also expressed in Stages 2-3. In Stage 5, when the sponge has a fully formed aquiferous system including an osculum (Figure 5 and Supplementary Figure 31), many genes are significantly upregulated, and among those are those involved in immune response and genes involved in the formation of stable epithelia (e.g., *type IV collagen, spongin short-chain collagen C4*, *par3/6, contactin, scribble, MAGUK,* and *laminin*) (Figure 5 and Supplementary Figure 31). Interestingly, once the osculum is formed, and the sponge is unequivocally in contact with its surroundings by filtering the bacteria and other microbes present in the water, and deploys the machinery involved in activating the immune response, including *macrophage mannose receptor 1* (involved in host-antigen recognition [118]), *neutrophil cytosol factor 1* (involved in the production of Reactive Oxygen Species, ROS, in vertebrates [119]), and *peroxiredoxin 4* (also involved in oxidative stress [120]), among others (Fig. 5 and Supplementary Figure 31). Remarkably, one of the genes responsible for the formation of leakage-resistant capilliaries in humans [121], *angiopoietin 1*, is significantly overexpressed in Stage 5, when the canals of *Ephydatia muelleri* are being formed. The overexpression of angiogenetic machineries during aquiferous system formation in *Ephydatia* strongly suggest that this process is worth studying in detail to understand potential similarities between vessel and canal formation.

Among the most enriched Gene Ontology (GO) categories in the analysis, we found the Cellular Component categories of membrane (including integral component of membrane and plasma membrane), cytosol, extracellular exosome, and to a lesser extent, cytoskeleton (Supplementary Figure 32A). In regard to the Biological Process categories, the main enriched ones were related to oxidation-reduction processes (occurring mainly in Stage 1) and focal adhesion (Stages 2 and 3), and in Molecular Function, ATP binding, protein binding, and calcium ion binding were the most enriched (Supplementary Figure 32). Using KEGG pathways, we identified the Wnt signaling pathway and the arachidonic acid metabolism as the most enriched (Supplementary Figure 32B).

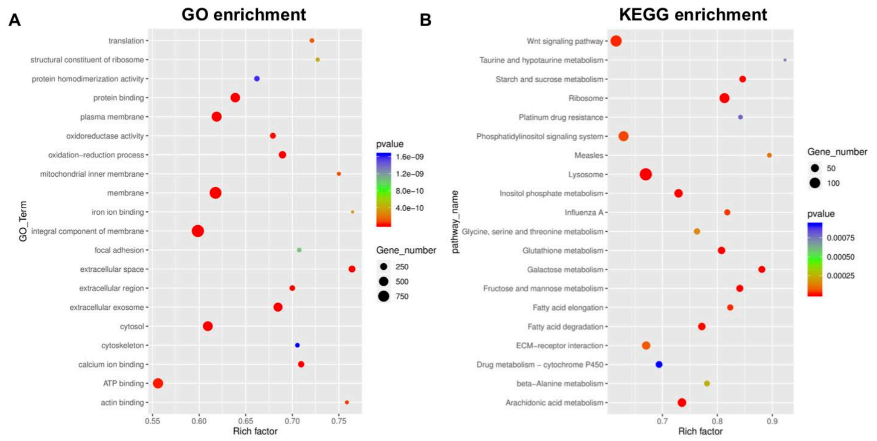

**Supplementary Figure 32:** A) GO enrichment and B) KEGG enrichment across the DGE analysis of the development of *Ephydatia muelleri*.

*Alternative splicing*

RNAseq analysis revealed a diversity of alternative splicing to be present in *E. muelleri.* We observe several different kinds of alternative splicing (Supplementary Figure 33) including: A) Exon skipping (SKIP) and cassette exons/multiple skipping (MSKIP), B) single intron retention of single (IR) and multiple intron retention (MIR), C) alternative exon ends (AE), D) alternative transcription start site (TSS), E) alternative transcription termination site (TTS), F) Approximate SKIP (XSKIP), G) Approximate MSKIP (XMKIP), H) Approximate IR(XIR), I) Approximate MIR(XMIR), and J) Approximate AE(XAE). These are plotted by frequency in each stage in Supplementary Figure 33.

The most common events of alternative splicing detected in our RNAseq experiment were alternative transcription start and termination sites (TSS and TTS), followed by IR and AE (Supplementary

Figure 33), as usually found in other organisms during different treatments or developmental stages (e.g.,[122]). However, in contrast with our results, marine sponges differentially use intron retention as the main mode of AS [50]. All samples showed relatively similar events of AS, but consistently samples at Stage 1 showed fewer AS events for all types (Supplementary Figure 33B). While the number of AS events (or spliced genes) is definitely correlated with the number of expressed genes and therefore likely to be less frequent in Stage 1 when the transcription has barely started, it could also reflect an evolutionary shift in the use of splice variants through development. In this sense, this supports the notion that AS might have evolved from mis-splicing as early metazoan cells evolved to use and benefit from multiple splicing outputs [123]. This new genome widens the possibilities of performing deep comparative genomic studies of the origins and differential use of alternative splicing at the onset of the acquisition of multicellularity.

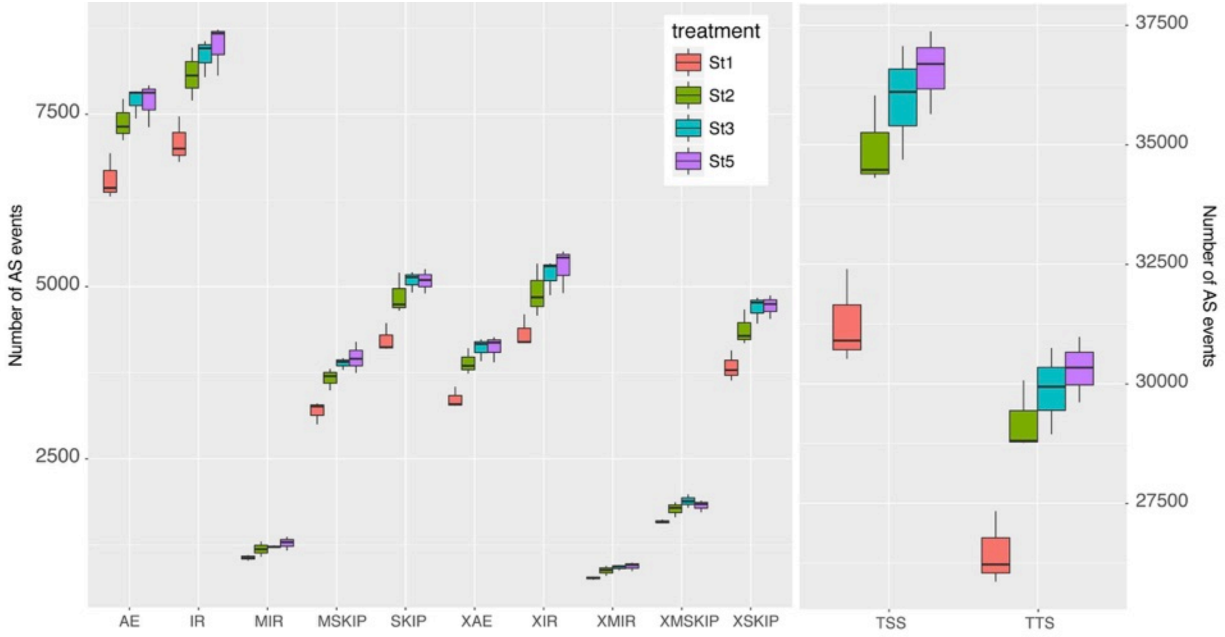

**Supplementary Figure 33:** Alternative splicing events occurring during the development of *Ephydatia muelleri.* Abbreviations as follows: SKIP, Exon skipping; MSKIP, cassette exons or multiple skipping; IR, single intron retention; MIR, multiple intron retention; AE, alternative exon ends; TSS, alternative transcription start site; TTS, alternative transcription termination site; XSKIP, Approximate SKIP; XMSKIP, Approximate MSKIP; XIR, Approximate IR; XMIR, Approximate MIR; and XAE, Approximate AE. Boxplot centre lines are medians, box limits are quartiles 1 (Q1) and 3 (Q3), whiskers are 1.5× interquartile range. 12 independent animal samples (3 per stage, Stages 1, 2, 3 and 5 as labelled) described in Section 8.1 were mapped to 39,245 *Ephydatia muelleri* genes, with results as shown here.

### Supplementary Note 9: Gene content in *Ephydatia muelleri*

We manually searched our *Ephydatia muelleri* gene models to annotate important gene families, including those involved in metazoan specific cellular signaling, chemical signaling. Epithelia, as well as cell and focal adhesion, innate immune function, germ-line, stem cell and sex determination genes. We also examined the regulatory machinery for RNA interference and the NuRD complex for recognizing methylated DNA and nucleosome remodeling. Manual searches and annotation was conducted as described in the Methods Supplementary Note, main text.

In some cases, putative orthologues appear to be highly divergent, however, in at least two cases, it has been shown that the homologues retain function. For example, while the methyl-cytosine binding domain (MBD) protein, MBD2/3, found in *E. muelleri* (Supplementary Table 18) appears to be highly divergent from human MBD2 and 3, it has been shown that *E. muelleri* MBD2/3 selectively binds methylated DNA, forms a coiled-coil interaction critical to recruitment of the NuRD complex, possesses a high degree of identity for residues needed for recognizing methylated cytosines, and binds with similar affinity to vertebrate MBDs [124].

Our manual annotation confirmed the completeness of our resource, and provides the starting point for a more detailed analysis of gene family presence and function in sponges and other non-bilaterian metazoans. As with our automated analyses, we note duplication in many genes.

#### 9.1 Ion channels

In terms of Ion Channels, we found at least five voltage gated calcium channels (with copies, two similar to L-type and 3 similar to 2-pore) and possibly one voltage gated K channel (remaining hits were for calcium channels). There were many gene hits to each of cyclic gated nucleotide channels and chloride channels and a ryanodine receptor. Interestingly absent in the *E. muelleri* genome are voltage gated sodium channels (all hits were identical to the calcium channels for which the e-values were stronger), epithelial sodium activated channels (ENaCs), Leak channels, as well as glutamate gated Ion channels (GICs). We found a large diversification of Transient Receptor Potential channels and because of their suspected role in sensing flow in the sponge [125] we proceeded with a more extensive analysis (below).

*Trp Channel Genes - Sequence collection*

We recovered sequences of experimentally characterized TRP channels from the HomologGene database at NCBI. Using CD-search [126], we located in each sequence the transmembrane domain (TM) region that includes the TRP domain. We used this portion as query to BLAST against various animal and choanoflagellate databases (non-redudant database at NCBI, proteomes from *N. vectensis, D. pulex, M. brevicollis* at JGI, proteome from *M. leidyi* at NHGRI, proteome from *P. bachei* at Neurobase and *P.*

*dumerilii* transcriptome at Jekely Lab MPI). We selected hits with an E value > 1 e-40 and trimmed them to include only the TM region. We used this set of sequences and the set of experimentally characterized sequences to search for TRP channel homologs in the E. muelleri genome (AUGUSTUS gene models), as well as in 16 published transcriptomes of sponges from the four main sponge clades as seen in Supplementary Table 18 (below). We translated *in silico* TBLASTN hits with an E value > 1 e-40 (alignment length > 400 aa) in all 6 possible frames and selected the required sequence for downstream phylogenetic analysis.

*Alignment and phylogenetic reconstruction*

We aligned the collected sequences with PROMALS3D [127] using as template resolved structures of TRPV (PDB ID: 5IWK), TRPN (PDB ID: 5VKQ), TRPM (PDB ID: 6BQR), TRPML (PDB ID: 5W3S), and TRPP2 (PDB ID: 5K47) channels. We used TrimAL v1.2 [44] with default parameters to select conserved regions of the alignment and filtered out sequences 90% or more redundant. We used the resulting alignment to reconstruct the maximum-likelihood phylogeny with the PhyML 3.0 algorithm [128]. The LG +G+F model was selected by the smart model selection tool [129] as the best fit for the data. We used SPR with 5 initial random trees to search for the tree topology. We used the aBayes scoring statistics [130] to evaluate the statistical support of the phylogeny, displayed in Supplementary Figure 34.

**Supplementary Table 18:** List of sponge species whose transcriptome was used to retrieve TRP channel homologues.

| Clade | Species | References |
|---|---|---|
| Demospongiae | *Amphimedon queenslandica* | (Fernandez-Valverde et al. 2015)[50] |
| | *Spongilla lacustris* | (Riesgo et al. 2014)[131] |
| | *Xestospongia testudinaria* | (Ryu et al. 2016)[132] |
| | *Stylissa carteri* | (Ryu et al. 2016)[132] |
| | *Ephydatia muelleri* | (Windsor Reid et al. 2018)[133] |
| | *Petrosia ficiformis* | (Riesgo et al. 2014)[131] |
| | *Ircinia fasciculata (syn:Sarcotragus fasciculatus)* | (Riesgo et al. 2014)[131] |
| | *Pseudospongosorites suberitoides* | (Riesgo et al. 2014)[131] |
| | *Chondrilla caribea* | (Riesgo et al. 2014)[131] |
| | *Crella elegans* | (Riesgo et al. 2012)[134] |
| | *Tethya wilhelma* | (Francis et al. 2017)[135] |
| Calcarea | *Leucosolenia complicata* | (Fortunato et al. 2014)[136] |
| | *Sycon ciliatum* | (Fortunato et al. 2014)[136] |
| Homoscleromorpha | *Corticium candelabrum* | (Riesgo et al. 2014)[131] |
| | *Oscarella carmela* | (Nichols et al. 2012)[137] |
| | *Oscarella sp.* | (Nichols et al. 2012)[137] |
| Hexactinellida | *Aphrocallistes vastus* | (Riesgo et al. 2014)[131] |

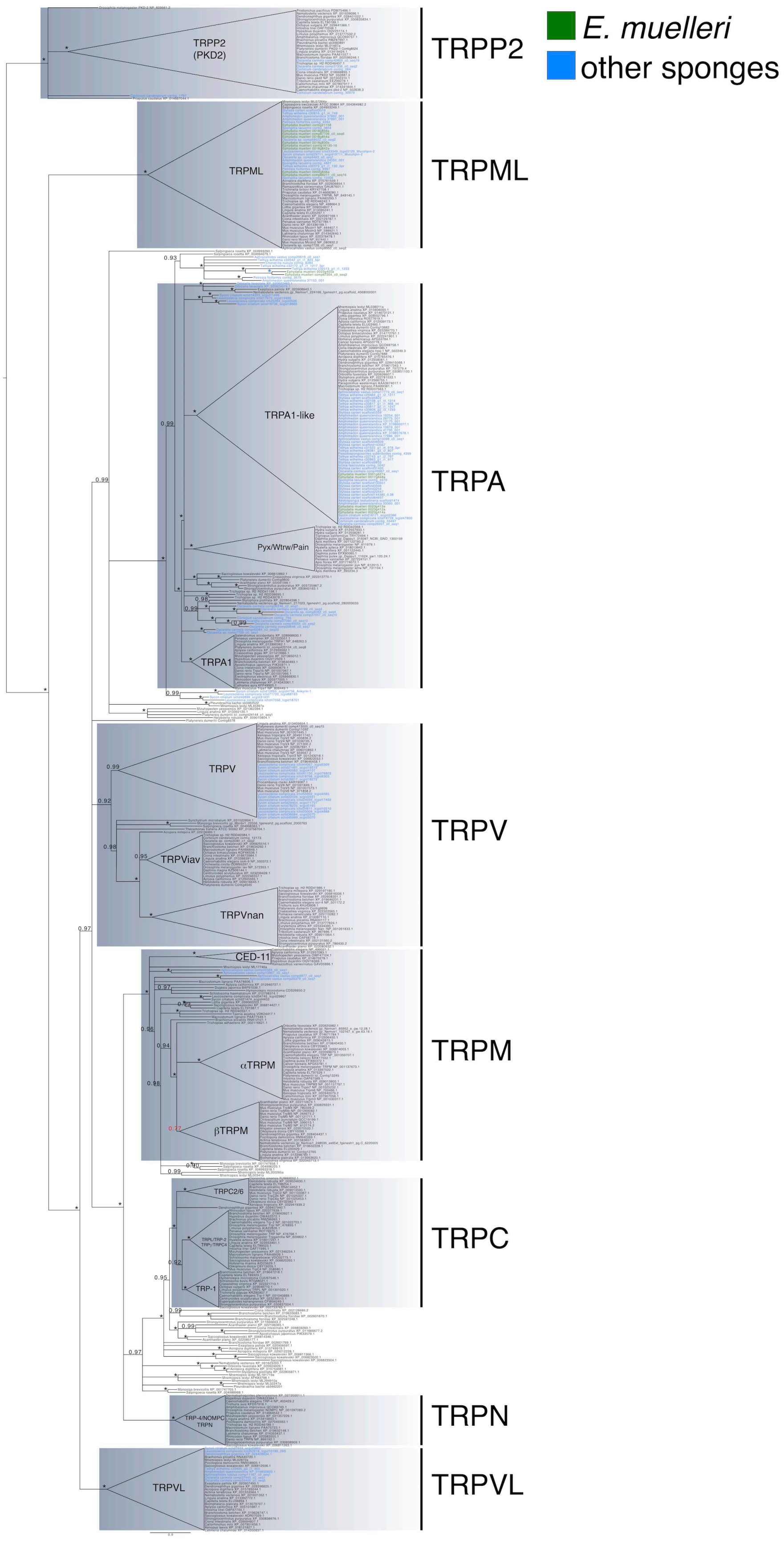

**Supplementary Figure 34:** Trp channel gene phylogeny. Sequences shown in green belong to *Ephydatia muelleri*, and those in blue are other sponge sequences. Phylogeny generated in PhyML based on an alignment generated in PROMALS3D, alongside other genes of known homology downloaded from the nr database (with accession numbers as given on figure). Maximum likelihood probabilities are shown at each major node. The tree was rooted at midpoint. A full size version of this figure and further information is available for download from Ephybase (https://spaces.facsci.ualberta.ca/ephybase/), under the High Quality Figures link.

#### 9.2 Epithelia

Sponges have been often claimed to lack conventional epithelia, and yet in the *E. muelleri* genome we found the full gene complements for adherens junctions, tight junctions and focal adhesion machinery, excluding occludins (Supplementary Tables 19–20). A *claudin* gene is present, and while the epithelium of *Ephydatia muelleri* has been demonstrated to prevent the passage of small molecules [138], it is still to be tested whether *Emu claudin* has a role in that. We also found a number of genes for *type IV collagen* and *spongin short chain collagen* as well as *laminin*, *perlecan,* and *nidogen* involved in attachment to the basement membrane. Also present are genes involved in cell polarity (*PAR3/6, patj,* and *contactin*).

#### 9.3 Wnt pathway

Another example where we see evidence of gene duplication is within the Wnt signaling pathway. Wnt signaling, a metazoan innovation, defines the animal kingdom playing roles in spatial organization and the creation of cellular diversity (reviewed in[139]). In *E. muelleri*, Wnt signaling has been shown to play roles in body plan polarity and development of the aquiferous system [133, 140, 141 142] and Wnt ligands appear to be involved in polarity in marine sponges as well [143, 144, 145]. We find a near full complement of putative orthologues for the canonical and non-canonical Wnt pathways in the *E. muelleri* genome (Supplementary Figure 35, Supplementary Tables 19–20) with evidence of expansion for several pathway members.

While we only find two predicted Wnt ligands and two complete Frizzled receptors, we find extensive gene duplication of a group of Wnt antagonists, the Secreted frizzled related proteins (SFRPs). One scaffold (Em0016) contains six SFRP-like genes along with another gene possessing two cysteine-rich Wnt binding domains (CRD-FZ), but no putative Wnt binding sites. Of the six SFRP genes, three of them contain netrin domains and three do not. A second scaffold (Em0012) is home to Frizzled A, another SFRP gene, as well as a gene containing both the CRD-FZ domain and the membrane spanning domain characteristic of frizzled receptors, but with only 4 transmembrane domains instead of seven. Thus, the repertoire of SFRPs in *E. muelleri* is seven, each with distinct, but overlapping expression during *Ephydatia* development (Supplementary Figure 35).

The *Amphimedon queenslandica* genome appears to have four SFRPs (none with netrin domains) and the human genome has five. Another scaffold (Em0011) has two protein wntless (WLS) genes which in other metazoans play roles in regulating Wnt proteins. Thus, unlike the expansion of Wnt ligands seen in some sponges (e.g., [145, 146]) it appears that in *Ephydatia*, duplication of genes involved in modulation of Wnt signaling may play important roles. There are 15 other Wnt pathway genes that have putative duplications that occur on the same scaffold and numerous other genes with possible paralogs within the genome (Supplementary Table 20). For example, another scaffold (Em0012) contains two β-catenin genes, though one of the paralogs contains only four of the twelve armadillo repeats and three different scaffolds hold two paralogs of specific SMAD1 (Em08) and SMAD4-like (Em06, Em0010) genes.

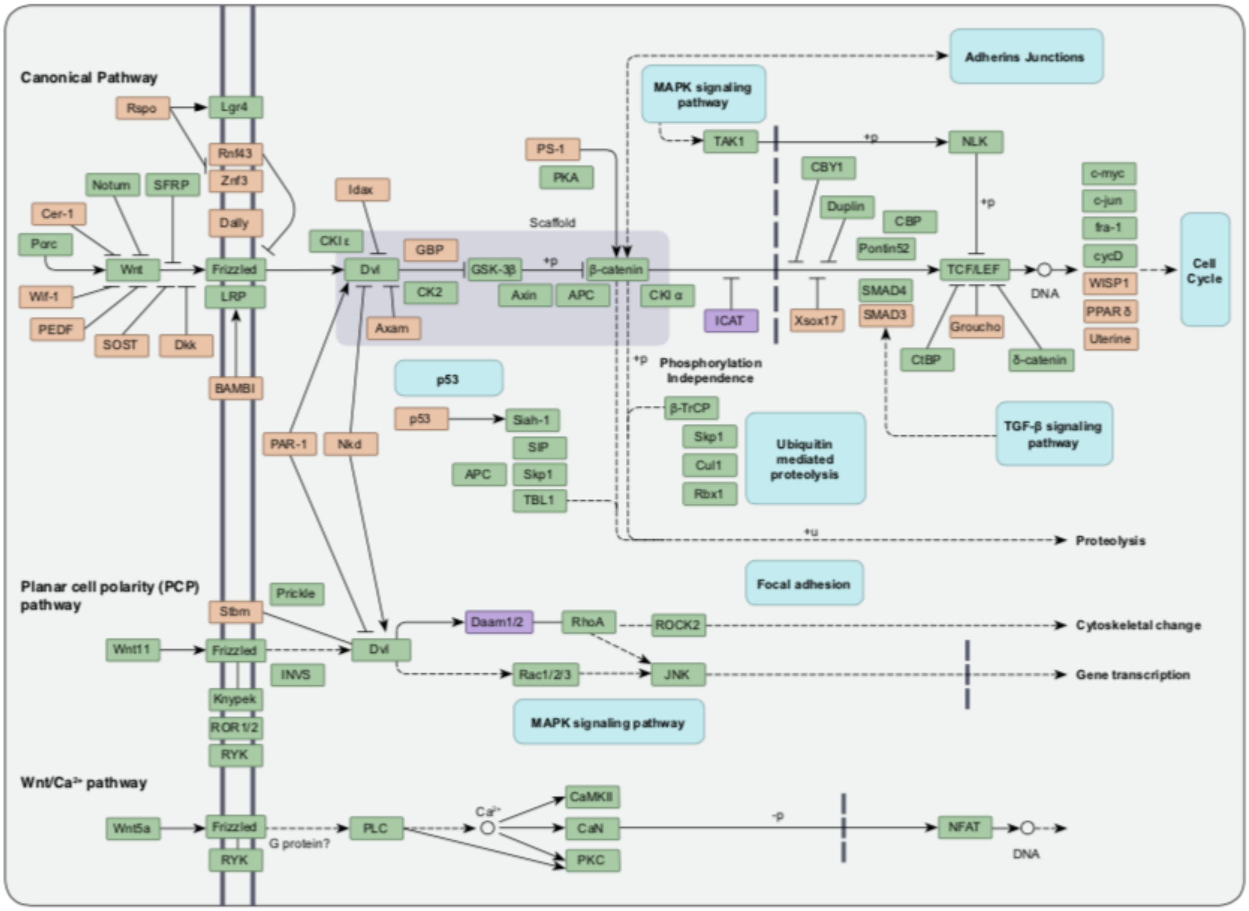

**Supplementary Figure 35:** Wnt signalling pathway gene presence, loss and likely ancestral absence in the *Ephydatia muelleri* gene complement, as mapped onto the canonical, planar cell polarity and Wnt/$Ca^{2+}$ gene pathways. Green = present. Purple = unclear absence, but presence of related gene. Orange = absent

#### 9.4 Comparative gene expression during development:

Several developmental pathways are conserved across metazoans, and they appear complete (or nearly complete) in sponges, cnidarians, ctenophores, and placozoans: the Wnt pathway, nuclear receptors, and the TGF-beta pathway [147]. Some others are restricted to metazoan lineages other than sponges, like the Hedgehog pathway in cnidarians and bilaterians, although several elements of the pathways are found in all sponge lineages [131]. Here we focused on the expression of Wnt, TGF-β and hedgehog pathways across *E. mulleri* development (see Supplementary Note 11 for details on stages) to understand how conserved these pathways are in early-splitting lineages of metazoans.

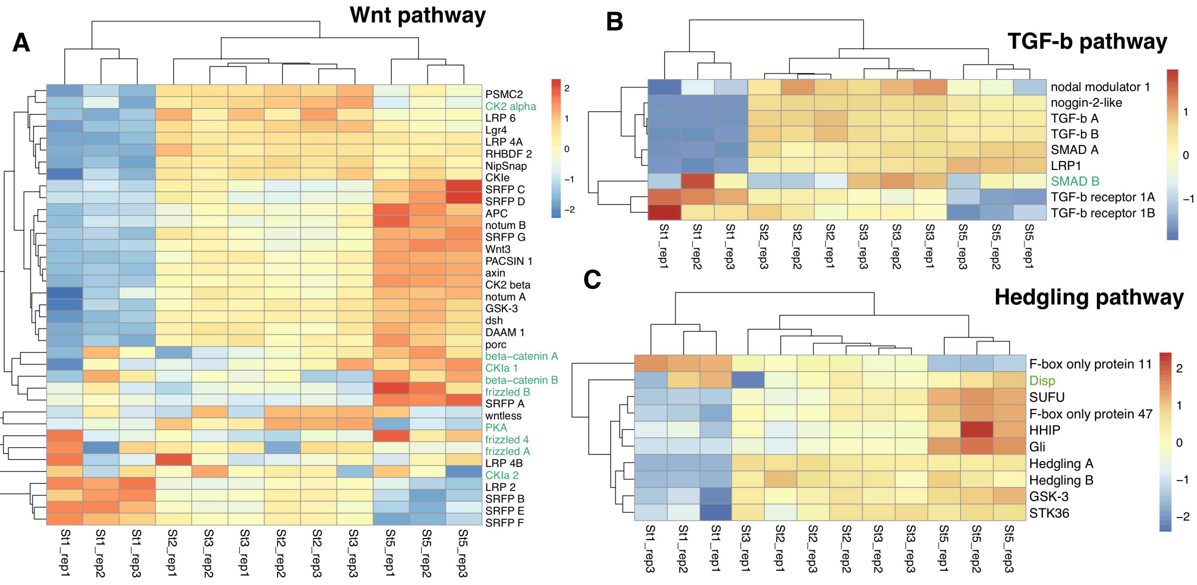

**Supplementary Figure 36:** Expression levels of A) Wnt pathway genes, B) TGF- β signalling related genes and C) Hedgling (hedgehog) pathway genes across the process of development. Expression shown on heat maps with relative expression levels as per scales shown at right of each figure. Names of genes shown in green indicate there is no differential expression between any time point for this gene.

*Wnt signalling pathway:*

Differential gene expression profiles indicate that three of the Secreted Frizzled Receptor Proteins (SFRP B, SFRP E, SFRP F) as well as two LRP receptors (LRP 2, LRP 4B) are upregulated during hatching from the stem cells stored in the gemmule (Stage 1), and subsequently downregulated with lowest expression levels at Stage 5 when the full sponge body plan is formed (see Supplementary Note 11 for more detail on stages). Interestingly, those three SFRPs are located in a tight cluster nested among the other SFRPs present on scaffold Em0016. The majority of genes in the canonical Wnt pathway, however, appear to be active at Stages 2-3, when choanocyte chambers, spicules, and the fundamental structures of the sponge begin to develop, and are most active at Stage 5 when the osculum is formed. This includes the other SFRP and LRP family members as well as the Wnt ligand (Wnt3). In past work we have found that Wnt signalling through inhibition of GSK3 is involved in formation of the osculum which develops at that stage [133, 140],while in other sponges, Wnt expression is associated with either ostia formation for example in *Oscarella lobularis* [143] or with polarization of the larva in *Amphimedon queenslandica* [115]. We also showed that SFRP G is expressed in a subpopulation of filopodia possessing amoeboid cells in the mesohyl of Stage

5 and that knockdown leads to ectopic oscula formation [142]. In *Nematostella vectensis*, SFRP is expressed on the aboral side during regeneration [148].

Frizzled receptors and beta-catenin orthologs are not differentially regulated across developmental stages at a significant level. In *Nematostella vectensis*, Wnt, Frizzled and beta-catenin are involved in determining the site of gastrulation [149], but generally the interaction of Wnt and beta-catenin in *Hydra* and *N. vectensis* is consistent with a role in cellular movements associated with invagination or evagination of epithelial sheets, whether gastrulation in *Nematostella* or budding as in *Hydra* [150]. In *Ephydatia* development from hatching through to osculum development, it is only when the osculum forms that an equivalent set of cellular movements is seen.

*The TGF- β signalling pathway:*

The TGF-β pathway regulates a variety of developmental processes in bilaterians, most notably dorsal-ventral axis specification, but also cell and tissue fate, immunity, and possibly germ layer identity [151,152]. In *Amphimedon queenslandica*, the expression of TGF-β in localized patterns that are distinct, but overlapping, with Wnt expression during embryogenesis suggests a role in axial polarity[144]. Gene expression data indicates that the *E. muelleri* TGF-β receptor (two are present) is upregulated in the hatching sponge (Stage 1) and less in the Stages 2 and 3, and expression is downregulated in Stage 5 once the sponge is fully formed. In contrast, expression of TGF-β ligands, agonists, and antagonists is significantly higher post hatching, when they are all relatively equally expressed across stages. Given that a key step to regulation of TGF- β signalling is the modulation of TGF-β receptor activity [153], it is notable that their upregulation occurs at the earliest developmental stage. Other genes potentially involved in TGF-β signalling are present including several tolloid-like proteins and keilin/chordin-like proteins.

*The Hedgehog Pathway:*

The Hedgehog (HH) pathway regulates cell and tissue identity during bilaterian development [154].In cnidarians, the expression patterns of the complete HH pathway are only known for *Nematostella vectensis*, where it is thought to indicate endodermal cell identity [155]. In sponges, there are no true *Hedgehog* genes [131, 144], but there are genes containing the Hedge domain that are able to interact with other signalling components. The expression of hedgling in the sponge *Amphimedon queenslandica* has only been studied in the larval stage, co-localized with the Wnt and TGF-β expression in the pigment ring [144]. In *E. muelleri*, expression of two orthologs of *Hedgling* is upregulated post-hatching, in Stage 2 through 5, and components of the HH signalling pathway including the Hedgehog Interacting Protein (HHIP), Suppressor of Fused (SUFU), and GLi like proteins are all significantly upregulated in Stage 5 sponges. It is difficult to draw any similarities between larval expression patterns and gemmule-hatching development, but it is clear that the HH pathway (minus Hedgehog) is playing a role during tissue organization in the development of the freshwater sponge *E. muelleri*.

Without spatial expression data it is not possible to draw inferences as to the specific role of these proteins in *E. muelleri* development, and which cells are involved in developmental patterning in this sponge. This is an open problem which represents an excellent starting point for future research.

**Supplementary Table 19:** Summary of gene content of *Ephydatia muelleri*. Full details in Supplementary Table 20 overleaf.

| **Process** | **Type** | **Genes found** |
|---|---|---|
| RNAi Machinery | | *Dicer, IFH1, DHX58, FAN CM, DDX58, PIWIL AGO1, TARBP2, DGCR8* |
| MBD Pathway | NuRD Complex core components | *CHD3/4/5, HDAC1/2, MBD2/3, MTA1/2/3, RBBP4/7,* |
| | Other key players | *GATAD2A/2B, DNMT1, DNMT3A* |
| Signalling | Hedgehog | *Patched, Smoothened, SUFU, HHIP, PKA, KIF7, Gli, CKI, GSK3, fused, Slimb/FBXW11, Dispatched, Hedgling* |
| | TGF-Beta | *TGF-β, TGF-β receptor, Noggin, Activin-receptor, LRP* |
| | Notch | *Notch/NOTC1, Delta, Jagged* |
| | Wnt | *See Table X for full complement* |
| Chemical signalling | Vesicle secretion/PSD | *β -catenin, CAMKII, citron, contactin, cript, GKAP, Ephrin-receptor, Homer, IP3R, PKC, PMCA, DLG, Erb-receptor, GRIP, MAGI, PICK1, Shank, SPAR, Lin7* |
| | Glutamate/GABA | *mGluR, GAD, EAAT, VGlut1, GABA, Tyrosine aminotransferase* |
| | Monamine neurotransmitter molecules | *AADC, Tyrosine hydroxylase, Qdpr, slcl8a2 (solute carrier organic anion transporter), PaH, Pnmt* |
| | Nitric oxide | *NOS, guanylate cyclase, cGMP-dependent protein kinase* |
| | Acetylcholine | *Ach (acelylcholinesterase precursor)* |
| Immune system | | *IRAK-4, Toll-receptor, MyD88, A2M, Nf_kb, MASP, IKK (inhibitor of nuclear factor kappa-B kinase), TAK1, TRAF (TNF receptor associated factor)* |
| Epithelia | | *Alpha-catenin, delta-catenin, vinculin, nidogen, perlecan, Collagen Type IV, short chain C4, Par3/6, Scribble, Stardust (MAGUK p55), Neurexin (potential ortholog), Claudin, Patj, contactin, laminin, Collagen XI, Paxillin, Talin, (FAK) Focal adhesion kinase, ILK (integrin-linked protein kinase)* |
| Reproduction | | *Vasa, PL10, nanos, Piwi, Mago-nashi, Tsunagi/RBM8A, Smaug/SAMD4A, maelstrom, Pumilio, Boule, FEM1, Attractin, mushashi, tudor* |
| Biomineralization | | *Silicatein, Aquaporin, Silintaphin, silicase, ArsB transporter, silicon cotransporter* |
| Ion channels | | *Voltage dependent Calcium channels, Kv channel, CNG-HCN, Leak, Ryanodine receptor, TRP* |

**Supplementary Table 20:** Full details of gene content in the genome of *Ephydatia muelleri*.

### Supplementary Table Gene Content

*Ephydatia muelleri* gene content detemined by blast. The top blast hits for each gene are shown with the probability (e-value). Green=present; Yellow=divergent ortholog; Red=absent from the genome

| Gene category | NCBI Gene name | E muelleri hits Gene names | e value | P/A |
|---|---|---|---|---|
| ***MDB Pathway*** | | | | |
| CHD3 | chromodomain helicase DNA binding protein 3 | Em0008g1027a | 0 | Green |
| CHD4 | chromodomain helicase DNA binding protein 4 | Em0008g1027a | 0 | Green |
| CHD5 | chromodomain helicase DNA binding protein 5 | Em0003g1398a | 0 | Green |
| HDAC1 | histone deacetylase 1 | Em0011g119a | -140 | Green |
| HDAC2 | histone deacetylase 2 | Em0011g119a | -137 | Green |
| MBD2 | methyl-CpG binding domain protein 2 | m0007g1367a | 0.21 | Yellow |
| MBD3 | methyl-CpG binding domain protein 3 | Em0015g1268a | 0.064 | Yellow |
| MTA1 | metastasis associated 1 | Em0019g569a, Em0019g560a | -105 | Green |
| RBBP4 | RB binding protein 4, chromatin remodeling factor | Em0023g375a | 0 | Green |
| GATAD2A | GATA zinc finger domain containing 2A | Em0009g427a | 0.007 | Yellow |
| GATAD2B | GATA zinc finger domain containing 2B | Em0012g312a | 0.025 | Yellow |
| DNMT1 | DNA methyltransferase 1 | Em0016g1020a, Em0020g6a | 0 | Green |
| DNMT3A | DNA methyltransferase 3 alpha | Em0006g900a | -89 | Green |
| ***RNAi Machinery*** | | | | |
| DICER1 | dicer 1, ribonuclease III | Em0019g657a | -114 | Green |
| IFIH1 | interferon induced with helicase C domain 1 | Em0004g961a, Em0006g501a, Em0039g31a | -56 | Green |
| DHX58 | DExH-box helicase 58 | Em0004g961a, Em0006g498a | -55 | Green |
| FANCM | FA complementation group M | Em0020g1101a | -179 | Green |
| DDX58 | ATP-dependent RNA helicase DDX58 | Em0006g498a, Em0006g501a | -71 | Green |
| PIWIL1 | piwi like RNA-mediated gene silencing 1 | Em0015g964a | 0 | Green |
| PIWIL2 | piwi like RNA-mediated gene silencing 2 | Em0017g775a | 0 | Green |
| PIWIL3 | piwi like RNA-mediated gene silencing 3 | Em0015g964a | 0 | Green |
| PIWIL4 | piwi like RNA-mediated gene silencing 4 | Em0015g964a | 0 | Green |
| AGO1 | argonaute RISC component 1 | Em0015g964a, Em0017g775a | -40 | Green |
| AGO2 | argonaute RISC component 2 | Em0017g775a, Em0015g964a | -40 | Green |
| TARBP2 | TARBP2 subunit of RISC loading complex | Em0023g405a | -7 | Yellow |
| DGCR8 | DGCR8 microprocessor complex subunit | Em0011g505a | -44 | Green |
| ***Cell signaling pathways*** | | | | |
| **Hedgehog Signaling** | | | | Red |
| Ptc | Patched 1 | Em0003g215a | -15 | Green |
| Smo | smoothened, frizzled class receptor | Em0012g384a | -30 | Green |
| SUFU | SUFU negative regulator of hedgehog signaling | Em0004g1352a | -87 | Green |
| HHIP | hedgehog interacting protein | Em007g755a, Em007g757a, Em0023g526a | 0 | Green |
| PKA | protein kinase cAMP-activated catalytic subunit | Em0020g922a | -129 | Green |
| cos2/ KIF7 | Kinesin-like protein KIF7 | Em0008g287a | -83 | Green |
| Gli | GLI family zinc finger 1 | Em0007g214a | -75 | Green |
| CKI | casein kinase 1 epsilon | Em0020g644a | -162 | Green |
| GSK3 | Glycogen synthase kinase-3 alpha | Em0006g1044a | -154 | Green |
| fused | Serine/threonine-protein kinase 36 (fused homo | Em0018g1081a, Em0015g1158a | -42 | Green |
| Slimb/ FBXW11 | F-box and WD repeat domain containing 11 | Em0013g821a | -178 | Green |
| Disp | dispatched RND transporter family member 1 | Em0008g1258a, Em0657g4a | -15 | Green |
| Hedgeling | Amphimedon hedgling | Em0007g755a | 0 | Green |
| Dsh | Double-stranded RNA-specific adenosine deam | Em0037g9a, Em0017g314a | -88 | Green |
| **TGF-beta** | | | | Red |
| TGF-b | transforming growth factor beta 1 | Em0015g1213a, Em0015g1236a, Em0019g98a | -28 | Green |
| TGF-b receptor | TGF-beta receptor type-1 | Em0012g37a | -103 | Green |
| Noggin/ Nog | Noggin | Em0014g630a | -12 | Green |
| Gro/ CXCL1 | C-X-C motif chemokine ligand 1 | Em0021g962a | 3.7 | Red |
| Activin-receptor | Activin receptor type-1 | Em0012g36a, Em0012g9a | -105 | Green |
| LRP | Low density lipoprotein | Em0011g684a, Em0010g765a, Em0002g1023a | -20 | Green |
| **Notch/Delta** | | | | Red |
| Notch/ NOTC1 | Neurogenic locus notch homolog | Em0115g6a | 0 | Green |
| Delta | Delta | Em0018g256a, Em0115g5a | -66 | Green |
| Jagged | Protein jagged-1 | Em0011g1192a | -137 | Green |
| **Wnt** | | | | |
| Wnt | Proto-oncogene Wnt-1 | Em0008g1080a | -43 | Green |
| 0 | Proto-oncogene Wnt-3 | Em0012g1035a | -41 | Green |
| WIF | WNT inhibitory factor 1 | Em0115g5a, Em0018g257a | -21 | Red |
| Porc | Protein-serine O-palmitoleoyltransferase porcup | Em0016g421a | -64 | Green |
| Notum | palmitoleoyl-protein carboxylesterase | Em0021g945a, Em0009g1239a | -25 | Green |
| CKIe | casein kinase 1 epsilon | Em0020g644a | -162 | Green |
| CKIa | casein kinase 1 alpha 1 like | Em0013g329a | -148 | Green |
| CK2 | casein kinase 2 alpha 1 | Em0018g165a, Em0016g68a, Em0016g82a | -158 | Green |
| B-catenin | Catenin beta-1 | Em0012g557a, Em0012g536a | -177 | Green |
| GSK3B | glycogen synthase kinase 3 beta | Em0006g1044a | -171 | Green |

| Gene category | NCBI Gene name | E muelleri hits Gene names | e value | P/A |
|---|---|---|---|---|
| GSK3-beta intera | GSK3-beta interaction protein | Em0004g929a | 0 | 🟢 |
| APC | APC regulator of WNT signaling pathway | Em0010g855a | -66 | 🟢 |
| Axin | axin 1 | Em0020g197a | -7 | 🟢 |
| SFRP | Secreted frizzled-related protein | Em0016g747a, Em0016g751a, Em0016g746a, Em0016g750a, Em0016g745a | -13 | 🟢 |
| 0 | Secreted frizzled-related protein 3 | Em0012g238a | -15 | |
| Frizzled | frizzled receptor/ Secreted frizzled-related protei | Em0012g384a, Em0023g388a, Em0012g14a | -15 | 🟢 |
| Dkk | dickkopf WNT signaling pathway inhibitor 1 | Em0010g848a | 0.62 | 🔴 |
| CBP | poly(rc)-binding protein | Em0007g260a, Em0022g450a | -60 | 🟢 |
| SMAD1 | Mothers against decapentaplegic homolog 1 | Em0022g765a, Em0810g2a, Em0810g3a | -148 | 🟢 |
| SMAD4 | Mothers against decapentaplegic homolog 4 | Em0010g75a, Em0010g45a, Em0648g4a | -106 | 🟢 |
| WLS | Wnt ligand secretion mediator | Em0011g389a, Em0011g6a | -38 | 🟢 |
| Cer-1 | cerberus 1, DAN family BMP antagonist | no hits | 0 | 🔴 |
| PEDF | serpin family F member 1 | no hits | 0 | 🔴 |
| SOST | sclerostin | no hits | 0 | 🔴 |
| RSPO1 | R-spondin 1 | Em0009g1128a, Em0009g1139a | -7 | 🔴 |
| LGR4 | leucine rich repeat containing G protein-coupled | Em0003g1326a | -32 | 🟢 |
| RNF43 | ring finger protein 43 | no sig hits | 0 | 🔴 |
| ZNRF3 | zinc and ring finger 3 | no sig hits | 0 | 🔴 |
| LRP5/6 | LDL receptor related protein 6 | Em0010g765a, Em0002g1023a, Em0003g1221a | -98 | 🟢 |
| BAMBI | BMP and activin membrane bound inhibitor | no sig hits | 0 | 🔴 |
| Idax | Dvl-binding protein IDAX | no sig hits | 0 | 🔴 |
| DVL1 | dishevelled segment polarity protein 1 | Em0008g347a, Em0019g830a | -73 | 🟢 |
| AXAM2 | sentrin-specific protease 2 (axin associating mo | Em0076g2a, Em0011g1018a, Em0768g1a | -22 | 🟢 |
| GBP (FRAT1) | FRAT regulator of WNT signaling pathway 1 | no sig hits | 0 | 🔴 |
| PS-1 | presenilin 1 | no sig hit | 0 | 🔴 |
| PKA | protein kinase cAMP-activated catalytic subunit | Em0011g81a | -155 | 🟢 |
| RYK | receptor like tyrosine kinase | Em0021g39a | -57 | 🟢 |
| ROR1/2 | receptor tyrosine kinase like orphan receptor 1 | Em0018g1165a | -83 | 🟢 |
| Knypek | K-glypican, glypican 4 | Em0021g142a | -12 | 🟢 |
| Stbm | VANGL planar cell polarity protein 2 | no sig hits | 0 | 🔴 |
| Nkd | NKD inhibitor of WNT signaling pathway 1 | no sig hits | 0 | 🔴 |
| p53 | tumor protein p53 | 0 | 0 | 🔴 |
| Siah-1 | siah E3 ubiquitin protein ligase 1 | Em0003g1558a | -112 | 🟢 |
| SIP | calcyclin binding protein | Em0016g544a | -31 | 🟢 |
| Skp1 | S-phase kinase associated protein 1 | Em0007g982a | -58 | 🟢 |
| TBL1 | transducin beta like 1 X-linked | Em1340g2a | -32 | 🟢 |
| Prickle | prickle planar cell polarity protein 1 | Em0023g581a | -58 | 🟢 |
| INVS | inversin | Em0012g1094a, Em0021g744a, Em0021g733a | -120 | 🟢 |
| Daam1/2 | dishevelled associated activator of morphogene | Em0022g422a, Em0020g318a, Em0015g844a | -36 | 🟡 |
| ROCK2 | Rho-associated protein kinase 2 | Em0015g877a | -98 | 🟢 |
| JNK | mitogen-activated protein kinase 8 | Em0002g1177a | -116 | 🟢 |
| PLC | phospholipase C beta 1 | Em0006g969a | -133 | 🟢 |
| CaMKII | calcium/calmodulin dependent protein kinase II | Em0012g965a, Em0001g1211a, Em0001g1272a | -135 | 🟢 |
| CaN | protein phosphatase 3 catalytic subunit alpha | Em0002g1605a | 0 | 🟢 |
| PKC | protein kinase C alpha | Em0020g1088a | 0 | 🟢 |
| NFAT | nuclear factor of activated T cells 1 | Em0022g704a | 0 | 🟢 |
| RhoA | Ras homolog gene family, member A | Em0004g1702a plus many | 0 | 🟢 |
| RAC1/2/3 | Rac family small GTPase 1, Rac family small G | Em0007g856a plus many | 0 | 🟢 |
| **MAPK signaling from Canonical Wnt** | | | | |
| CKIa | casein kinase 1 alpha 1 like | Em0013g329a | -148 | 🟢 |
| B-TrCP | F-box and WD repeat domain containing 11 | Em0013g821a | -178 | 🟢 |
| Skp1 | S-phase kinase associated protein 1 | Em0007g982a | -58 | 🟢 |
| Cul1 | cullin 1 | Em0007g577a | 0 | 🟢 |
| Rbx1 | ring-box 1 | Em0022g219a | -54 | 🟢 |
| TAK1 | mitogen-activated protein kinase kinase kinase | Em0023g663a | -44 | 🟢 |
| ICAT | catenin beta interacting protein 1 | Em0008g1271a | -7 | 🟡 |
| NLK | nemo like kinase | Em0015g1255a | -122 | 🟢 |
| CBY1 | chibby family member 1, beta catenin antagonis | Em0020g453a | -9 | 🟢 |
| Duplin | chromodomain helicase DNA binding protein 8 | Em0018g1127a | -151 | 🟢 |
| Xsox17 | SRY-box transcription factor 17 | no sig hits | 0 | 🔴 |
| Pontin52 | RuvB like AAA ATPase 1 | Em0011g786a | 0 | 🟢 |
| SMAD4 | Mothers against decapentaplegic homolog 4 | Em0010g75a | -106 | 🟢 |
| SMAD3 | SMAD family member 3 | Em0022g765a | 0 | 🔴 |
| CBP | poly(rc)-binding protein | Em0007g260a | -60 | 🟢 |
| TCF/LEF | lymphoid enhancer binding factor 1 | Em0002g42a | -34 | 🟢 |
| CtBP | C-terminal binding protein 1 | Em0013g121a | -54 | 🟢 |
| d-Catenin | catenin delta 2 | Em0017g517a | -38 | 🟢 |
| c-myc | MYC proto-oncogene, bHLH transcription factor | Em0023g45a | -19 | 🟢 |
| c-jun | Jun proto-oncogene, AP-1 transcription factor su | Em0011g1167a | -17 | 🟢 |
| fra-1 | FOS like 1, AP-1 transcription factor subunit | Em0019g326a | 0 | 🟢 |
| cycD | cyclin D2 | Em0021g487a | -15 | 🟢 |
| WISP1 | WNT1-inducible-signaling pathway protein 1 | Em0012g778a | -5 | 🔴 |
| PPARg | peroxisome proliferator activated receptor delta | Em0001g69a | -29 | 🔴 |
| Uterine | matrix metallopeptidase 7 | no sig hits | 0 | 🔴 |

| Gene category | NCBI Gene name | E muelleri hits Gene names | e value | P/A |
|---|---|---|---|---|
| ***Chemical signaling*** | | | | |
| **Vesicle secretion/PSD type genes** | | | | |
| B-catenin | Catenin beta-1 | Em0012g557a | -177 | 🟢 |
| CAMKII | calcium calmodulin protein Kinase II | Em0012g965a | -167 | 🟢 |
| citron | Citron Rho-interacting kinase | Em0015g877a | -107 | 🟢 |
| cortactin | Src substrate cortactin | Em0021g307a | -109 | 🟢 |
| cript | Cysteine-rich PDZ-binding protein | Em0019g314a | -41 | 🟢 |
| GKAP | G kinase-anchoring protein 1 | Em0009g162a, Em0009g138a | -20 | 🟢 |
| Ephrin-receptor | Ephrin receptor | Em0008g185a, Em0008g186a | -93 | 🟢 |
| Homer | Homer protein homolog 1 | Em0009g974a | -46 | 🟢 |
| IP3R | Inositol 1,4,5-trisphosphate receptor type 1 | Em0014g49a | -95 | 🟢 |
| PKC | Protein kinase C alpha type | Em0020g1088a, Em0019g897a, Em0019g897a | 0 | 🟢 |
| PMCA | Plasma membrane calcium-transporting ATPas | Em0014g439a | 0 | 🟢 |
| DLG | Disks large homolog 1 | Em0021g795a, Em0021g788a | -163 | 🟢 |
| Erb-receptor | Receptor tyrosine-protein kinase erbB-2 | Em0012g156a, Em0001g1133a | -78 | 🟢 |
| GRIP | Glutamate receptor-interacting protein 1 | Em0021g795a, Em0021g788a | -17 | 🟢 |
| MAGI | Membrane-associated guanylate kinase, WW a | Em0001g3274a, Em0001g3266a | -27 | 🟢 |
| PICK1 | PRKCA-binding protein | Em0020g828a | -45 | 🟢 |
| Shank | SH3 and multiple ankyrin repeat domains protei | Em0021g394a | -63 | 🟢 |
| SPAR | Small regulatory polypeptide of amino acid resp | Em0006g871a | 0.35 | 🔴 |
| Lin7 | Protein lin-7 homolog A | Em0003g1413a | -51 | 🟢 |
| **Glutamate** | | | | |
| Gl5 | glutaminase [Homo sapiens] | Em0019g204a | -12 | 🔴 |
| Glud1 | glutamate dehydrogenase [Homo sapiens] | none | 0 | 🔴 |
| mGluR | metabotropic glutamate receptor 1 isoform alph | Em0026g49a, Em0026g54a | -128 | 🟢 |
| | | Em0468g1a, Em0312g2a, Em0022g766a | -118 | 🟢 |
| | | Em0003g578a, Em0003g599a, Em0003g477a | -54 | 🟢 |
| | | Em0024g62a, Em0024g79a | -116 | 🟢 |
| iGluR | Glutamate receptor ionotropic, kainate 1 | Em0024g62a, Em0006g1337a | -4 | 🔴 |
| GAD | Glutamate decarboxylase GAD1 protein [Homo | Em0001g1154a | -65 | 🟢 |
| EAAT | glutamate transporter [Mus musculus] | Em0011g1007a | -16 | 🟢 |
| VGlut1 | Brain-specific Na(+)-dependent inorganic phosp | Em0074g13a | -89 | 🟢 |
| GABA | | | | |
| GABA | Gamma-aminobutyric acid type B receptor subu | Em0016g376a | -97 | 🟢 |
| | | Em0016g375a | | |
| | | Em0020g696a | | 🟢 |
| | | Em0020g695a | | |
| | | Em0013g421a | | 🟢 |
| | | Em0013g917a | | |
| | | Em0013g425a | | |
| | | Em0816g3a | | 🟢 |
| | | Em0006g1271a | | 🟢 |
| | | Em0006g1258a | | |
| ABAT | 4-aminobutyrate aminotransferase, mitochondri | >Em0013g271a | -5 | 🔴 |
| TAT | tyrosine aminotransferase [Homo sapiens] | Em0018g57a | -146 | 🟢 |
| **Monamine neurotransmitter molecules** | | | | |
| 5HT receptor | 5-hydroxytryptamine receptor 5B | Em0010g751a | -10 | 🔴 |
| TpH | tryptophan 5-hydroxylase 1 [Homo sapiens] | Em0021g279a | -145 | 🔴 |
| AADC | aromatic-L-amino-acid decarboxylase isoform 4 | Em0001g1154a, Em0001g1241a | -37 | 🟢 |
| 0 | dopa decarboxylase, isoform D [Drosophila mel | Em0001g1241a | -28 | 🟢 |
| Th | tyrosine hydroxylase [Mus musculus] | >Em0021g279a | -134 | 🔴 |
| Qdpr | Quinoid dihydropteridine reductase [Mus muscu | >Em0013g272a | -85 | 🟢 |
| slc18a2 | solute carrier organic anion transporter family m | Em0005g710a, Em0094g15a, Em0005g832a | -75 | 🟢 |
| PaH | phenylalanine hydroxylase [Homo sapiens] | Em0021g279a, Em0009g692a | -167 | 🟢 |
| Dbh | dopamine-beta-hydroxylase [Mus musculus] | >Em0006g282a | -77 | 🟢 |
| Pnmt | phenylethanolamine-N-methyltransferase [Mus | Em0014g635a | -15 | 🟡 |
| Dopamine rece | beta-2 adrenergic receptor [Mus musculus] | Em0010g751a | -9 | 🔴 |
| | | | 0 | |
| | | | 0 | |
| **Nitric Oxide** | | | 0 | |
| NOS | nitric oxide synthase [Mus musculus domesticus | Em0015g78a, Em0162g6a | 0 | 🟢 |
| sGC | guanylate cyclase [Mus musculus] | Em0011g518a | -149 | 🟢 |
| PKG-1 | cGMP-dependent protein kinase 1 isoform 1 [H | >Em0010g719a | 0 | 🟢 |
| **Acetylcholinesterase** | | | | |
| Ach | acetylcholinesterase precursor [Bos taurus] | Em0021g924a | -36 | 🟢 |
| ***Immune Genes*** | | | | |
| IRAK-4 | Interleukin-1 receptor-associated kinase 4 | Em0009g49a | -46 | 🟢 |
| Toll-receptor-lik | Toll-like receptor 2 (TLR2) | Em0012g672a, Em0012g665a | -11 | 🟡 |
| MyD88 | Myeloid differentiation primary response protein | Em0024g416a | -22 | 🟢 |
| A2M | Alpha-2-macroglobulin | Em0003g56a | 0.98 | 🔴 |

| Gene category | NCBI Gene name | E muelleri hits Gene names | e value | P/A |
|---|---|---|---|---|
| Nf_kb | Nuclear factor NF-kappa-B p105 subunit | Em0006g1255a, Em0249g3a | -109 | 🟢 |
| MASP | Mannan-binding lectin serine protease 1 | Em0023g351a | -23 | 🟢 |
| IKK | Inhibitor of nuclear factor kappa-B kinase subun | Em0010g935a, Em0019g38a, Em0006g1249a, Em0011g81a, Em0010g723a | -22 | 🟢 |
| | Inhibitor of nuclear factor kappa-B kinase subun | Em0015g304a, Em0015g296a | -26 | 🟢 |
| | Inhibitor of nuclear factor kappa-B kinase subun | Em0001g894a | -37 | 🟢 |
| TAK1 | Nuclear receptor subfamily 2 group C member 2 | Em0001g69a, Em0001g70a, Em0001g73a, Em0001g71a | -25 | 🟢 |
| TRAF | TNF receptor-associated factor 1 | Em0004g908a | -51 | 🟢 |
| | TNF receptor-associated factor 2 | Em0016g278a | -77 | 🟢 |
| | TNF receptor-associated factor 3 | Em0008g709a | -98 | 🟢 |
| | TNF receptor-associated factor 4 | Em0008g751a | -44 | 🟢 |
| | E3 ubiquitin-protein ligase TRAF7 | Em0020g1004a | -177 | 🟢 |
| ***Epithelial genes*** | | | | |
| alpha-catenin | Catenin alpha-1 | Em0013g145a | -95 | 🟢 |
| delta-catenin | Catenin delta-1 | Em0017g517a | -31 | 🟢 |
| Vinculin | Vinculin | Em0017g182a | -61 | 🟢 |
| nidogen | Nidogen-1 | Em0010g765a | -38 | 🟢 |
| 0 | Nidogen-2 | Em0023g304a, Em0001g322a, Em0023g300a | -34 | 🟢 |
| | Basement membrane-specific heparan sulfate p | Em0011g1158a | -95 | 🟢 |
| Collagen-Type-I | Collagen type IV hydra vulgaris | Em0006g916a | -98 | 🟢 |
| SSCC | Spongin short chain collagen (Efluviatilis) | Em0001g1342a, Em0001g1339a | 0 | 🟢 |
| Par6 | Partitioning defective 6 homolog alpha | Em0018g717a | -65 | 🟢 |
| Scribble (SCRIE | Protein scribble homolog | Em0005g1163a | -104 | 🟢 |
| Stardust | MAGUK p55 subfamily member 7 (MPP7) | Em0012g1042a | -66 | 🟢 |
| Crumbs | Protein crumbs homolog 1 | Em0011g1192a | -101 | 🟢 |
| Neurexin | Neurexin-1 | Em0022g301a | -23 | 🔴 |
| Claudin | Claudin | Em0008g1042a | -6 | 🟢 |
| Patj | InaD-like protein/ Pals1-associated tight junctior | Em0011g11a | -45 | 🟢 |
| Contactin | Contactin-1 | Em0006g1150a | -39 | 🟢 |
| Laminin | Laminin subunit alpha-1 | Em0016g522a, Em0016g530a, Em0019g102a | -98 | 🟢 |
| Collagen-XI | Collagen alpha-1(XI) chain | Em0075g5a, Em0007g546a, Em0007g528a, Em0007g559a | -23 | 🟢 |
| 0 | Collagen alpha-2(XI) chain | Em0006g932a | -32 | 🟢 |
| Paxillin | Paxillin (PXN) | Em0015g714a | -113 | 🟢 |
| Talin | Talin-1 | Em0010g62a | 0 | 🟢 |
| 0 | Talin-2 | Em0010g50a | 0 | 🟢 |
| alpha-actinin | Alpha-actinin 1 | Em0019g756a, Em0014g868a | -125 | 🟢 |
| 0 | Alpha-actinin 4 | Em0019g756a | -122 | 🟢 |
| FAK | Focal adhesion kinase 1 | Em0008g879a, Em0010g799a, Em0021g626a | -59 | 🟢 |
| ILK | Integrin-linked protein kinase | Em0022g205a | -49 | 🟢 |
| ***Reproductive genes*** | | | | |
| vasa | vasa isoform A | Em0011g60a, Em0021g68a, Em0021g63a, Em0009g187a | 4e-162, | 🟢 |
| PL10 | PL10 protein [Ephydatia muelleri] | Em0021g68a, Em0021g63a | 3e-133, | 🟢 |
| nanos | nanos | Em0013g491a | 9e-47, | 🟢 |
| Piwi | Piwi | Em0015g964a | 0.0, | 🟢 |
| Piwi | Piwi | Em0017g775a | 9e-158, | 🟢 |
| mago-nashi | Protein mago nashi homolog | Em0010g175a | -32 | 🟢 |
| Tsunagi/ RBM8 | RNA-binding protein 8A | Em0009g1156a | -37 | 🟢 |
| Smaug/ SAMD4 | Protein Smaug homolog 1 | Em0019g306a | -26 | 🟢 |
| maelstrom | maelstrom | Em0006g1227a, Em0006g1236a | 3e-133, | 🟢 |
| Germ-cell-less | Germ cell-less protein-like 1 | Em0011g247a | -11 | 🟡 |
| Pumilio | Pumilio homolog 1 | Em0005g1324a, Em0011g28a | -78 | 🟢 |
| Boule | Protein boule-like (BOLL) | Em0004g1595a | -20 | 🟢 |
| bruno | bruno 1, isoform G [Drosophila melanogaster] | NONE | 0 | 🔴 |
| FEM1 | Protein fem-1 homolog A | Em0005g1212a, Em0018g302a | -34 | 🟢 |
| DMRT1 | Doublesex- and mab-3-related transcription fact | Em0006g916a | 5 | 🔴 |
| Vitellogenin | Microsomal triglyceride transfer protein large sul | Em0092g12a | 0.018 | 🔴 |
| Attractin | Attractin | Em0013g44a | -48 | 🟢 |
| musashi | RNA-binding protein Musashi homolog 2 isoforn | Em0015g1117a | 2e-11, | 🟢 |
| tudor | tudor [Tribolium castaneum] | Em0151g7a | 1e-29, | 🟢 |
| | | Em0023g207a | 3e-29, | |
| | | Em0011g41a | 3e-24, | |
| | | Em0010g41a | 6e-18, | |
| | | Em0013g844a | 5e-14, | |
| | | Em1196g1a | 1e-13, | |
| | | Em1275g1a | 2e-13, | |
| | | Em0013g845a | 2e-13, | |
| | | Em0531g2a | 8e-13, | |
| | | Em0016g1178a | 1e-11, | |
| R-spondin 1 | R-Spondin 1 mouse | NONE | 0 | 🔴 |
| Sox9 | SOX9 [Homo sapiens] | Em0023g195a | 2e-34, | 🔴 |
| Sox9 | SOX9 [Homo sapiens] | Em0021g940a | 5e-27, | 🔴 |
| Sox8 | SRY (sex determining region Y)-box 8, isoform ( | Em0023g195a | 6e-10, | 🔴 |
| Sox8 | SRY (sex determining region Y)-box 8, isoform ( | Em0021g940a | 2e-08, | 🔴 |
| Sox8 | SRY (sex determining region Y)-box 8, isoform ( | Em0023g270a | 3e-06, | 🔴 |

| Gene category | NCBI Gene name | E muelleri hits Gene names | evalue | P/A |
|---|---|---|---|---|
| ***Biomineralization*** | | | | |
| Silicatein | Silicatein (Petrosia ficiformis) | Em0017g930a | 2e-129, | 🟢 |
| Silicatein | Silicatein (Petrosia ficiformis) | Em0011g351a | 4e-116, | 🟢 |
| Silicatein | Silicatein (Petrosia ficiformis) | Em0056g25a | 3e-115, | 🟢 |
| Silicatein | Silicatein (Petrosia ficiformis) | Em0003g1460a | 3e-115, | 🟢 |
| Silicatein | Silicatein (Petrosia ficiformis) | Em0011g357a | 4e-115, | 🟢 |
| Silicatein | Silicatein (Petrosia ficiformis) | Em0011g353a | 3e-114, | 🟢 |
| Aquaporin 9 | Aquaporin (Ephydatia fluviatilis) | Em0014g564a | 0.0, | 🟢 |
| Aquaporin 9 | Aquaporin (Ephydatia fluviatilis) | Em0014g563a | 0.0, | 🟢 |
| Aquaporin 9 | Aquaporin (Ephydatia fluviatilis) | Em0014g565a | 2e-131, | 🟢 |
| Aquaporin 9 | Aquaporin (Ephydatia fluviatilis) | Em0614g1a | 1e-87, | 🟢 |
| Aquaporin 9 | Aquaporin (Ephydatia fluviatilis) | Em0019g920a | 5e-28, | 🟢 |
| Aquaporin 9 | Aquaporin (Ephydatia fluviatilis) | Em0246g8a | 3e-27, | 🟢 |
| Aquaporin 9 | Aquaporin (Ephydatia fluviatilis) | Em0007g1442a | 5e-26, | 🟢 |
| Aquaporin 9 | Aquaporin (Ephydatia fluviatilis) | Em0019g930a | 6e-22, | 🟢 |
| Aquaporin 9 | Aquaporin (Ephydatia fluviatilis) | Em0019g928a | 1e-21, | 🟢 |
| Aquaporin 9 | Aquaporin (Ephydatia fluviatilis) | Em0246g9a | 2e-21, | 🟢 |
| Silintaphin 1 | silintaphin-1 [Suberites domuncula] | Em0022g61a | 2e-27, | 🟢 |
| Silicase | CA_SubDo [Suberites domuncula] (Carbonic an | Em0014g599a | 2e-47, | 🟢 |
| Silicase | CA_SubDo [Suberites domuncula] (Carbonic an | Em0014g559a | 2e-45, | 🟢 |
| Silicase | CA_SubDo [Suberites domuncula] (Carbonic an | Em0014g586a | 1e-38, | 🟢 |
| Silicase | CA_SubDo [Suberites domuncula] (Carbonic an | Em0557g4a | 3e-35, | 🟢 |
| Silicase | CA_SubDo [Suberites domuncula] (Carbonic an | Em0016g908a | 3e-35, | 🟢 |
| Silicase | CA_SubDo [Suberites domuncula] (Carbonic an | Em0008g787a | 3e-35, | 🟢 |
| Silicase | CA_SubDo [Suberites domuncula] (Carbonic an | Em0436g4a | 3e-35, | 🟢 |
| Silicase | CA_SubDo [Suberites domuncula] (Carbonic an | Em0436g3a | 3e-35, | 🟢 |
| Silicase | CA_SubDo [Suberites domuncula] (Carbonic an | Em0003g1270a | 3e-35, | 🟢 |
| Silicase | CA_SubDo [Suberites domuncula] (Carbonic an | Em0003g1269a | 3e-35, | 🟢 |
| ArsB transporte | PREDICTED: putative transporter arsB [Amphin | Em0023g939a | 0.0, | 🟢 |
| ArsB transporte | PREDICTED: putative transporter arsB [Amphin | Em0007g624a | 1e-170, | 🟢 |
| ArsB transporte | PREDICTED: putative transporter arsB [Amphin | Em0007g622a | 9e-153, | 🟢 |
| ArsB transporte | PREDICTED: putative transporter arsB [Amphin | Em0007g629a | 3e-76, | 🟢 |
| Silicon cotransp | natriumbicarbonate_silicic_acid_cotransporter [S | Em0012g1050a | 6e-100, | 🟢 |
| Silicon cotransp | natriumbicarbonate_silicic_acid_cotransporter [S | Em0086g4a | 2e-77, | 🟢 |
| Silicon cotransp | natriumbicarbonate_silicic_acid_cotransporter [S | Em0023g877a | 5e-68, | 🟢 |
| 0 | | | | |
| ***Ion channels*** | | | | |
| Calcium | voltage-dependent L-type calcium channel subu | Em0001g1153a, Em0001g1240a | -234 | 🟢 |
| | | Em0001g1152a | -66 | 🟢 |
| | | Em0016g930a, Em0015g615a | -36 | 🟢 |
| | | Em0015g616a, Em0015g619a | -28 | 🟢 |
| | | Em0006g661a, Em0015g618a | -14 | 🟢 |
| Sodium | sodium channel, voltage-gated | Em0001g1240a | -150 | 🔴 |
| Potassium | Potassium - v-gated | Em0001g1240a | -18 | 🔴 |
| | | Em0019g967a | -18 | 🟢 |
| | | Em0001g1153a | -15 | |
| | | Em0005g1627a | -10 | |
| Enac | epithelial sodium activated channel | NONE | | 🔴 |
| CNG-HCN | cyclic nucleotide gated-hyperpolarization activat | Em0022g408a, Em0022g420a, Em0022g419a, Em0023g424a | 0 | 🟢 |
| CIC | Chloride channels | Em0008g534a | 0 | 🟢 |
| | | Em0022g736a | -205 | |
| | | Em0008g535a | -166 | |
| | | Em0008g497a | -165 | |
| GIC | Glutamate gated Ion channels | NONE | | 🔴 |
| Leak | Leak channels | Em0001g1240a | -148 | 🔴 |
| RyR | Ryanodine Receptor | Em0014g49a, Em0015g80a, Em1035g1a | 0 | |
| TRP | Transient receptor potential | Em0023g413a | -62 | 🟢 |
| | | Em0023g412a | -57 | 🟢 |
| | | Em0001g1088a | -51 | 🟢 |
| | | Em0023g414a | -51 | |
| | | Em0011g571a | -51 | |
| | | Em0023g455a | -37 | |
| | | Em0023g456a | -34 | |
| | | Em0017g848a | -33 | |

### Supplementary Note 10: Amplicon analysis of holobiont content

Sponges are considered holobionts, hosting a huge diversity of microbes within their bodies [156]. The microbiome of freshwater sponges, although much less known than that of marine sponges, has recently been assessed as even more diverse than that of their marine counterparts [157].

#### 10.1 Microbial community structure methods

To analyse the microbial community composition of *Ephydatia muelleri* across more than 6,800 km, we collected gemmules and tissue containing gemmules from adult sponges in 6 locations in the northern hemisphere (Supplementary Table 21, Supplementary Figure 37A). Both unhatched and hatched gemmules were analysed, as well as adult tissues containing gemmules. Gemmules from the adult tissue were removed as much as possible before DNA extraction. When gemmules were hatched, hatching was performed for 1 week following [19]. Each sample was amplified and sequenced in duplicate (pseudoreplicates *a* and *b*). For adult tissue this implied two different Supplementary Notes of the tissue, and for gemmules, each pseudoreplicate included 5 gemmules which were processed separately.

**Supplementary Table 21.** Details of sample collection.

| Location | Coordinates | Stage | Phase | N |
|---|---|---|---|---|
| Montgomery Canal, Oswestry, UK | 52.782, -3.089 | adult | adult | 3 |
| Youngs Pond, Virginia, USA | 37.598, -77.468 | gemmule | hatched | 1 |
| Sooke Reservoir, British Columbia, Canada | 48.50952, -123.691286 | gemmule | unhatched | 2 |
| | | | hatched | 1 |
| Androscoggin River, Maine, USA | 44.0141, -70.0579 | gemmule | hatched | 1 |
| Twitchell Brook, Maine, USA | 44.214, -70.490 | gemmule | hatched | 1 |
| O'Connor Lake, British Columbia, Canada | 50.541161, -127.250219 | gemmule | unhatched | 1 |
| | | | hatched | 1 |

We targeted the V4 hypervariable region of the 16S bacterial ribosomal gene and amplified it with general bacterial primers 515F-Y (GTGYCAGCMGCCGCGGTAA) [158] and 806R (GGACTACNVGGGTWTCTAAT) [159] with the Illumina adapter overhang sequences in both primers. These primers include degenerated bases to be able to amplify Crenarchaeota/Thaumarchaeota and the Alphaproteobacterial clade SAR11. For the PCR, we used the PCRBIO HiFi Polymerase (PCR Biosystems Ltd, UK) and the following conditions: 95˚C for 3 min, followed by 25 cycles of 95˚C for 20 s, 60˚C for 20 s and 72°C for 30 s, with a final elongation step at 72°C for 5 min. We performed DNA amplifications in duplicates, and PCR products were checked in 1% agarose gel and combined. We purified PCR products with AgencourtAMPure XP Beads (Beckman Coulter Inc., USA), and the final libraries were prepared with the Nextera XT DNA Library Preparation Kit (Illumina Inc., USA). All samples were normalized at 4 nM and we generated an equimolar pool of DNA for sequencing. Libraries were run on an Illumina MiSeq device using v3 chemistry (2x300bp) at the Sequencing Facility of the Natural History Museum of London (https://www.nhm.ac.uk/). The resulting amplicon sequence length was ca. 298 bp for the V4 region. Reads were deposited at the Sequence Read Archive (SRA) of the NCBI as BioProject with accession ID PRJNA599541.

The bioinformatic pipeline started with feeding the raw paired reads into Mothur v.1.41.3 and then we followed an adaptation of MiSeq SOP protocol [160]. Briefly, we removed primer sequences and then built sequence contigs using only overlapping paired reads. Sequences with >0 N bases or with >15 homopolymers were discarded. Then we aligned unique sequences to the Silva reference dataset (release 132), and those that aligned poorly, assessed by the Mothur MiSeq SOP pipeline as generating alignments that eliminated too many bases, were removed from the dataset. Denoising of unique aligned sequences was performed with Unoise3 within Mothur [160], allowing 1 difference for every 100 bp of the sequence. Resulting amplicon sequence variants (ASVs[,161)] were checked for singletons and removed at this stage. UCHIME 4.1 was used to check for chimeras with the Silva reference dataset and parameter minh = 1. The taxonomic affiliation of ASVs was obtained using the Silva database v.132, with a cut-off value of 80. The algorithm used for assigning taxonomic identity is that included in the Mothur pipeline (https://mothur.org/wiki/classify.seqs/), using the classify.seqs command. This algorithm is specifically a bayesian naive classifier [162]. We discarded all ASVs classified as eukaryotic, chloroplast or mitochondria. The core microbiome was defined as ASVs that were present in 100% of samples at any abundance. Calculation of alpha diversity (ShannonH index) was done using rarefied counts, and description of the microbial community composition using the total number of ASVs transformed to relative abundances within each individual. Bray-Curtis dissimilarity, calculated with *vegan* package [163], was used in Hierarchical Cluster analysis and Principal Component Analysis ('cmdscale' in *vegan*) for ordination of samples.

In addition, the R package Tax4Fun2 v1.1.5 [164] was used to explore the functional roles of the sponge microbiomes. This package performs similarity searches of the ASVs sequences against annotated genomes on the Kyoto Encyclopedia of Genes and Genomes (KEGG), extracts functional profiles from matching sequences, and creates a predicted metagenome for each sample that incorporates the abundance of the various ASVs. Abundances of the predicted protein orthologues (KO) were used to calculate Bray-Curtis dissimilarity and for PCo Analysis, similarly as before.

### 10.2 Results for microbial community structure within *Ephydatia muelleri*

All samples had between 46,744 to 85,607 counts, showing saturation in rarefaction curves (Supplementary Data 9: 9A), and between 865 to 4,172 unique ASVs. Individual 3 from UK samples had less than 300 ASVs, and was discarded. During the taxonomic annotation, we noticed that between 18.8 and 84.2% of raw counts were annotated as unknown, which was larger than expected. In fact, only one unknown ASVs was dominant in all samples (>97% of the unknown counts). A Blast similarity search of this ASV on NCBI matched with *Ephydatia muelleri* genome assembly, organelle: mitochondrion (100% sequence identity, Genbank: LT158504.1). This ASV, and other unknown ASVs were excluded from further analysis, leaving a filtered dataset with 12,922 to 65,258 raw counts, which showed saturation in rarefaction curves (Supplementary Data 10: 9A), and between 248 to 4,061 unique ASVs.

Microbial taxonomic analysis:

The best BLAST hit for the top 10 most abundant ASVs often retrieved uncultured bacteria from wastewater treatment systems or lake water from Canada and Argentina. Other sources identified were sediment, soil and some bacterial isolates from labs with no further information (Supplementary Data 9: 9B).

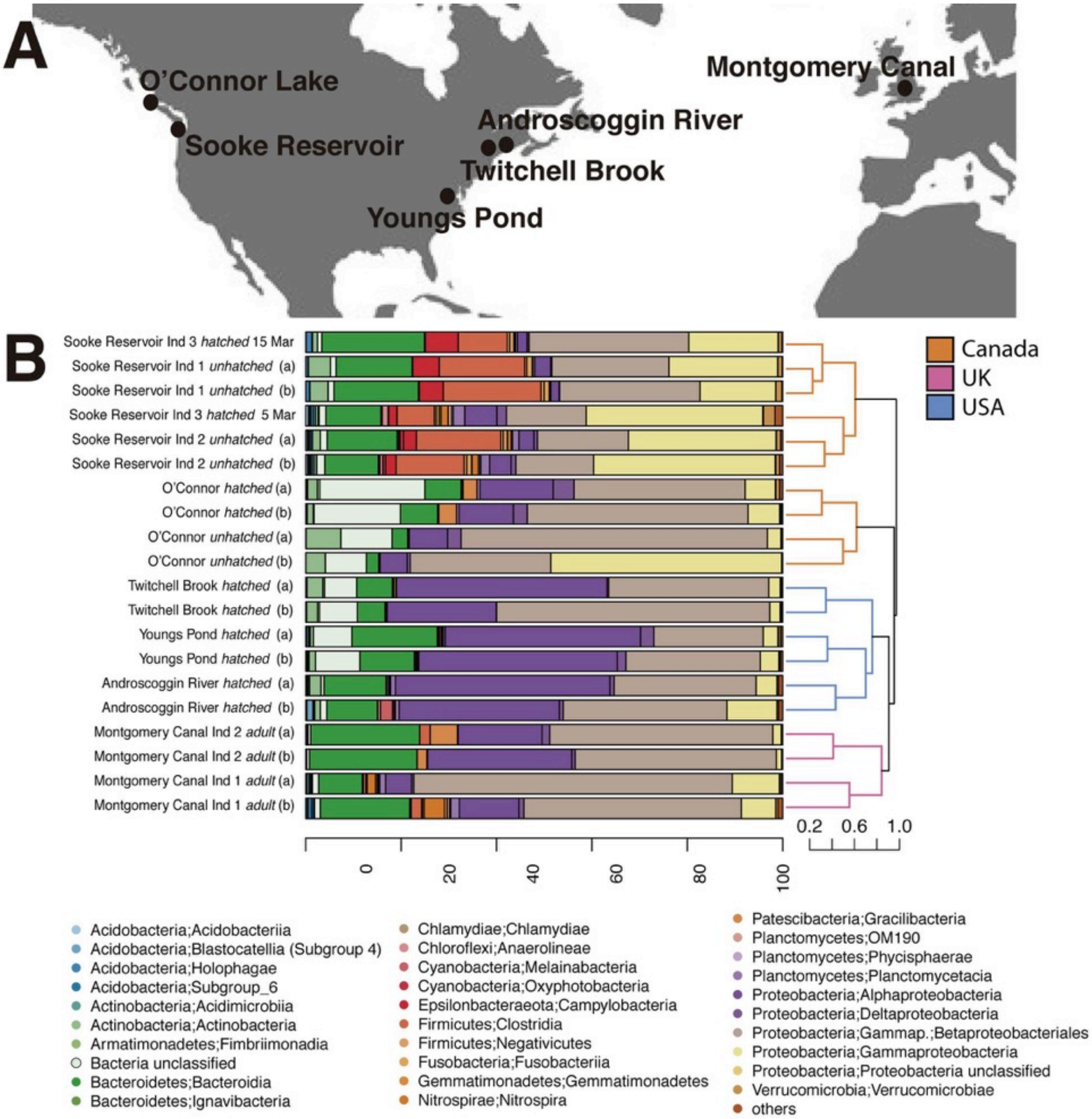

**Supplementary Figure 37.** A) Map showing collection sites. B) Relative abundance of ASVs classified to class level (note that Betaproteobacteriales is an order within Betaproteobacteria). On the right, a dendrogram based on Bray Curtis dissimilarity of all ASVs separates the samples based on their geographic location. Any bacterial class with less than 1% relative abundance was collated under the others category in the plot legend.

Globally, the microbial community was largely dominated by Proteobacteria across all samples (mostly Gammaproteobacteria), followed by Bacteroidia (phylum Bacteroidetes) in most samples, whereas Alphaproteobacteria was particularly abundant in USA samples, and Clostridia (phylum Firmicutes) in

Canadian samples from Sooke Reservoir (Supplementary Table 22 and Supplementary Figure 37B). In general, the microbiome of freshwater sponges is highly abundant on Proteobacteria, especially Alphaproteobacteria and also Betaproteobacteriales [157, 165, 166, 167, 168], which are now within Gammaproteobacteria [39]. This is in part because of the well known differences in pH and nutrient content of freshwater systems, which allow more growth of Alpha and Gammaproteobacteria [169]. In marine sponges, however, other Gammaproteobacteria different than Betaproteobacteriales are more abundant [156]. Planctomycetes, which are taxa abundant in other species of *Ephydatia*, such as *E. fluviatilis* [165], were mostly abundant in hatched gemmules from O'Connor Lake in Canada and adult tissues from the UK (Supplementary Table 22 and Supplementary Figure 37B). The phylum Bacteroidetes is also highly enriched in the tissues of *Ephydatia muelleri* (Supplementary Table 22 and Supplementary Figure 37B), similar to the data previously obtained from the tissues of the tropical freshwater sponge *Tubella variabilis* in comparison to the surrounding waters [157]. Interestingly, only the sponges from Sooke Reservoir seemed to host an abundant pool of Firmicutes within their gemmules (Supplementary Table 22 and Supplementary Figure 37B). Cyanobacteria were moderately abundant (1.29% rel. ab.) in the sponges from Androscoggin river (Supplementary Table 22 and Supplementary Figure 37B), and surprisingly they represented 0.63% rel. ab. (more specifically Oxyphotobacteria) of the total abundance in some unhatched gemmules from Sooke Reservoir even though these sponges were not exposed to light. Archaea were totally absent in half of the sample, and representing an average of only 0.02% rel. ab. in the rest. A number of reads were assigned to uncultured Bacteria mostly among samples recovered from lakes (Supplementary Table 22).

Based on the Bray Curtis dissimilarity of all ASVs, the microbial community was most well-grouped within each sampling site, with samples from the USA more similar to those of the UK than to samples from Canada (Supplementary Figure 37B). A more detailed analysis of the microbial community structure at the species level (top 200 most abundant ASVs), showed that Sooke Reservoir harboured the most different community of all (Supplementary Figure 37A). Indeed, the gemmules from Sooke Reservoir showed large numbers of Gammaproteobacteria (other than Betaproteobacteriales), Clostridia (Firmicutes) and Campylobacteria (Epsilonbacteraeota) within them, which were not very abundant in the rest of samples (Supplementary Table 22 and Supplementary Figure 37B).

**Supplementary Table 22.** Summary of average relative abundance between replicates of ASVs associated to the different taxa. Abbreviations: MC, Montgomery Canal (UK); YP, Youngs Pond (USA); AR, Androscoggin River (USA); TB, Twitchell Brook (USA); OC, O'Connor Lake (Canada); SR, Sooke Reservoir (Canada); u, unhatched; h, hatched.

| | **Average abundance between replicates** | | | | | | | | | |
|---|---|---|---|---|---|---|---|---|---|---|
| **Taxa** | **MC_1** | **MC_2** | **YP_1** | **AR_1** | **TB_1** | **OC_1** | **OC_2** | **SR_1_u** | **SR_2_u** | **SR_3_h** |
| Proteobacteria; Gammaproteobacteria; Betaproteobacteriales | 56.2 | 44.5 | 25.5 | 32.0 | 45.4 | 46.8 | 41.1 | 27.0 | 17.7 | 25.1 |
| Proteobacteria; Gammaproteobacteria | 8.5 | 1.5 | 3.5 | 7.4 | 2.3 | 25.6 | 6.5 | 19.3 | 34.5 | 27.9 |
| Proteobacteria; Alphaproteobacteria | 9.0 | 23.9 | 41.3 | 39.3 | 33.5 | 6.9 | 13.4 | 2.5 | 3.8 | 4.4 |
| Bacteria unclassified | 1.3 | 0.6 | 8.7 | 1.0 | 7.3 | 9.6 | 20.1 | 1.2 | 1.6 | 1.2 |
| Bacteroidetes; Bacteroidia | 14.0 | 22.7 | 14.7 | 11.8 | 6.6 | 2.9 | 7.7 | 16.8 | 13.0 | 16.5 |
| Firmicutes; Clostridia | 1.4 | 1.1 | 0.3 | 0.0 | 0.0 | 0.0 | 0.0 | 19.2 | 16.0 | 9.1 |
| Actinobacteria; Actinobacteria | 0.0 | 0.2 | 1.0 | 1.8 | 2.8 | 5.7 | 1.7 | 4.2 | 1.1 | 0.8 |
| Proteobacteria; Deltaproteobacteria | 0.7 | 1.2 | 2.3 | 0.9 | 0.2 | 1.7 | 3.6 | 0.2 | 0.9 | 1.2 |
| Epsilonbacteraeota; Campylobacteria | 0.0 | 0.0 | 0.0 | 0.0 | 0.0 | 0.0 | 0.0 | 5.3 | 2.4 | 4.3 |
| Patescibacteria; Gracilibacteria | 0.4 | 3.8 | 0.1 | 0.0 | 0.0 | 0.1 | 3.3 | 0.0 | 0.0 | 0.0 |
| Planctomycetes; Planctomycetacia | 1.5 | 0.1 | 0.3 | 0.9 | 0.1 | 0.1 | 0.6 | 0.3 | 1.6 | 1.4 |
| Verrucomicrobia; Verrucomicrobiae | 0.4 | 0.1 | 0.4 | 0.3 | 0.2 | 0.1 | 0.5 | 1.1 | 0.9 | 1.6 |
| Nitrospirae; Nitrospira | 3.0 | 0.0 | 0.0 | 0.1 | 0.0 | 0.0 | 0.0 | 0.1 | 1.1 | 0.8 |
| Fusobacteria; Fusobacteriia | 0.0 | 0.0 | 0.0 | 0.0 | 0.0 | 0.0 | 0.0 | 1.1 | 0.9 | 0.7 |
| Acidobacteria; Holophagae | 0.4 | 0.0 | 0.2 | 0.0 | 0.0 | 0.0 | 0.0 | 0.6 | 0.3 | 0.7 |
| Chloroflexi; Anaerolineae | 0.1 | 0.0 | 0.1 | 0.5 | 0.1 | 0.1 | 0.2 | 0.0 | 0.4 | 0.6 |
| Acidobacteria; Blastocatellia_(Subgroup_4) | 0.4 | 0.0 | 0.3 | 0.7 | 0.1 | 0.0 | 0.1 | 0.0 | 0.2 | 0.1 |
| Firmicutes; Negativicutes | 0.2 | 0.0 | 0.1 | 0.0 | 0.0 | 0.0 | 0.0 | 0.6 | 0.6 | 0.5 |
| Acidobacteria; Acidobacteriia | 0.2 | 0.0 | 0.3 | 0.2 | 0.1 | 0.0 | 0.2 | 0.0 | 0.4 | 0.3 |
| Cyanobacteria; Melainabacteria | 0.0 | 0.0 | 0.0 | 1.3 | 0.0 | 0.0 | 0.0 | 0.0 | 0.0 | 0.0 |
| Actinobacteria; Acidimicrobiia | 0.1 | 0.1 | 0.1 | 0.2 | 0.0 | 0.0 | 0.1 | 0.0 | 0.4 | 0.3 |
| Gemmatimonadetes; Gemmatimonadetes | 0.2 | 0.0 | 0.1 | 0.2 | 0.3 | 0.0 | 0.1 | 0.0 | 0.1 | 0.2 |
| Acidobacteria; Subgroup_6 | 0.4 | 0.0 | 0.0 | 0.2 | 0.0 | 0.0 | 0.0 | 0.0 | 0.3 | 0.3 |
| Planctomycetes; OM190 | 0.4 | 0.0 | 0.0 | 0.0 | 0.0 | 0.0 | 0.0 | 0.0 | 0.3 | 0.4 |
| Spirochaetes; Leptospirae | 0.0 | 0.0 | 0.0 | 0.0 | 0.0 | 0.1 | 0.4 | 0.0 | 0.0 | 0.0 |
| Armatimonadetes; Fimbriimonadia | 0.0 | 0.0 | 0.0 | 0.0 | 0.4 | 0.0 | 0.3 | 0.0 | 0.0 | 0.1 |
| Proteobacteria; Proteobacteria unclassified | 0.1 | 0.0 | 0.1 | 0.0 | 0.1 | 0.1 | 0.1 | 0.0 | 0.1 | 0.1 |
| Cyanobacteria; Oxyphotobacteria | 0.0 | 0.0 | 0.0 | 0.0 | 0.0 | 0.0 | 0.0 | 0.1 | 0.6 | 0.1 |
| Planctomycetes; Phycisphaerae | 0.3 | 0.0 | 0.0 | 0.1 | 0.0 | 0.0 | 0.0 | 0.0 | 0.1 | 0.2 |
| Bacteroidetes; Ignavibacteria | 0.0 | 0.0 | 0.1 | 0.1 | 0.0 | 0.0 | 0.0 | 0.0 | 0.1 | 0.2 |
| Chlamydiae; Chlamydiae | 0.1 | 0.0 | 0.0 | 0.0 | 0.1 | 0.0 | 0.0 | 0.0 | 0.3 | 0.0 |
| Others | 0.6 | 0.1 | 0.4 | 0.8 | 0.2 | 0.1 | 0.1 | 0.1 | 0.5 | 0.9 |

Interestingly, even though the entire genome of a *Flavobacterium* sp. was recovered from the assembled genome of *E. muelleri* of Sooke Reservoir (S4), Flavobacteriales were only abundant in the UK sponges (i.e. 9.8 and 15.7% rel. ab.). In fact, all *Flavobacterium* sequences accounted for only 2.1 % of average rel. ab. among samples of Sooke Reservoir. The 16S rRNA sequence retrieved from the *Flavobacterium* sp. genome was identical to one of the ASVs that was abundant in UK ind. 2 sample (13% rel. ab.), but absent in Sooke Reservoir samples. The most abundant ASVs (among genus Flavobacterium) in Sooke samples accounted for only 0.71% average rel. ab., and this ASVs had a sequence similarity of 98.8% to the *Flavobacterium* sp. ribosomal gene. It is unclear how the genome of a *Flavobacterium* sp. (low abundant bacterial genera in the sponge community) was sequenced instead of the most abundant bacterial member in the Sooke Reservoir samples (i.e. genus Methyloglobulus, Gammaproteobacteria).

<u>Alpha and Beta diversity</u>: Most of the samples presented an alpha diversity of 2 to 3 (Shannon index). Samples from Sooke Reservoir (Canada), however, were the most diverse of all, while Montgomery Canal samples (UK) had the lowest diversity (Supplementary Figure 38A).

Ordination of the samples, based on the Bray Curtis dissimilarity matrix, indicated that Youngs Pond and Androscoggin River from USA, were very similar in their microbial community composition and different to Twitchell Brook. Sooke Reservoir and O'Connor lake from Canada, however, harbored more distant microbial communities. Moreover, samples from Canada included unhatched and hatched gemmules in both locations, which overlapped in the ordination plot suggesting small differences among them. The sponge tissue from the two adult individuals in Montgomery Canal (UK) presented a more variable communities among them than any gemmule samples coming from different individuals (Supplementary Figure 38B). Interestingly, grouping of samples by the predicted functions followed a different distribution. Young Pond samples (hatched) were closer to ind. 1 (hatched) and ind. 3 (unhatched) of Sooke Reservoir and O'Connor samples (hatched), while ind. 2 (unhatched) was distant from this group. Moreover, O'Connor hatched and unhatched samples were highly similar based on the microbiome composition, but they seemed to include rather different bacterial functions (Supplementary Figure 38B).

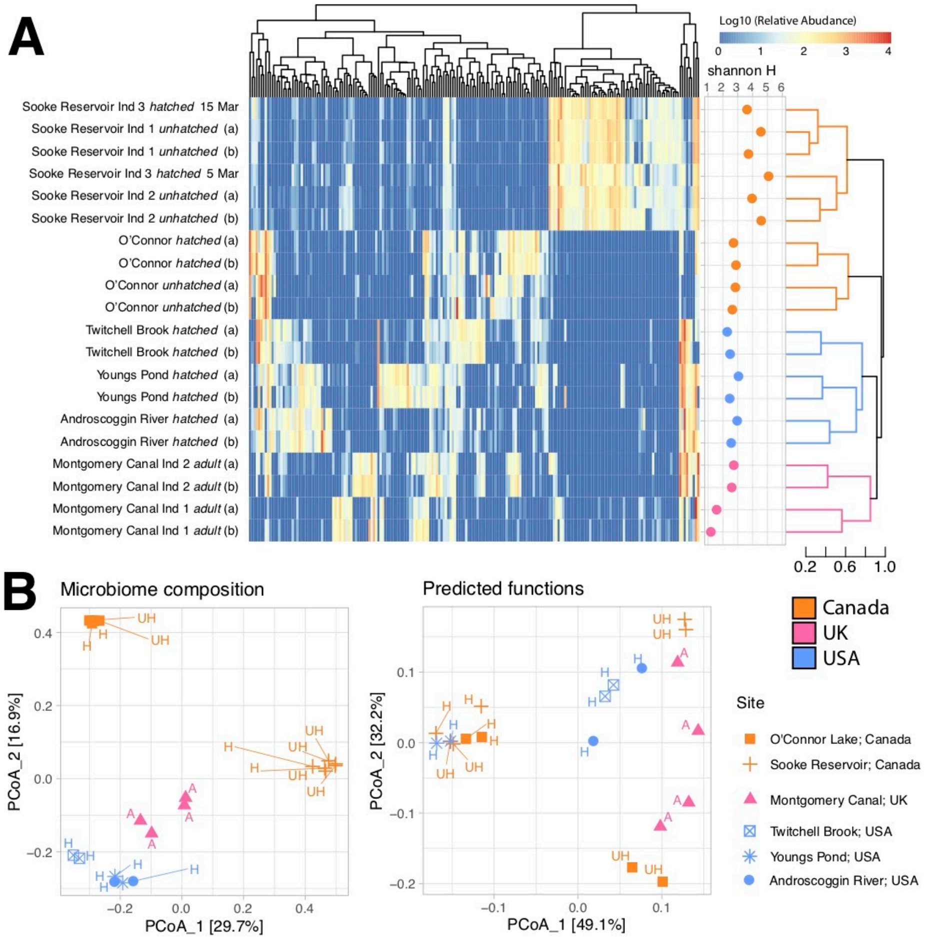

**Supplementary Figure 38.** A) Heatmap of the most abundant ASVs across samples and alpha diversity (right). B) Principal component analysis based on the microbiome composition (left) and predicted functions (right) of all samples. Abbreviations: A, adult; H, hatched gemmules; UH, unhatched gemmules.

Core community: Surprisingly, despite the 6,500 km separating the furthest collection sites, there were 4 ASVs present in all *E. muelleri* samples (Supplementary Figure 39). These were two Burkholderiaceae (Gammaproteobacteria), one Ferruginibacter (Chitinophagales) and one unclassified Bacteria. The percentage these core ASVs represented in their samples was rather variable, from < 1% in Sooke Reservoir samples, to > 20 % in Twitchell Brook or Montgomery Canal samples. The most abundant Burkholderiaceae ASV, had a best blast hit of 100% identity to MH776330.1 which was a clone from a study on lab-scale wastewater treatment with artificial wastewater in Japan (https://www.ncbi.nlm.nih.gov/nuccore/MH776330.1/). Considering a core microbiome of 80% of the samples, the number of shared ASVs increased to 17, representing from 6 to 59% relative abundance of the samples. These included another seven Burkholderiaceae ASVs, two Rhodobacteraceae, one Devosiaceae, one Bradyrhizobium, one Methylophilaceae and one unclassified Bacteria. The number and abundance (avg. 83% of the core abundance) of ASVs assigned to genera Burkholderiaceae among the core microbiome points to an important role of this group within the microbial community of *E. muelleri*.

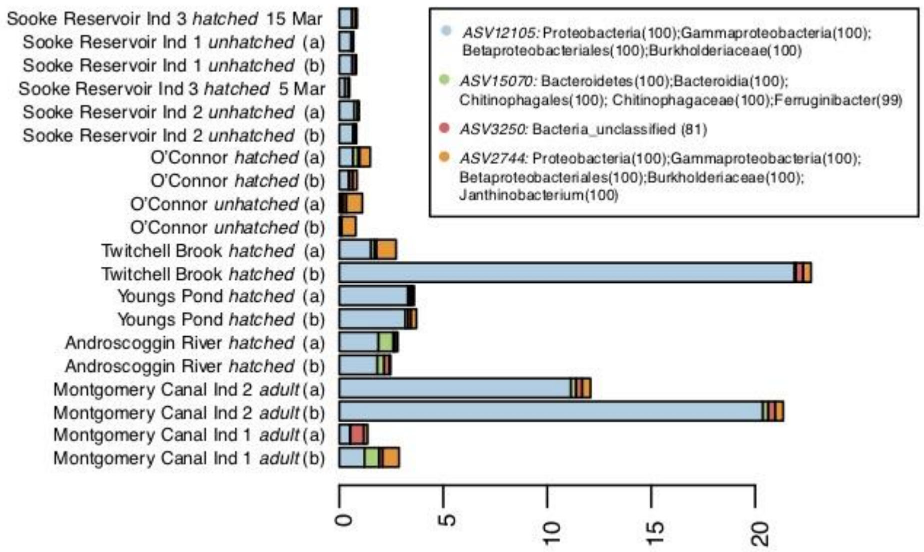

**Supplementary Figure 39.** Relative abundance of the four ASVs shared by all studied samples. Taxonomic classification by Silva v. 132 is shown in the figure legend with confidence values in brackets.

We also investigated the number of ASVs shared between any pair of samples. Sooke Reservoir samples had the largest core community of all locations (610 – 1,160 core ASVs). O'Connor Lake samples, also from Canada, had a much smaller core community of 246 – 501 core ASVs, while USA samples shared 184 – 616 core ASVs, and UK samples 115 – 346 core ASVs (Supplementary Figure 40). Interestingly, in other freshwater sponges, up to 1,305 OTUs were shared among individuals of the same species and collection site (*Tubella variabilis*; [157]), but only less than 200 between sponges of different species [167]. In this sense, our study sheds light on the strong effect of the geographic location over the microbiome of several specimens of the same species collected in a large scale study, which results in very few ASVs shared among different distant locations, while large numbers within specimens of the same location (especially among the Sooke Reservoir samples).

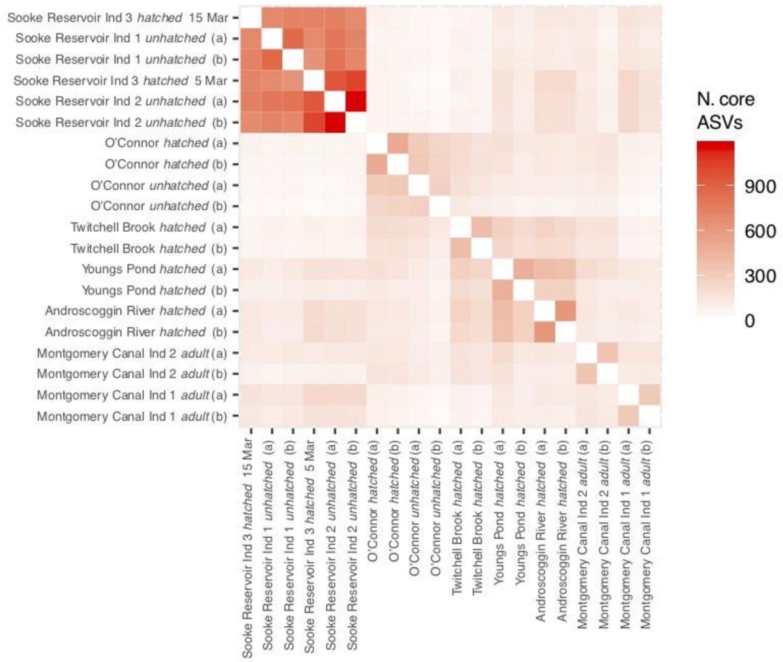

**Supplementary Figure 40.** Number of shared ASVs between every pair of samples.

The % of the sequences UNUSED in the prediction ranged from 0.14% to 0.83%. Specific values are:

*Montgomery Canal:* Ind 1 adult (a) 0.28743, Ind 1 adult (b) 0.45442, Ind 2 adult (a) 0.33108, Ind 2 adult (b) 0.28994, Ind 3 adult (a) 0.80292, Ind 3 adult (b) 0.8281

*Youngs Pond:* hatched (a) 0.33143, hatched (b) 0.26188

*Sooke Reservoir:* Ind 1 unhatched (a) 0.59985, Ind 1 unhatched (b) 0.55047

*Androscoggin River:* hatched (a) 0.23593, hatched (b) 0.24604

*Twitchell Brook:* hatched (a) 0.18642, hatched (b) 0.16903

*O'Connor:* unhatched (a) 0.23265, unhatched (b) 0.14755

*O'Connor:* hatched (a) 0.44069, hatched (b) 0.37871

*Sooke Reservoir:* Ind 2 unhatched (a) 0.71708, Ind 2 unhatched (b) 0.75985

*Sooke Reservoir:* Ind 3 hatched 5 Mar 0.71608, Ind 3 hatched 15 Mar 0.5216

By way of comparison, unused values in the example Tax4Fun dataset are always above 0.90%.

### Supplementary Note 11: *Ephydatia* as a research model

*Ephydatia as an emerging model system:*

The cosmopolitan nature of this freshwater sponge species makes it an ideal choice for a model system. The ease of collection, storage, and use in laboratory work of *Ephydatia muelleri*, especially when compared to the current demosponge model *Amphimedon queenslandica,* which can only be collected by researchers in one location worldwide, and cannot be cultured easily in the laboratory, further contributes to its practicality as a model system. Assets of this system include:

1. The gemmules can be frozen in liquid nitrogen and stored below -80°C and hatched up to several years later without loss of viability [19, 170].
2. *E. muelleri* gemmules are clones. Each individual sponge contains thousands of gemmules enabling studies to use many highly uniform replicates. Each clone is either male or female [171, 172].
3. The cost of collecting, storing, and culturing *E. muelleri* is extremely low compared to many other animal-models. Gemmules are collected by hand in rivers and lakes in winter months. Gemmules are stored at 3-4°C in fridges or incubators in freshwater for months. Gemmules hatch in sterile freshwater media in Petri dishes (Supplementary Figures 41, 42) and develop into juvenile sponges on the bench-top [19].
4. Living tissues of freshwater sponges are transparent allowing study of cell activity *in vivo* (e.g.[173, 174]).
5. Methods for preservation of tissues for light, scanning and transmission electron microscopy, and in situ hybridization are well-established [133, 175, 176].
6. RNA interference [177] and DNA methylation [124] methods have been demonstrated.
7. The karyotype of *E. muelleri* is known [24]. The mitochondrial genome is known [178, 179].

There is a rich history of research on this species. Topics cover the distribution, tolerance to cold, pH, temperature and environmental pollutants, silica production, development, physiology and behaviour. Genomic, transcriptomic, and other genetic resources will expand possibilities for primary research as well as projects for educational and citizen-science initiatives.

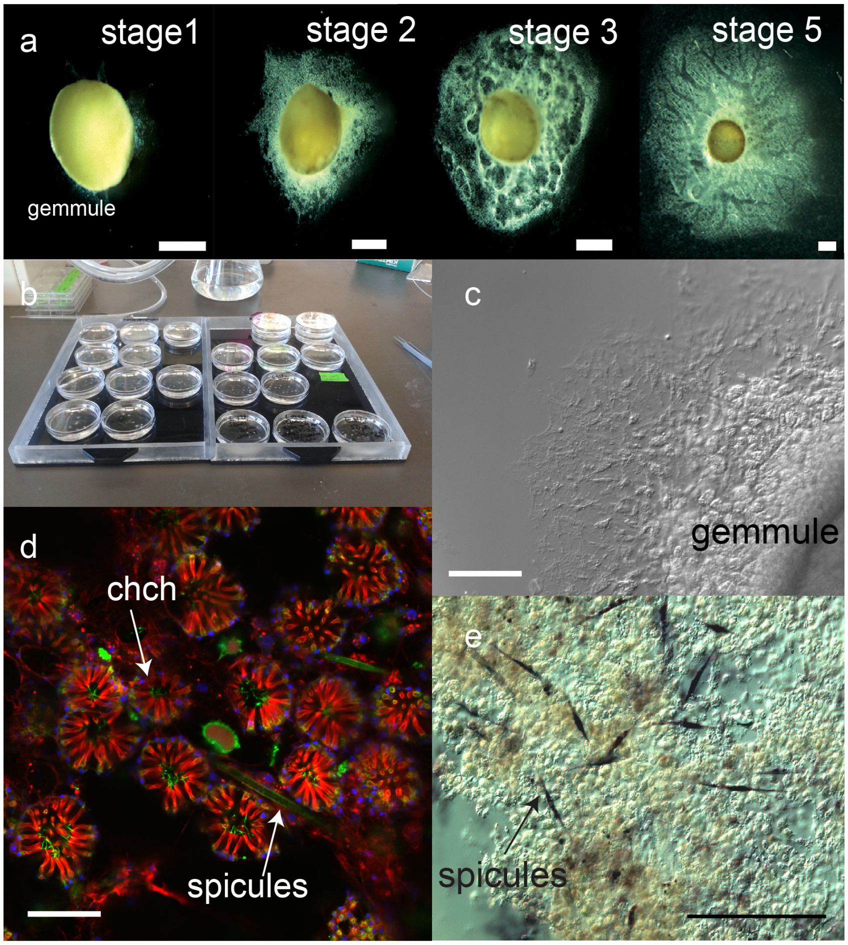

**Supplementary Figure 41.** A) Developmental stages of *Ephydatia muelleri* grown in the laboratory from cleaned gemmules: Stage 1, cells starting to emerge from the micropyle and anchor the gemmule to the substrate; Stage 2, a full epithelium has formed, covering cells that are undergoing differentiation into choanocytes, sclerocytes and pinacocytes; Stage 3, regionalization of the sponge by formation of epithelial-lined lacunae to which choanocyte chambers are attached; Stage 5, a fully organized sponge with ostia, canals, chambers and osculum. B) Gemmules from an individual are clones and multiple sponges can be grown for experiments using a single clone or different clones. C) Developing sponges are transparent to light microscopy and cell behaviour during development can be observed by differential interference microscopy. D) Sponges attached to coverslips are easily preserved for fluorescence microscopy. Choanocyte chambers (chch) and spicules are easily identified by the actin (red) labelling microvilli of collars, and tubulin (green) labelling flagella in cc and microtubules around spicules. E) In situ hybridization using silicatein as a control identifies expression in cells called sclerocytes that form spicules (Image: Pamela Windsor-Reid). Scales: a, 0.5mm; c, 100µm; d, 30 µm; e, 60µm. In these images only one replicate is shown, but in all cases the number of individual choanocyte chambers, spicules or animals that could be observed with these patterns is >100.

<u>*Description of the stages of development shown in Supplementary Figure 41.*</u>

*Stage 1:* approximately in the first days after 'plating' a sponge gemmule in culture medium.

The gemmule contains ‘thesocytes’, which are storage cells full of glycogen inclusions. As the gemmule warms from 3°C storage temperature to room temperature, thesocytes begin to differentiate. In other species (e.g. *Spongilla lacustris*, Saller, 1988 [180]) this has been shown to involve duplication of nuclei, typically forming four nuclei, followed by division of the cell into four cells, which are now called amoebocytes. Differentiation begins at the site of the micropyle, a vase-like opening at one side of the spherical gemmule. The micropyle has a collagenous cover, which breaks down, and amoebocytes emerge.

The first amoebocytes to emerge must develop into rudimentary pinacocytes because no stage is visible that lacks an external layer of cells covering subsequent emigration of amoebocytes. Thus stage 1 sponges have both thesocytes undergoing differentiation and division into amoebocytes, breaking down glycogen reserves, amoebocytes differentiating into early pinacocytes (which lack the stable plate-like form of later pinacocytes and presumably lack fixed cell-cell junctions at this early stage).

*Stage 2:* between 1 and 3 days post hatching.

At this point all cells have emerged from the gemmule husk, which now lies empty and forms a substrate over which the cells form the new sponge. Stage 2 sponges possess a complete exo-pinacoderm that covers a mass of opaque cells consisting of amoebocytes, many still with glycogen inclusions, but also sclerocytes with developing spicules, as well as choanocyte chambers either lying ‘loose’ among the amoebocytes, or attached to rudimentary epithelial-lined pockets. Most chambers are in the tissue adjacent to the gemmule husk. Some ostia are present on the exopinacoderm.

*Stage 3:* between 2 and 4 days post hatching

Stage 3 sponges are most clearly identified by large clear lacunae (pockets) that take up most of the sponge tissue surrounding the gemmule husk. The lacunae are the first steps in excurrent canal formation and vary in size and shape, merging and dividing over time. At this stage choanocyte chambers are attached to lacunae in tidy rows, and yet others are forming and yet to be attached to canals. The pinacoderm is complete, but no subdermal cavity has yet formed, and there is no osculum. Spicules have formed and are being positioned around the sponge, as has been described previously for a sister species *Ephydatia fluviatilis* by [174].

*Stage 4:* between 3-5 days post hatching

Lacunae are now formed into canals which snake around the sponge without pattern. Choanocyte chambers are dense along canals and begin to take on a three-dimensional structure, although most canals are still only in the plane of the dish on which the sponge is growing. Importantly by stage 4 an osculum has formed, often at a more peripheral location away from the gemmule husk.

*Stage 5*: 4-7 days post hatching, and a terminal stage for lab grown sponges that are not fed and cultured for longer. Sponges one week and older are typically called ‘stage 5’

At stage 5 the canals have merged further to form either just one or more typically two large excurrent canals. Often these straddle the gemmule husk. The canals are larger at the base of the osculum and the osculum has developed a wider base and appears longer than in stage 4 sponges. Chambers, spicules and the subdermal space (which is an area lying below the three-layered ‘tent’-like structure that forms the outer tissue of the sponge) are fully formed.

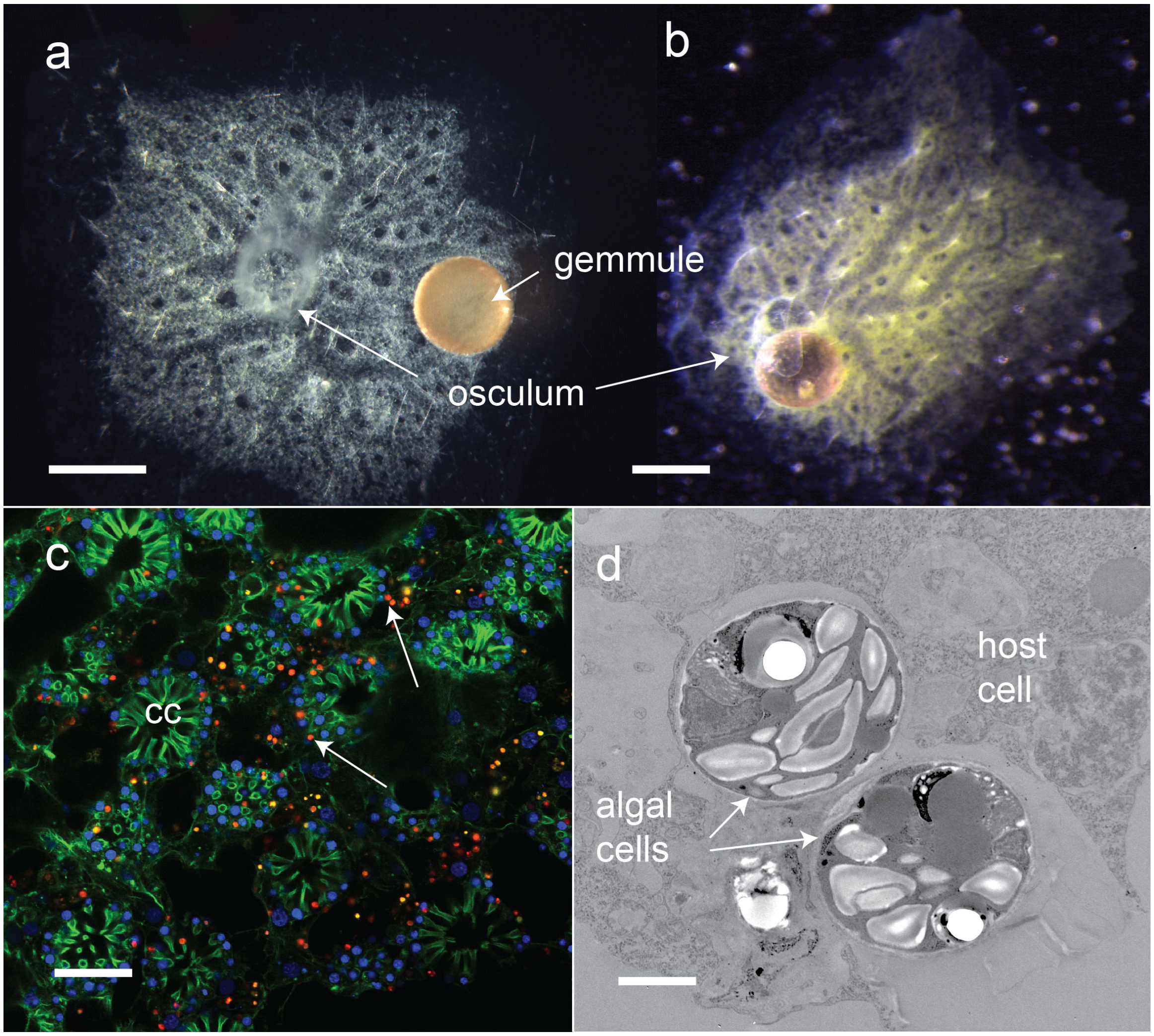

**Supplementary Figure 42.** *Ephydatia muelleri* is an excellent model for studies of symbioses and can be grown in the lab without (A) or with (B) algal symbionts. Aposymbiotic cultures can be infected with algal symbionts. C) Symbionts are readily identified by autofluorescence (arrows, symbionts red). D) Algal symbionts are taken up by host cells. Microbial symbionts are also variable in different populations of *E.*

*muelleri* (further information in Supplementary Note 10) and can be identified at both the light and electron microscope levels. Scales: a, b, 1mm; c, 30 µm; d, 2µm.

*<u>Ephydatia muelleri</u> as a model for intracellular symbiosis*

*E. muelleri* is also an emerging model for the study of intracellular endosymbiosis. Symbioses are ubiquitous features of all aquatic ecosystems having effects that extend beyond the partners involved in the association. Indeed, the origin of the eukaryotic cell occurred in an aquatic environment as the result of symbiotic integration[181]. Thus, understanding ecological and evolutionary forces that shape intracellular symbioses as well as the cellular and genetic factors that contribute to the integration of host and symbiont is an important goal. *E. muelleri* can form associations with microalgae resulting in intracellular symbioses between a phototrophic symbiont and heterotrophic host (Supplementary Figure 42 C,D). The ecological importance of photosynthetic sponges in freshwater ecosystems has been documented for decades (e.g., [182,183,184, 185, 186, 187, 188]), and now with genome resources, *E. muelleri* provides a tractable model system to study mechanisms and cellular pathways leading to long-term intracellular relationships. *E. muelleri* can be cultured in the lab with and without algal symbionts and the infection and intracellular occupancy can be monitored by microscopy as well as interrogated with molecular and genetic tools.
